# Emergent feedforward and isohydric responses to soil and atmospheric aridity: Insights from a time-dependent hydraulic model

**DOI:** 10.1101/2025.07.04.663235

**Authors:** Fulton E. Rockwell

## Abstract

Drought and heat waves have synergistic effects on mortality as plants experience both supply and demand-side water stress. Which of these stresses ultimately controls stomatal conductance (*g_s_*) and whether patterns of regulation represent biological strategies or are imposed on the plant are addressed by way of a mechanistic hydraulic ‘null’ model.

- Dynamic losses of soil hydraulic conductivity are modeled in a soil domain fed by a maximum water flux from a deep-water source. A root-uptake plane extracts water from the soil in a two-node (root and leaf) plant model, with *g_s_* a function of leaf water potential.
- Multi-day simulations reproduce previously published experimental observations of anisohydric to isohydric transitions, driven by the balance of soil supply and atmospheric demand. Feedforward control of transpiration *E* emerges from steep declines in soil hydraulic diffusivity that confine diurnal variation in moisture gradients and water discharge/recharge cycles to a shallow region at the root plane that thins with increased demand.
- Empirical models of *VPD* sensitivity are compared to the full model to provide mechanistic insight into empirical parameters. Apparent responses of *g_s_* to *VPD* are shown to emerge from plant, soil and atmospheric feedbacks that are both time and scale dependent.

## Introduction

The dual role of stomata in admitting an inward flux of *CO_2_*while preventing the desiccation of plant tissues has been recognized at least since Van Den Honert (1948), who proposed that stomata serve as the dominant resistance to water transport in the soil-plant-atmosphere continuum (‘SPAC’; Philip, 1966). While guard cells, as hydraulicly actuated valves, respond ‘passively’ to local water status, pore aperture is also decoupled from the local water potential of the epidermis by ‘active’ biochemistry (e.g., light activated ion pumping, ABA induced closure; McAdam & Brodribb, 2016; Jezek & Blatt, 2017), with the resulting complexity of regulation obscuring the control points that sense water status and govern stomatal behavior (Tardieu & Simonneau, 1998; Buckley, 2005; McAdam & Brodribb, 2016; Novick *et al*., 2019; Jezek *et al*., 2019; Grossiord *et al*., 2020; Wankmüller & Carminati, 2022). While the twin stressors of heat and drought are known to have synergistic mortality effects (Mitchell *et al*., 2014; Matusick *et al*., 2018), the lack of a clear understanding of how stomata integrate supply and demand stresses has led to controversy regarding plant responses to future climates, a debate focused on whether stomatal conductance is more limited by available soil water or atmospheric aridity (Roderick *et al*., 2015; Novick *et al*., 2016a, 2024; Berg & Sheffield, 2018; Grossiord *et al*., 2020; Liu *et al*., 2020a; McColl & Tang, 2024).

The biologically ‘active’ nature of stomatal regulation suggests that the 60% of global terrestrial evapotranspiration mediated by plant canopies is subject to a diversity of stomatal control strategies varying across species, functional groups and biomes (Tardieu & Simonneau, 1998; Oren *et al*., 1999; Martinez-Vilalta & Garcia-Forner, 2017; Novick *et al*., 2019; Wu *et al*., 2021). Yet, caught between Equivalent Water Thicknesses (EWT) for both the unsaturated soil zone and atmosphere on the order of centimeters, the plant canopy with an EWT on the order of millimeters may have little scope to express independent control of its water status at equilibrium (Shiklomanov, 1993; Hunt, Jr. *et al*., 2009). The behavior of the plant canopy may then follow that of any ‘passive’ material that loses conductivity as it dries (Hochberg *et al*., 2018; Feng *et al*., 2019; Vargas Zeppetello *et al*., 2023). With respect to the global water cycle, the plant canopy may approximate ‘green dirt.’

Physiologically, interest in vapor pressure deficit (*VPD*) responses as distinct from source water limitations accelerated with evidence of ‘feed-forward’ stomatal behavior: stomatal closure that does not merely reduce increases in transpiration *E* with increases in atmospheric demand, but results in *lower E* (Cowan & Troughton, 1971). Such behavior cannot be explained by local hydraulic resistances that create steady-state feed-backs of *E* on guard cell water potential and stomatal apertures. Local feed-backs describe an incremental draw-down in water potential resulting from an incremental *increase* in *E*, thereby reducing the realized increase in *E* but not *E* itself (Cowan & Troughton, 1971; Franks *et al*., 1997,1999; Buckley, 2005). The inadequacy of local hydraulic feedbacks, together with demonstrations that leaf water potential and stomatal conductance do not follow a one-to-one relationship, has turned attention toward metabolic and chemical controls on stomatal responses to water stress (Turner *et al*., 1984, 1985; Gollan *et al*., 1985; Simonneau & Tardieu, 1998; McAdam & Brodribb, 2016).

This change of research focus makes sense: the principal activity of guard cell physiology is to break any one-to-one relationship of stomatal pore aperture to water potential by manipulating ion concentrations in response to an array of signals and forcings, including ABA, photon flux density, light quality, substrate availability, enzyme activation, *CO_2_* concentrations and temperature (Schroeder *et al*., 2001; Messinger *et al*., 2006; Brodribb & McAdam, 2011; Buckley, 2019; Jezek *et al*., 2019). Despite, or perhaps because of the richness of physiological detail that has been described at the guard cell level, efforts to predict whole plant responses to environmental forcing at scale have adopted empirical approaches that produce behavior consistent with physiological response patterns but do not require mechanistic closure (Anderegg *et al*., 2017; Mrad *et al*., 2019; Grossiord *et al*., 2020; Wang *et al*., 2020; Mencuccini *et al*., 2024).

The most widely adopted empirical approach to partitioning changes in stomatal aperture between supply and demand limitation has been to model maximum stomatal conductance (*g_s_*,_ref_) as a function of soil moisture, and fit a slope parameter (*m*) that describes the erosion of conductance with increasing vapor pressure deficit (*VPD*) (Oren *et al*., 1999; Novick *et al*., 2016a, 2019; Grossiord *et al*., 2020). This approach has attracted the criticism that soil moisture and *VPD* are not independent at the spatial and temporal scales relevant to climate (Roderick *et al*., 2015; Novick *et al*., 2016a; Berg & Sheffield, 2018; Liu *et al*., 2020a). Even at the sub-daily physiological scale of flux tower data over which bulk soil moisture and *VPD* are more weakly correlated (Novick *et al.,* 2024) it is unclear to what extent apparent stomatal responses to *VPD* are truly independent of soil moisture stress (Novick *et al*., 2019; Koehler *et al*., 2024), and so whether parameterizing independent soil and *VPD* responses amounts to double counting (Liu *et al*., 2020b).

At daily time scales, an empirical approach has been developed that predicts *E* and surface conductance solely from weather data. The ETRHEQ hypothesis (EvapoTranspiration from Relative Humidity at Equilibrium) minimizes the vertical variance of RH through the Atmospheric Boundary Layer (ABL) to find the surface conductance (interpreted as *g_s_*; Grossiord *et al.,* 2020). ETRHEQ can then be used to construct a response of *g_s_* to *VPD* (Salvucci & Gentine, 2013; Rigden & Salvucci, 2015). As an empirical steady-state theory, ETRHEQ does not specify the mechanisms driving the relationships it describes. To address this gap, McColl *et al*. (2019) and McColl & Rigden (2020) provided a simple theory of Surface Flux Equilibrium (SFE) that relates the RH of the ABL to the balance of latent and sensible heat at the surface. As the system evolves toward an ‘equilibrium’ steady-state at which inputs from the surface match the fluxes out of the top of the ABL, the state of the land surface becomes fully reflected in the state of the atmosphere over time scales ranging from daily for ‘wet’ surfaces, to somewhat longer for ‘dry’ surfaces (McColl & Rigden, 2020).

SFE does not necessarily offer a unique explanation for ETRHEQ, nor can it establish the direction of causality between RH and *g_s_*based on their internal relationship in the stationary-state. What SFE does do is capture the ETRHEQ hypothesis in a model that includes feedbacks on *E* due to humidification and heating of the atmosphere by the surface heat and moisture fluxes, without imposing a prescribed biological response of *g_s_* to *VPD*. The success of SFE/ETRHEQ begs the question of whether surface conductance is determined by active biological optimization and adaptive strategies for stress avoidance, or is primarily physically constrained (Konings & Gentine, 2017; Wu *et al.,* 2021; Vargas Zeppetello *et al.,* 2023; Mencuccini *et al.,* 2024), a gap in understanding that has led to calls for further work on the problem (Novick et al., 2019; Grossiord et al., 2020). There is also continued related uncertainty as to whether the isohydric-anisohydric spectrum of behavior describes biological adaptation, or reflects physical limits imposed on the plant by soil and atmospheric aridity (Konings & Gentine, 2017; Martínez-Vilalta & Garcia-Forner, 2017; Hochberg *et al*., 2018; Feng *et al*., 2019; Novick *et al*., 2019; Wu *et al*., 2021; Mencuccini *et al*., 2024).

One element often missing from these debates is an appropriate ‘null model’ for environmental control of stomatal conductance that might be employed to separate biological versus purely physical regulation of *E* (Hochberg *et al.,* 2018; Carminati & Javaux, 2020; Rodriguez-Dominguez & Brodribb, 2020; Vargas Zeppetello *et al.,* 2023; Manandhar *et al.,* 2024; Mencuccini *et al.,* 2024). Toward this end, this study develops a time-dependent SPAC model from a continuum approach to soil water transport coupled to an ohmic (discrete) plant with stomatal conductance responding solely to leaf water potential. The goal is not to build an exhaustive mechanistic model of water and *CO_2_* fluxes in the SPAC (Tuzet *et al*., 2003), nor to provide a sub-model appropriate for incorporation into large-scale land surface and atmosphere models (Sloan *et al*., 2021). Rather the aim is to construct a minimal hydraulic representation of the SPAC for understanding environmental contributions to experimentally observed patterns of transpiration. Such a model may help resolve the physical factors contributing to the fitted parameters in scalable empirical models of stomatal responses to *VPD* and soil drought (Attia *et al*., 2015; Konings & Gentine, 2017; Novick *et al*., 2019; Mrad *et al*., 2019; Grossiord *et al*., 2020; Mencuccini *et al*., 2024).

The null model developed here is first employed to test whether a minimal hydraulic time-dependent approach can successfully recover observations of 1) intra-day feed-forward stomatal behavior in response to diurnally varying *VPD*, 2) conservation of total daily transpiration across days and *VPD* levels, and 3) isohydric midday leaf water potentials in response to a doubling of *VPD,* as previously observed in a whole-tree chamber experiment (Drake *et al.,* 2018). As in leaf-level studies of VPD responses using gas exchange cuvettes that minimize boundary layer effects (Franks *et al.,* 1997; Mott & Parkhurst, 1991), whole tree chambers allow the isolation of stomatal responses to stress *in situ* and at the whole plant level. The model is then employed to explore the effectiveness of a hypothetical osmotic adjustment to maintain carbon gain, and a sensitivity analysis is undertaken to understand the basis for the emergence of feed-forward and aniso/isohydric behavior from soil water dynamics. Finally, two empirical steady-state models are compared for their ability to capture the patterns of stomatal sensitivity to *VPD* that emerge over physiological (hourly) versus diel (daily) time scales, and these are then compared to the patterns expected under SFE over multiday time scales.

## Model Description

### Model structure and analysis of basic behavior

In SPAC models the three-dimensional transfer of water from soil to roots and from leaves to the atmosphere is collapsed into a single dimension. Here, time-dependent flow in the soil is modeled in cartesian rather than cylindrical coordinates (See Supplemental Information Notes **S1** for a full discussion and derivation). The extra ‘resistance’ created by the concentration of the flux as it approaches the root surface is neglected, which for the purposes of demonstrating soil constraints on transpiration is a conservative choice. The dynamics of infiltration by rainfall into the soil domain are not modeled; in principle meteoric inputs re-set the initial condition for soil moisture with which the model is run. As in general the soil is wetter below the root zone as plants extract water during meteoric drought (Miguez-Macho & Fan, 2021), the model domain is fed at the lower boundary by an asymptotic (maximum) soil flux determined by soil properties and the effective depth *L* (m) to a deep source of soil moisture (Gardner, 1958; Gardner & Fireman, 1958; Jury *et al*., 1991). A rooting surface located at a characteristic depth bounds the top of the model domain (Supplemental Information Notes S1; Binks *et al*., 2022), serving as a sink driven by transpiration through a discrete plant composed of two nodes (root and leaf water potential) separated by a whole plant conductance (Fig. **1a**). The neglect of plant capacitance follows Cowan’s (1965) arguments that soil capacitance is expected to dominate the time scale of storage effects. Plant hydraulic conductance *K_plant_* is held constant based on the hypothesis that stomatal closure precedes cavitation (Hochberg *et al.,* 2017), while varying tissue hydraulic conductivity lies outside the scope of a minimal physical model.

**Figure 1.**
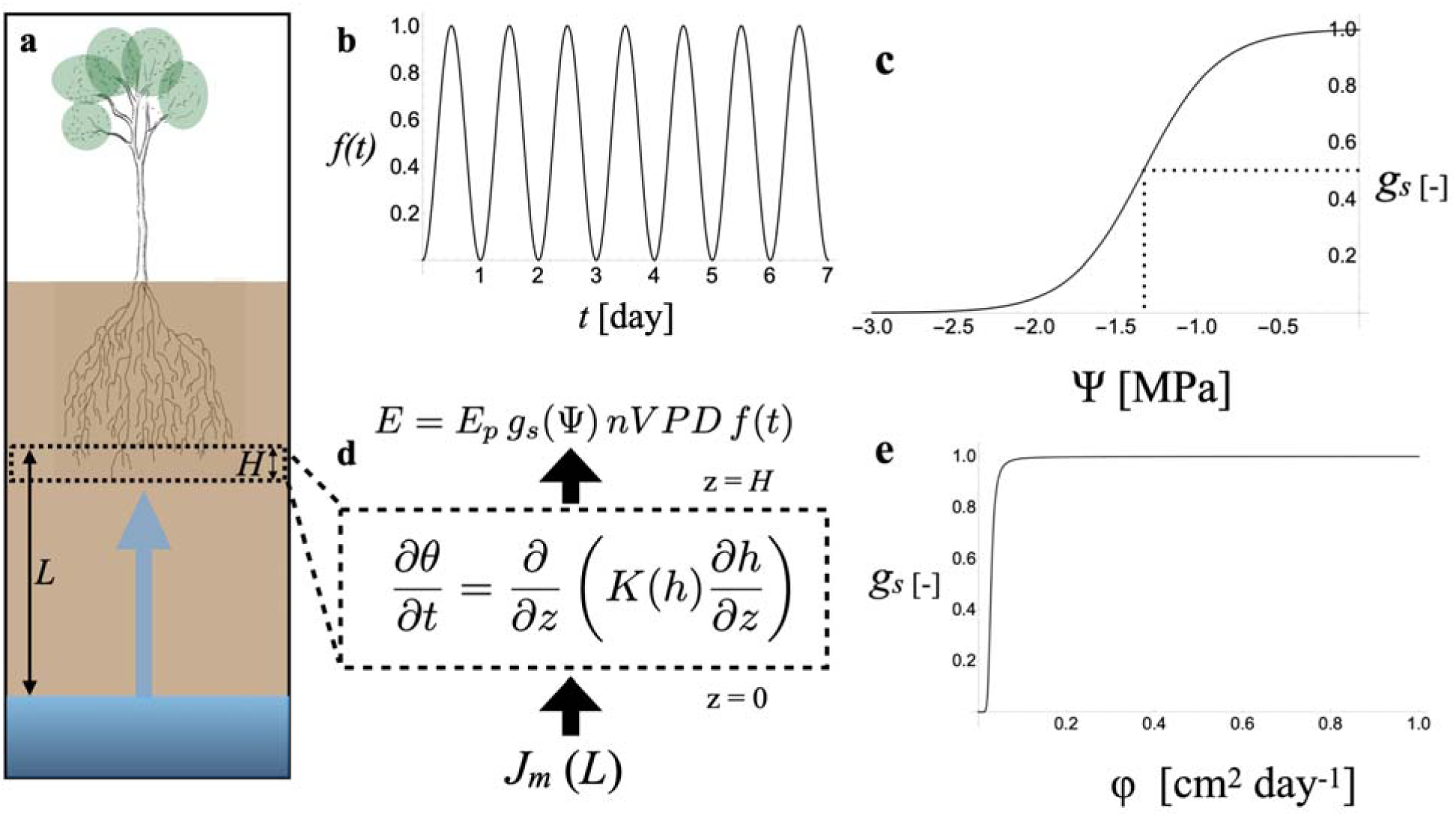
Minimal hydraulic model of the soil-plant-atmosphere continuum. **a** The last ten centimeters of the distance *L* from the water table to rooting depth forms the modeled soil domain, bounded at the lower surface by a maximum flux from the water table *J_m_*, and at the upper surface by a variable root uptake flux equivalent to transpiration *E*, as driven by the product of a diurnal demand function *f(t)* (in **b**) and *nVPD*. **c** Dimensionless stomatal conductance *g_s_* as a function of leaf water potential *Ψ* [MPa], with the *g_s_Ψ_50_* for a 50% decline in stomatal conductance set at -1.33 MPa (dotted lines). In the modeled domain **d**, water content θ depends on soil hydraulic conductivity *K(h)* [cm day^-1^] and the gradient in matric potential *h* [cm]. **e** Transformation of the water content, matric and water potentials to the potential flux domain [φ, cm^2^ day^-1^] linearizes the governing equation, but results in an extremely steep function for *g_s_* that hinders numerical solution.

Transpiration is represented as the product of a peak transpiration rate (*E_p_*) that carries the units of cm day^-1^ with three dimensionless factors that introduce variations into *E*. The first is a diurnal demand function *f(t)* (Fig. **1b**) that produces a sinusoidal range from 0 to 1, representing the effects of diurnal variations in air temperature on *VPD* in the well mixed air above the canopy as well as variations in stomatal aperture as light levels change over the day. The second is a non-dimensional nominal *VPD* level (*nVPD*) that represents daily peak atmospheric sink strength, and the third is a non-dimensional total conductance (stomatal plus boundary layer, scaled 0 to 1) that depends on leaf water potential and captures the effect of water stress on the realized evaporation rate (Fig. **1c**). In this framework, it is useful to distinguish between ‘Potential transpiration’ *PE*, which represents the transpiration rate that would be realized at maximum total conductance, and the actual transpiration *E* inclusive of stomatal water limitation effects,

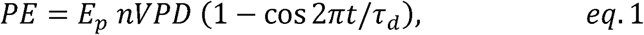

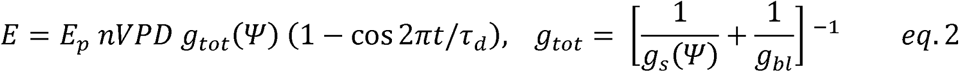

where τ*_d_*is the period of diel oscillation of one day.

In the soil domain changes in water content are related to changes in hydraulic conductivity and flux by Richard’s equation (Fig. **1d**), neglecting gravitational effects over the centimeter scale (Cowan, 1965). The governing equation, boundary conditions, and initial conditions are then linearized using the Kirchhoff transform, leading to the expression of soil water content *θ*, soil matric potential *h*, and leaf water potential *Ψ* in terms of a single variable φ, a potential flux with diffusive units of cm^2^ day^-1^. To organize all the parameters in the model into dimensionless groups that describe the behavior of the solution, φ, time *t* and length *z*, are re-scaled to express the governing equations and boundary conditions as functions of the non-dimensional potential flux Φ= *φ* /(*H J_m_*), time *T=t/*τ*_d_*, and length *Z=z/H*, where *H* is the height of the model domain:

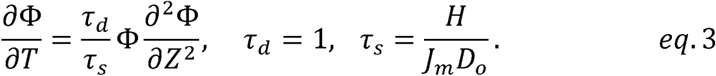

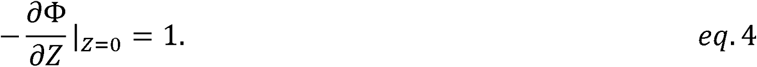

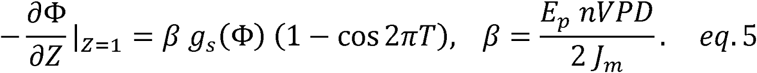

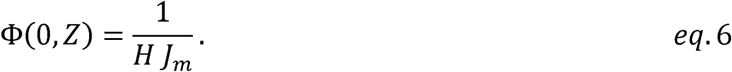

The model takes the form of a non-linear diffusion equation, where the soil diffusivity, *D*(Φ)=*D_o_*Φ, introduces non-linearity through its dependence on the potential flux itself (note that *D_o_* is a pure number that sets the initial magnitude of the soil diffusivity). Conceptually, soil diffusivity represents a ratio of hydraulic conductivity to hydraulic capacity (water storage).

There are two timescales in the problem (eq. 3). The first is the period of oscillations in potential transpiration of one day, τ*_d_*(eq. 2). The second is the diffusive timescale τ*_s_*, the characteristic time for the soil domain to attain a new steady state after a perturbation at the boundary, which is larger (slower) for larger domain sizes (*H*) and lower fluxes from the water source *J_m_*. Larger values of *D_o_* mean that conductance dominates soil capacitance contributing to faster response times (damping is low), while smaller values mean soil capacitance dominates conductance, slowing response times (damping is high). These two timescales appear as a ratio, meaning their relative values determine the time-dependent behavior of the model. However, this behavior is not entirely predictable as *D_o_*Φ varies with time and position in the soil domain, with important consequences.

The remaining parameter driving model behavior is *β* (eq. 5). *β* represents the balance of potential evaporation *PE* to a steady asymptotic flux into the soil domain. As the value of the time-varying term in eq. 5 integrated over a single day is 1, a *β* of 1 means that the *total daily* flux from the source (eq. 4, always equal to 1) balances *total daily PE*, allowing normalized stomatal conductance to remain near its maximum value of 1. Yet, even under this daily balance of supply and demand peak potential transpiration rate exceeds the steady flux into the soil domain by a factor of 2. Such excess demand can be met by the discharge of soil water as φ falls in the domain, and stomatal closure need not occur over the course of the day. If the inward flux from the source is sufficient to re-charge the soil domain overnight, a cycle of daily peak *E* exceeding the inward flux without stomatal limitation can continue. For *β* >1, stomatal regulation will at some point be triggered according to the scale of the imbalance and the ratio of timescales. But there is nothing in the model that forces a balance of inputs and outputs at the daily scale. It is helpful to keep in mind that the *instantaneous* balance of supply and demand seen by the plant is the sum of a steady inward soil flux that originates outside the domain and a transient capacitive or storage flux that originates within the domain. This combined flux can become asymptotic at the root plane at an emergent maximum *larger* than the maximum steady inward flux at the lower domain boundary that is represented in *β*. Note that *β* is numerically equivalent to *nVPD*.

### Parameterization for the Drake experiment

In the Drake *et al*. (2018) experiment, individual eucalyptus trees (∼ 6 m tall) were enclosed in 9 m tall temperature and humidity-controlled gas exchange chambers (*CO_2_*, *H_2_O*). The experiment subjected pre-droughted trees to a ‘Heat Wave’ treatment, combining air temperature (daily peak 43-44°C) and *VPD* stress (daily peak 5-6 kPa), versus a ‘Control’ subject to typical conditions (daily peaks of 28-29 °C, 2-2.5 kPa). Irrigation was withheld, making ground water the only possible source recharging the soil domain. As the trees were enclosed in well-stirred air, total canopy conductance was close to stomatal conductance and leaf temperature close to air temperature (*e.g.,* Fig. 4 in Drake *et al.,* 2018), such that atmospheric humidity was ‘imposed’ at the leaf surface (Jarvis & McNaughton, 1986). The potential transpiration rate *E_p_* was set to 0.3 cm day^-1^ to match the intraday peak of 2 mmol m^-2^ s^-1^ seen in the controls (Drake *et al.,* 2018; Fig. 2c). Peak *VPD* for the controls (∼ 3 kPa) was set as the reference value for normalizing *VPD*. The maximum flux from the water table *J_m_* was set to half of *E_p_*, 0.15 cm day^-1^ in accord with the definition of *β*=1 at *nVPD*=1 (eq. 5). Stomatal conductance is given by a sigmoidal function of leaf water potential (Anderegg *et al*., 2017; Franks & Farquhar, 2007; Levin *et al*., 2019). The shape parameter *b* (MPa) and water potential for 50% stomatal closure (*g_s_Ψ_50_*) were fit together with total plant hydraulic conductance (*K_plant_*; cm day^-1^ MPa^-1^) to reproduce the midday water potentials reported in Drake *et al.,* (2018). Stomatal conductance transforms to Φ (and *g_s_Ψ_50_* to *g_s_*Φ_50_) as,

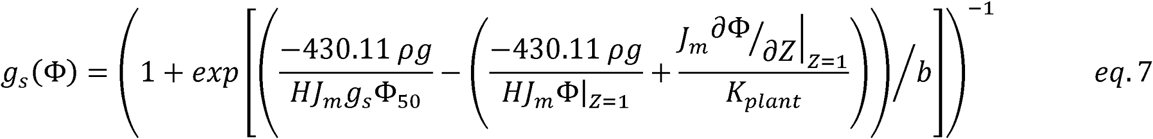

**Figure 2.**
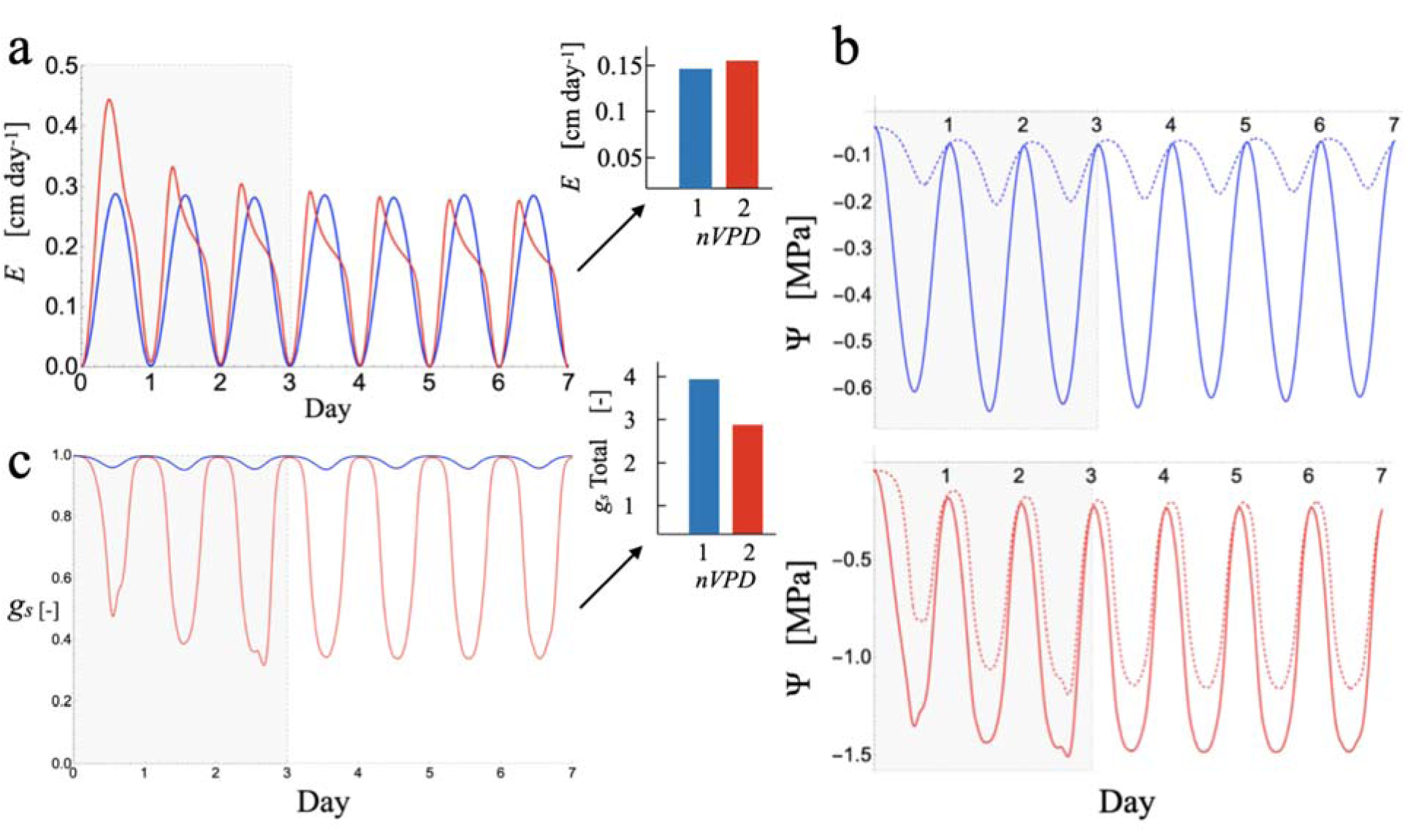
Emergence of feedforward and counter-factual anisohydric behavior in the model for the heat wave experiment in Drake *et al*. (2018) for normalized vapor pressure deficit (*nVPD*) levels of 1 (Control, blue lines) and 2 (Heat Wave, red lines). The first three days (shaded) represent a ‘dry-down’ from a SMC θ = 0.25 used to initialize the model to a four-day quasi-steady diurnal cycle comparable to the physical experiment. (**a**) Diurnal cycles of *E* for the Heat Wave peak earlier and show feed-forward declines relative to the control that conserves total daily *E* (inset bar chart). (**b**) Diurnal cycles of root (dashed) and leaf (solid) water potential *Ψ* show an anisohydric response across *nVPD* levels as leaf *Ψ* more than doubles in magnitude. (**c**) Diurnal variations in *g_s_* show minor *g_s_* limitation at *nVPD* 1 while at *nVPD* 2 *g_s_* is forced to below 50% at midday: inset bar chart of integrated total *g_s_* over four days shows that *A* and *E* become decoupled at high *nVPD*.

**Figure 3.**
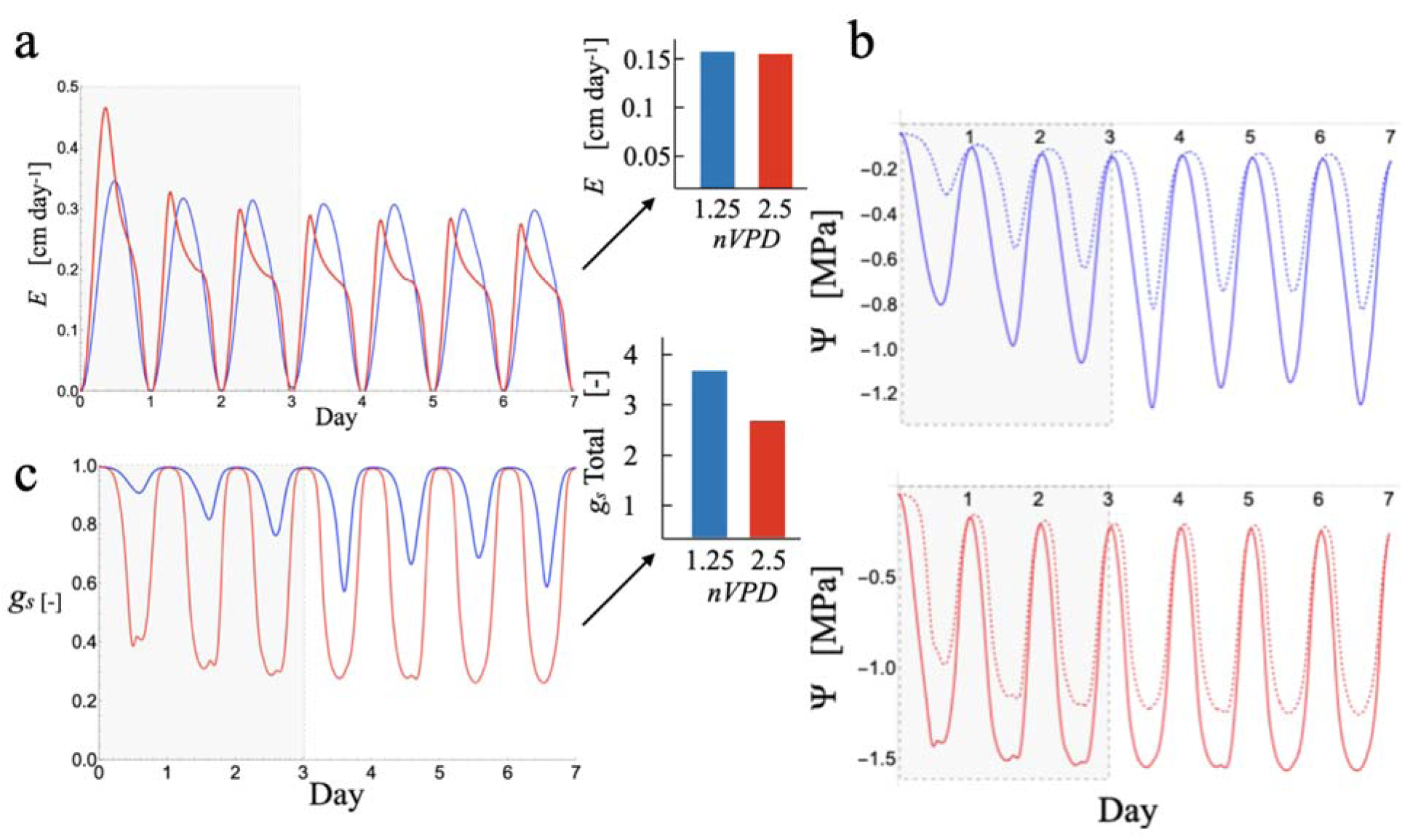
Emergence of feedforward and isohydric behavior as observed in the heat wave experiment in Drake *et al*. (2018) for normalized vapor pressure deficit (*nVPD)* levels of 1.25 (Control, blue lines) and 2.5 (Heat Wave, red lines). The first three days (shaded) represent a ‘dry-down’ from a SMC of θ = 0.25 used to initialize the model to a four-day stationary diurnal cycle comparable to the physical experiment. (**a**) Diurnal cycles of *E* for the Heat Wave peak earlier and show feedforward declines relative to the control that conserve total daily *E* (inset bar chart). (**b**) Diurnal variations in root (dashed) and leaf (solid) water potential create a near ‘isohydric response’ of *Ψ_leaf_* to *nVPD*. (**c**) Diurnal variations in *g_s_* show stomatal limitation at midday at both *nVPD* levels: inset bar chart of total integrated *g_s_* shows *A* and *E* are more decoupled at higher *nVPD*.

**Figure 4.**
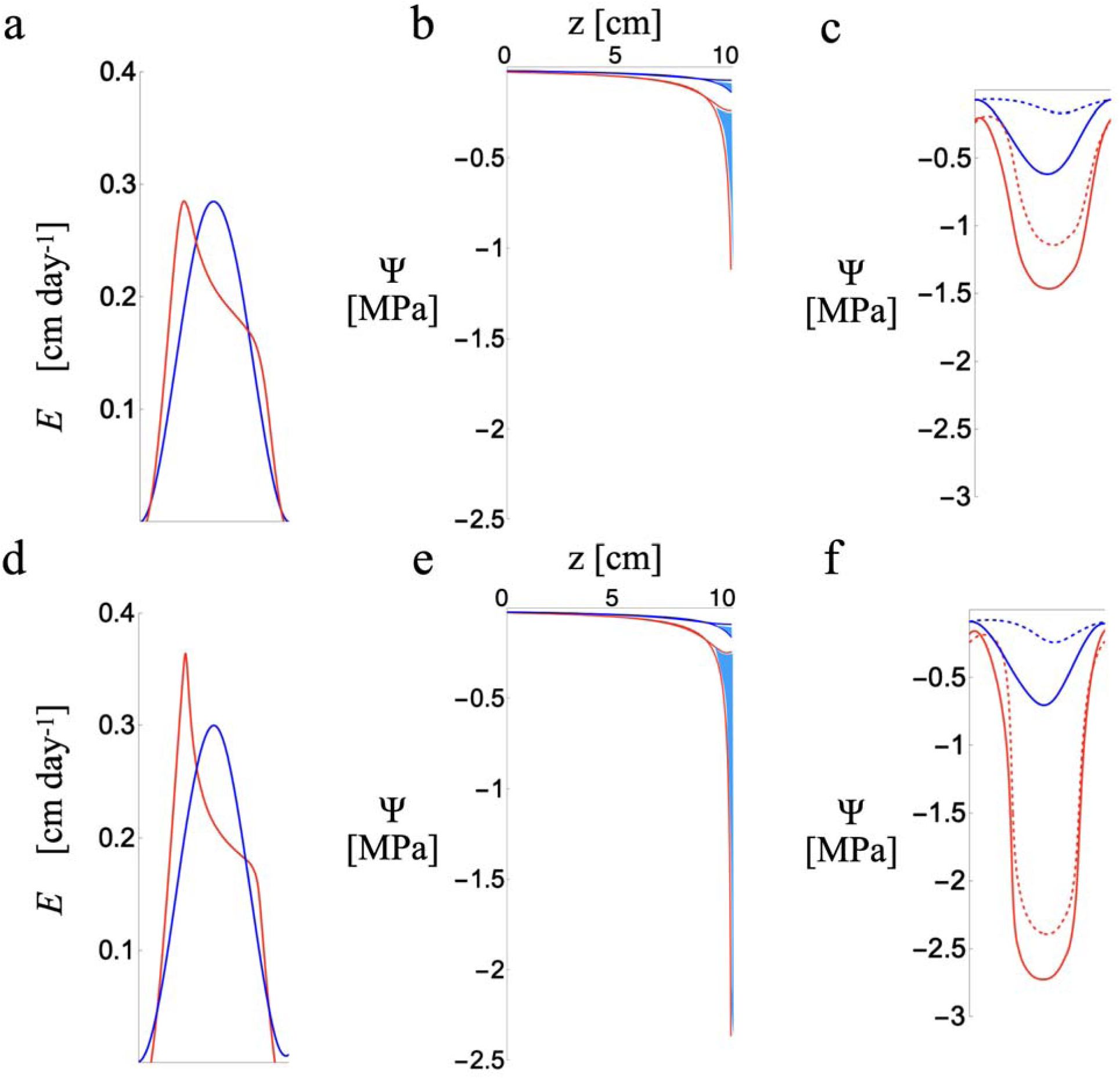
Effect of doubling *g_s_Ψ_50_* (‘osmotic adjustment’) from -1.33 MPa (**a**,**b**,**c**) to -2.66 MPa (**d**,**e**,**f**) with *nVPD* 1 (blue) and *nVPD* 2 (red). **a,d** Diel course of transpiration in the stationary state: maintaining higher *g_s_* to lower *Ψ* stretches the a.m. peak, but *E* for both setpoints converges rapidly once stomata start to constrain the water flux. **b**,**e** Gradients of water potential through the soil domain at the nocturnal maximum and diurnal minimum of *nVPD*: the space between the two lines (light blue) for each level of *nVPD* shows the region of the domain subject to daily fluctuations in water content, which remain confined to near the rooting plane (z=10 cm). **c**,**f** Diel course of soil (dashed) and leaf (solid) water potential; at *nVPD* 2 shifting stomatal closure to a value twice as negative allows leaf water potential to fall to values almost twice as dry but fails to pull a larger midday or total daily water flux from the soil (**a**,**d**).

As soil characteristics were not reported in Drake *et al*. (2018), soil parameters were taken from Cowan (1965).

### Model runs

To generate a range of behavior in the model, a series of values of *β* were investigated ranging from *β=*1 to *β=*2.5. The first three days of each model run represent a transient period during which the initial uniform soil moisture decayed to a repeating daily cycle (stationary-state) comparable to the Drake experiment. To test the effect of a reduction in the water potential for a 50% reduction in *g_s_* (*i.e*., *g_s_Ψ_50_*), a proxy for leaf osmotic adjustment, two additional model runs were compared with *g_s_Ψ_50_* set to -1.33 or -2.66 MPa, and *β=*1 or 2, with other parameters unchanged. The soil diffusivity and *β* were then varied in a series of model runs to explore the general conditions for the emergence of aniso/isohydry and feed-forward behavior. Model runs and figures were coded and evaluated in Mathematica 14 (Wolfram Inc., Champaign IL, USA; Supplemental Information Notes **S2**).

## Results

### Recovery of intra- and inter-day patterns of transpiration under a doubling of VPD

Following a three-day initialization period, the model reproduced the diel stationary states and inter-day conservation of *E* across both treatments seen in the physical experiment (Drake *et al*., 2018; their Fig. **2**). Other behaviors could be produced for some initial conditions, including negative values (foliar uptake and hydration of the soil from the atmosphere at the dew point, not observed by Drake *et al*. (2018)) or longer-phase (multi-day) oscillations of the potential flux (data not shown). The model was judged to have converged on a solution when the total daily flux into the rooting plane reached a stable value of 0.15 +/- 0.005 (3.33%) cm day^-1^. Root surface water potentials were sufficiently dry that no serious errors were introduced by the steady asymptotic flux boundary condition (Fig. 2b, Supplemental Information Figs. **S1**-**S4**), and diurnal variations in φ, *h*, and *Ψ* did not reach the source side of the model domain from the rooting plane, indicating the 10 cm depth chosen for *H* was sufficient to capture all time dependent soil effects (Supplementary Information Figure **S5**).

For the Control conditions of *nVPD* 1 and 1.25 transpiration closely tracked diurnal variations in potential transpiration (Fig. **2a**, **3a** blue). Doubling *nVPD* (and so *β*) to 2 and 2.5 respectively for the Heat Wave treatment showed earlier peaks and feed-forward declines in *E* (*i.e*., *E* falling as *nVPD* is still increasing) relative to the control (Fig. **2a**, **3a** red). Examination of the diurnal cycles of root and leaf water potentials shows that the loss of soil conductivity at the higher levels of *β* and *nVPD* of 2 and 2.5 similarly depressed root and leaf water potential to about -1.2 and -1.5 MPa (Fig. **2b****, 3b** dotted and solid red lines respectively). Yet, the low-stress conditions of *β=nVPD=*1and *β=nVPD=*1.25 showed very different midday water potential depressions of - 0.15 MPa (root) and -0.6 MPa (leaf), versus -0.7 and -1.2 MPa (**Fig. 2b,3b** dotted and solid blue lines for root and leaf respectively). Interpreted as a response to *VPD* (Martínez-Vilalta *et al*., 2014), the doubling from *β=nVPD=*1 to 2 produced an anisohydric shift in midday leaf water potential from -0.6 to -1.5, while the doubling from *β=nVPD=*1.25 to 2.5 showed a more isohydric pattern of -1.2 to -1.5 MPa. This shift in behavior derived from root surface drying pushing stomatal conductance into the region of steep declines near *g_s_Ψ_50_*, resulting in more isohydric regulation of *E* (Fig. **1c**, **2c**, **3c**). In the physical experiment, both Control and Heat Wave leaf water potentials fell in a similar range around -1.5 MPa (Drake *et al.,* 2018) in an apparent isohydric response to demand consistent with the model behavior for doubling *nVPD* from 1.25 to 2.5.

The above results suggest that both *g_s_Ψ_50_* and the degree of soil drying control the transition from anisohydric to isohydric behavior. Yet, running the model for both *nVPD* 1 and 2 with *g_s_Ψ_50_* doubled from -1.33 to -2.66 MPa (Fig. **4**) shows that in response to elevated *nVPD* the water potential at the rooting surface (Fig. **4e** vs **4b**) drops precipitously to force leaf water potential to whatever level is required to limit transpiration to the available soil flux (Fig. **4f** vs **4c**). The ultimate effect of ‘osmotic adjustment’ in the stomatal setpoint *g_s_Ψ_50_* was then to steepen the feedforward behavior of the Heat Wave trees (Fig **4a** vs **4d**) and to lower the minimum leaf water potential (Fig. **4c** vs **4f**) without extracting more water out of the soil over the day.

### General controls on the emergence of feed-forward and isohydric behavior

That the doubling *nVPD* from 1.25 to 2.5 showed an isohydric response of leaf water potential without the emergence of strong feed-forward declines in *E* at *nVPD* 1.25 suggests these behaviors are independent. Sensitivity analyses show that this apparent independence is true only in part (Fig. **5**). Varying *β* from 1 to 5 (with nominal diffusivity *D_o_*=25.7) shows that feedforward control steepens as the imbalance of supply and demand becomes large (Fig. **5a**), while isohydry emerges between *β* 1 and 2, with only a small further decline in midday leaf water potential as *β* grows to *β*=5 (Fig. **5b**). Both the emergence of feedforward and isohydry can result from increasing *β*. Yet, this coupled response is not seen for low values of the soil diffusivity (*i.e.,* higher clay fractions: Jury *et al.,* 1991; Koehler *et al*., 2024; Wankmüller *et al*., 2024)).

**Figure 5.**
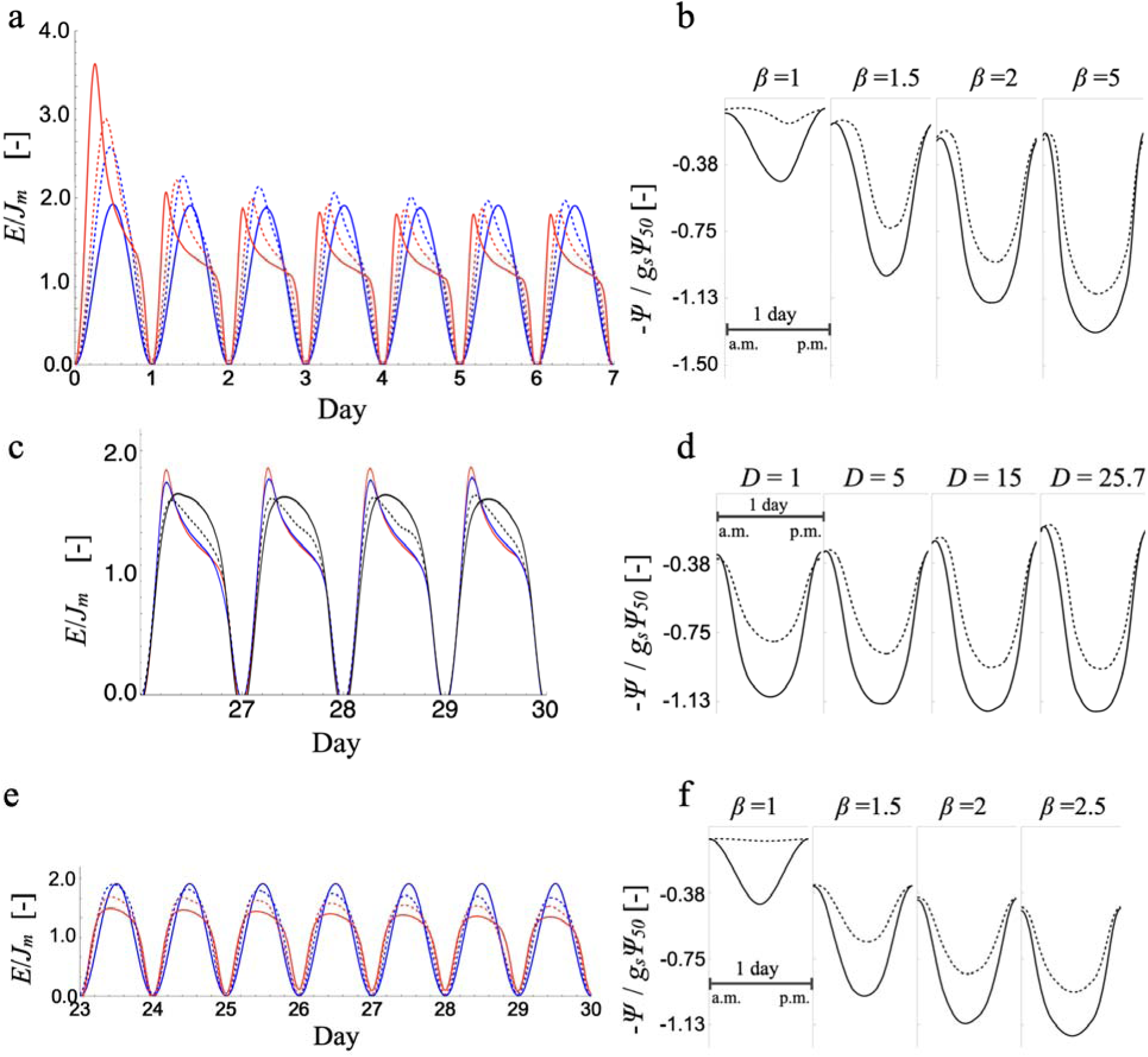
Emergence of feedforward control of transpiration *E* and isohydry in the soil-plant system. *E* (a,c,e) is normalized to the flux recharging the soil domain *J_m_*, and water potential (b,d,f) is normalized to the value at 50% stomatal closure. **a** A parametric sweep across *β* equal to 1 (blue solid), 1.5 (blue dashed), 2 (red dashed) and 5 (red solid) shows the emergence of feedforward control at high *D_o_*=25.7. **b** Diel course of leaf (solid) and root (dashed) water potentials show a transition from anisohydric to isohydric for the same simulations as in **a** as the drying root surface forces stomata to shut. **c** Decay of feedforward control at *β*=2.5 as *D_o_* falls from 25.7 (red) to 15 (blue), 5 (black dashed) and 1 (black solid). **d** Diel course of leaf (solid) and root (dashed) water potentials for the simulations in **c** showing no change in the pattern of midday leaf water potentials but greater relaxation at pre-dawn as *D_o_* falls. **e** Parametric sweep across *β* levels 1 (blue solid), 1.5 (blue dashed), 2 (red dashed) and 2.5 (red solid) showing that at *D_o_*=1 peak broadening replaces the emergence of feedforward control due as *β* increases. **f** The emergence of isohydry at *D_o_*=1 follows the same pattern as at *D_o_*=25.7.

Lowering the nominal diffusivity in steps from 25.7 to 1 while holding *β* at 2.5 shows that feedforward behavior vanishes as the time scales for the soil to respond to diurnal changes in demand become longer, smoothing *E* over the course of the day (Fig. **5c**). The lack of any significant change in the response of midday leaf water potentials shows that isohydry is far less sensitive (Fig. **5d**). This is confirmed by a sweep of *β* from 1 to 2.5 at *D_o_*=1 (Fig. **5e,f**). In the ‘slow’ soil *D_o_*=1 regime, increasing *β* results in peak broadening instead of feedforward behavior (Fig. **5e**), while midday leaf water potentials show the same transition to isohydry as for high *D_o_* (Fig. **5f**). Isohydry develops due to supply imbalance (β >1) alone.

How low soil diffusivity suppresses the development of feedforward behavior in response to *β* >1 can be understood by examining the spread between the gradients of water potential in the soil domain at the minima and maxima of *nVPD* (Fig. **6**). At *β*=1, *E* tracks potential *E* at both low (*D_o_*=1; Fig. **6a**) and high (*D_o_*=25.7; Fig. **6b**) nominal soil diffusivities, yet the soil gradients behave very differently. At the lower diffusivity, the soil is slow to respond to the diel cycle and capacitive discharge is damped and overnight relaxation is small (Fig. **6c**). At the higher nominal diffusivity, the soil response is fast and the soil gradient relaxes overnight to a large degree (Fig. **6e**). This difference is captured by the ‘damping depth,’ δ, the distance from the rooting plane into the soil domain that diel oscillations are expected to reach:

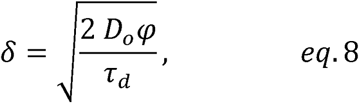

**Figure 6.**
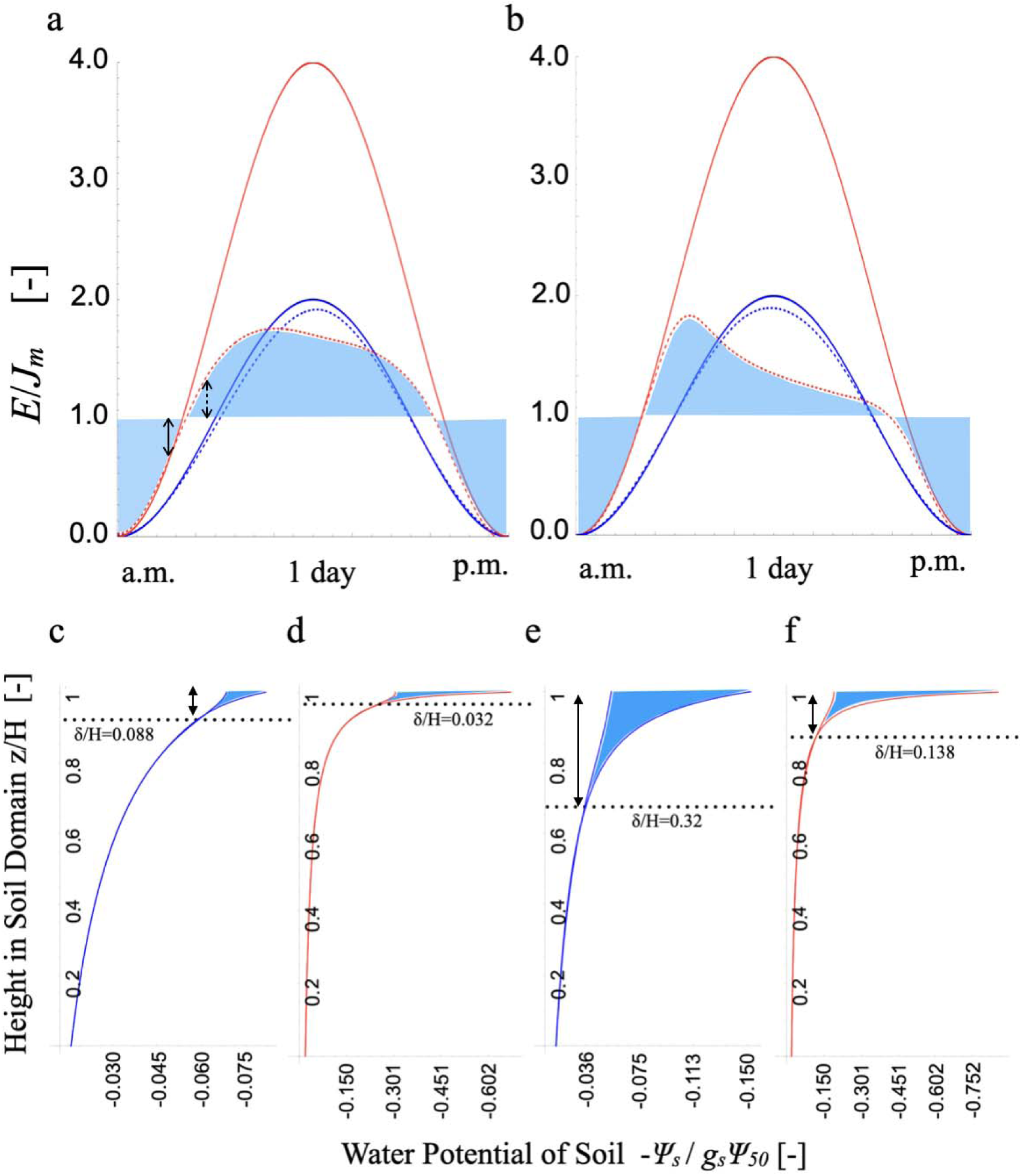
Diel course of potential transpiration (solid) and actual transpiration *E* (dashed) normalized to the inward soil recharge flux *J_m_* at *β*=1 (blue) and *β* =2 (red) and *D_o_* =1 (**a**) and 25.7 (**b**); light blue shading indicates periods of net recharge (below *E*= *J_m_*) and discharge (above *E*= *J_m_*) with respect to the *β*=2 case, with the heights proportional to the magnitude of the recharge/discharge flux at that instant. **c**,**d**,**e**,**f**: Gradients in soil water potential relative to the leaf water potential for 50% stomatal closure at the daily min and max of *E*; the light blue region shows the location and potential range for transient release of water (note the difference in potential scales), relative to the damping depth δ, or how deep into the soil domain diel variations are expected to reach. Panels **c,d** correspond to **a**, and **e,f** to **b.** Note that position in the soil domain *z* and damping depth δ are expressed relative to the domain size *H*.

At low nominal diffusivity, the time-invariant gradient from z=0 to z=9 cm moves the steady flux of 0.15 cm day^-1^ with no capacitive discharge (Fig. **6c,d**), while the midday excess transient flux up to the peak of 0.3 cm day^-1^ is met by capacitive discharge within the last cm. As the gradient hardly relaxes, when *β* is doubled to 2 it is already diverging and the total soil flux (*J_m_* + capacitive discharge) becoming asymptotic at the root plane (compare Fig. **6c** to **6d**). As a result the stomata are rapidly pushed toward their *g_s_Ψ_50_* and *E* fails to follow potential *E* the more it exceeds *J_m_* (Fig. **6a**, dashed red vs solid red lines).

For both fast and slow soils, the doubling of potential *E* relative to *J_m_* spreads the net discharge of stored water over a longer period that starts earlier in the day. This leaves less dischargeable water to meet peak midday demand, leading to reduced midday *E* relative to *β*=1 (Fig. **6a**) for both high and low nominal diffusivity. Yet, at high nominal diffusivity, relaxation of the soil gradient is pronounced relative to low diffusivity as the damping depth extends below 9 cm (Fig. **6b,e**). When potential *E* is doubled at *nVPD* 2, this increased relaxation allows capacitive discharge to be ‘pulled-forward’ in the early a.m. (skewed light blue region in Fig. **6b**) to track potential *E* for longer before the soil gradient diverges (Fig. **6f**). The early peak then collapses in a feedforward manner as soil capacitive water release becomes exhausted. The transient asymptotic level of *E* then decays toward the steady flux (Fig. **6b**, red dashed line) as the soil gradient diverges to push stomata to their *g_s_Ψ_50_* (Fig. **6f**).

Why the collapse in capacitive discharge? Water release becomes increasingly confined to a region near the rooting plane as demand increases. While at *β*=2 a faster soil leads to a deeper damping depth and greater overnight gradient relaxation (Fig. **6d,f**) that allows *E* to track potential *E* farther along its curve (Fig. **6a,b**), the doubling of *β* itself drops the damping depth by more than half (Fig. **6d** vs **6c**, **6f** vs **6e**). This decrease occurs due to the realized diffusivity’s (*D_o_*φ) dependence on the local moisture state of the soil. Evaluating δ (eq. 7) at the rooting plane (*z=H*) selects the lowest value of the soil diffusivity *D_o_*φ in the system, and it is this minimum diffusivity that controls the damping depth (dotted lines Fig. **6c,d,e,f**). As potential *E* increases both in the a.m. and across treatments, *D_o_*φ at the rooting plane decreases, and this shallowing of the damping depth constrains soil water release: the shallow drier soil increasingly feels the ‘squeeze’ of atmospheric demand rather than the deeper wetter soil. Diurnally increasing confinement of the gradient causes capacitive discharge to collapse, pushing stomata into feedforward regulation. This amplification of stress in the soil region nearest the rooting plane then plays an important role in creating the behavior of a ‘hydraulic rope’ that as the system pulls harder breaks strands rather than pulls more water (Sperry & Love, 2015).

How does the steady flux from outside the domain contribute to the above dynamics? An inward flux from deeper soil controls the total amount of water that can be transpired in a diel stationary-state: as *J_m_* goes to zero no stationary state occurs as the total moisture of the root and soil domain declines monotonically (Cowan, 1965), and the maximum asymptotic flux *at the root surface* has only a transient component derived from capacitive discharge. The capacitive confinement effects due to declines in *D_o_*φ over and within days that drive feed-forward behavior will still occur. *β* is not defined for this case, with the balance of atmospheric demand and capacitive soil supply entering instead into the time scale for soil moisture depletion that also controls feedforward and isohydric behavior (Tuzet *et al*., 2003; Supplemental Information Notes S1).

### Interpretation of empirical model parameters

Plotting the instantaneous diurnal values for stomatal conductance against instantaneous *nVPD* (i.e., *nVPD* level multiplied by the diurnal function) results in an apparent stomatal response to *nVPD* (Fig. **7**). Fig. **7a** shows the fit of the *VPD* sensitivity model of Oren *et al*. (1999) *g_s_* = *g_s,ref_* (1- *m* ln[*nVPD*]) to the simulated data. Across all four experimental model runs, the bulk soil moisture at the source-edge of the model domain remained the same (φ ∼ 2, θ ∼ 0.28, Supplemental Information Fig. **S6**), justifying the aggregation of observations of *g_s_* across all *nVPD* and fitting a single *g_s,ref_* and *m.* These parameters are highly sensitive to the specific level of *VPD* chosen as a reference, 3 kPa (Control trees: Drake *et al.,* 2018). Re-basing *nVPD* to the standard reference of 1 kPa results in a value for the slope *m* of 0.42, consistent with previous estimates for soil moisture limited systems (Oren *et al.,* 1999; Novick *et al.,* 2016a; Grossiord *et al.,* 2020). As there was considerable a.m./p.m. hystersis in the relation of *g_s_* to *nVPD,* the reference value at *nVPD* 1 was *g_s,ref_*= 0.78 rather than the expected ‘stress-free’ value of 1: nevertheless the fitted slope of 0.79 picked-out the midday values of *g_s_* as the diurnal spread converged at peak stress.

**Figure 7.**
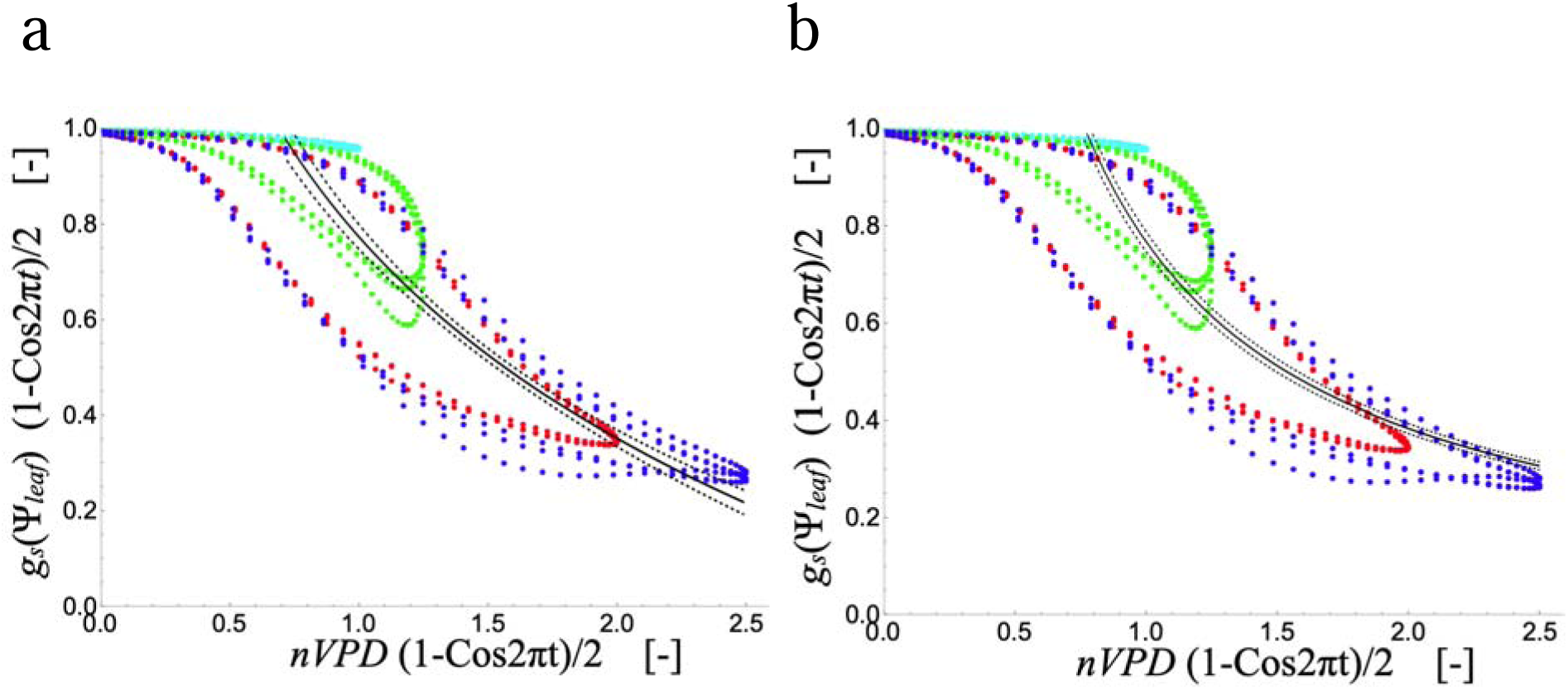
Fits of simulated diurnal values (hysteretic loops) of stomatal conductance for two contrasting models of stomatal sensitivity to *VPD.* Data are for all four modeled *nVPD* levels (cyan 1, green 1.25, red 2, blue 2.5); black solid lines show the fit and dashed lines the 99% CI. **a** Log-based model: *g_s_*=*g_s,ref_* (1- *m* ln[*nVPD*]), *g_s,ref_* =0.78, *m* = 0.79, adjR^2^ = 0.96. **b** Mass-balance based model: *g_s_*= *g_s,ref_* /*nVPD, g_s,ref_* =0.78, adjR^2^ = 0.96.

A second empirical model of *VPD* sensitivity is that stomata adjust to conserve a steady-state balance between liquid and vapor phase transport, forcing *g_s_* to fall as 1/*vpd* (Fig. **7b**; Oren *et al.,* 1999; Grossiord *et al.,* 2020; Mencuccini et al., 2024). For the Drake experiment, a steady mass-balance holds on average over a diel stationary-state, taking the form *g_s_*= *g_s,ref_* /*nVPD* (Fig. **7b**; derivation in Supplemental Information Notes **S1**). Fit to the same simulated data as the log-based model, neither is preferred based on fit statistics, and both capture the peak-stress relationships between *nVPD* and *g_s_* that are taken as representative of a system (Martínez-Vilalta *et al*., 2014; Hochberg *et al*., 2018; Knipfer *et al*., 2020).

The two empirical approaches do predict contrasting behaviors of *E* that coincide with the physical experiment at different time scales (Fig. **8a,b** top panels). For the log-based model the response of *E* transitions from feedback to feedforward at about 2 *nVPD*, qualitatively consistent with the diurnal development of feedforward responses in the physical experiment (Figs. **2****, 3**; Drake *et al.,* 2018), and responses observed in leaf-scale gas-exchange experiments (Mott & Parkhurst, 1991). Yet the log-based model fails to reproduce the observed conservation of *E* across days (Fig. **8a**). The mass balance approach by contrast could *only* capture conservation across days, as it predicts a single value of *E* independent of diel variations in *nVPD* (Fig. **8b**).

**Figure 8.**
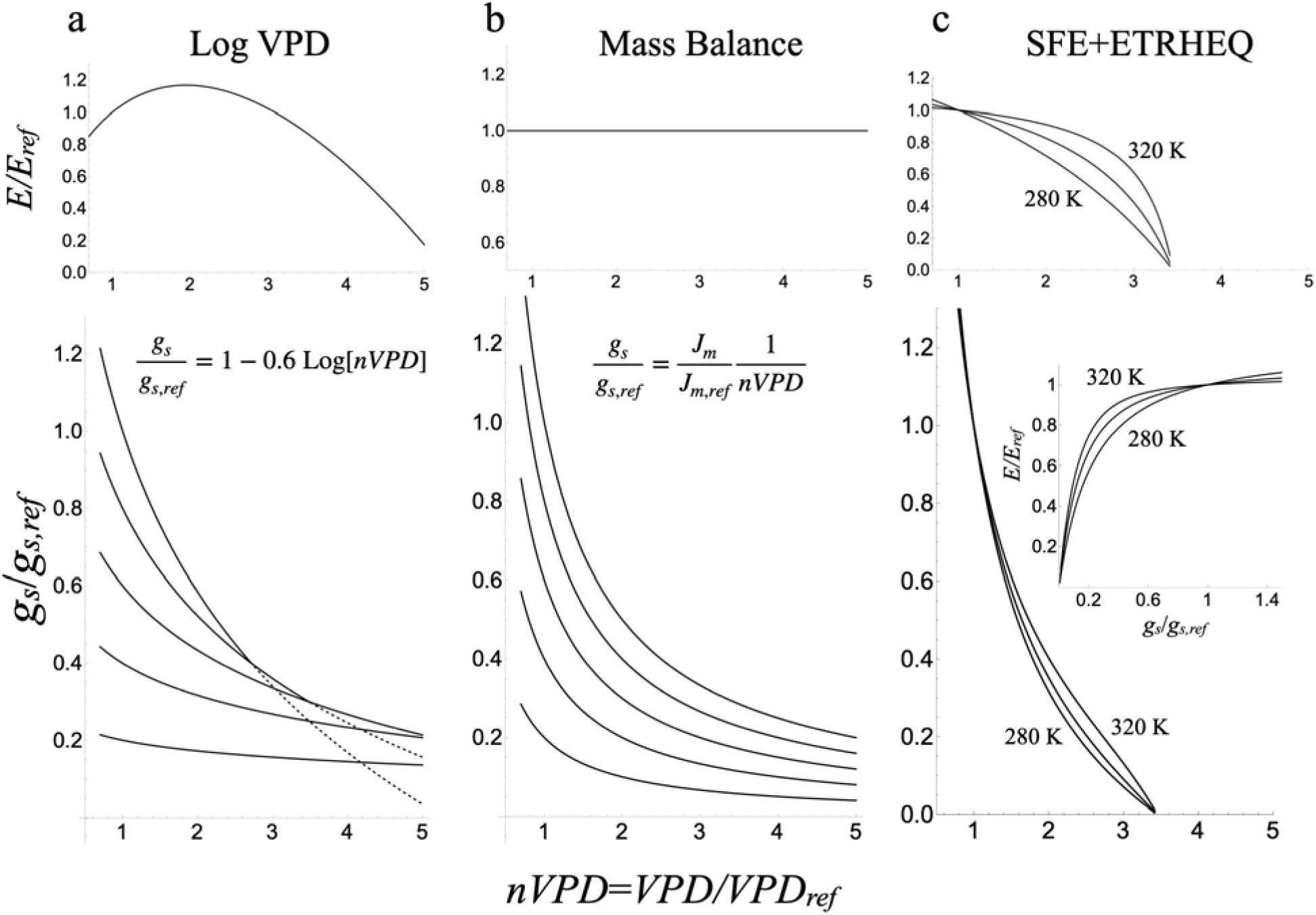
A comparison of the normalized conductance extinction curves (*g_s_*/*g_s,ref_*) and transpiration responses (*E/E_ref_*) for three simplified models of stomatal responses to soil drying and atmospheric aridity, with 1 kPa as the reference state for *nVPD*. **a** Based on the behavior of the dry sites in figure 2 of Novick *et al*. (2016), as the soil dries in the model *g_s_*/*g_s,ref_* = *a* (1- *m* ln[*nVPD*]) the intercept *a* falls from 1 at *nVPD* =1 to 0.2 while the slope *m* falls from 0.6 to 0.2. Note that the curves representing *nVPD* responses at higher soil moisture are predicted to cross (at the transition from solid to dashed) the curves for lower soil moisture. **b** For the mass balance model *g_s_*/*g_s,ref_* = (*J_m_* /*J_m,ref_*) (1 / *nVPD)*, *J_m_* /*J_m,ref_* falls from 1 to 0.2 to simulate a falling maximum flux as the soil dries. **c** Under the condition of Surface Flux Equilibrium (SFE), the atmospheric boundary layer approaches a constant RH (and potential *nVPD*) at all heights; responses are shown for air temperatures at two meters above the canopy of 280, 300 and 320 K.

In the full model presented here, static declines in bulk soil water potential lower the maximum stomatal conductance that would pertain at zero *E*, while non-zero *E* induces a dynamic drop in water potential that drives feed-forward stomatal behavior. The log-based model renders this partitioning between static (*g_s_*,_ref_) and time-dependent dynamic (*m*) effects as steady-state behaviors, and an apparent pathology of this model structure is that at high *VPD* the model predicts lower *g_s_* at higher soil moisture than at low (dotted lines in Fig. **8a**). By contrast, in the mass balance model as the soil dries out falling *J_m_* generates a more realistic sequence of curves (Fig. **8b**) that decay toward a converging response to *VPD*.

### The contribution of boundary layer feed-backs to apparent stomatal VPD responses

The above discussion applies directly to the case that stomata represent the controlling resistance between plants and a well-mixed atmosphere. Yet, over time transpiration can humidify the air within and above the canopy to create a feed-back on *E* as the driving force for transpiration decays. Treating the state of the soil and *VPD* as independent variables represents a zero-coupling limit of land-atmosphere interactions that then may not pertain at the time and spatial scales of interest (McColl *et al*., 2019; McColl & Rigden, 2020). For a fully coupled land-atmosphere system in steady-state, ETRHEQ/SFE picks out the surface conductance consistent with the observed RH of the ABL (Fig. **8c**; see Supplemental Information Notes **S1**). At high RH (low VPD) states of SFE, large reductions in initially high values of *g_s_* are associated with small reductions in RH (increases in *VPD*). Yet in this humid limit large changes in *g_s_* are associated with very small changes in *E* (Fig. **8C**, inset). This behavior recalls that of the thick boundary layer limit (Jarvis & McNaughton, 1986) and equilibrium evaporation (Raupach, 2001).

### Limitations of the model

That the neglect of the cylindrical geometry of the soil around a root did not materially affect the results was confirmed by a comparison of the solutions for *β=*1 and 2 in radial and cartesian domains: in both cases for high *β* the potential at the root surface diverges to force stomatal closure (Supplemental Information Fig. **S6**). Similarly, the neglected possibility of a decline in plant hydraulic conductance would be expected to reduce how far root surface potentials diverge to shut stomata, but not to qualitatively change the emergence of a limiting soil flux. The steep non-linearity of the stomatal conductance function in the φ domain (Fig. **1e**) did appear to introduce numerical artifacts evident as sharp deviations from otherwise smooth curves (*e.g*., Day 1 and 4 minimum *g_s_* in Fig. **3c**). Such artifacts could be diagnosed by their vanishing in model runs after small changes to the initial conditions.

## Discussion

### Limitation of transpiration by an emergent maximum flux as a general property of the land surface

The emergence of a steady-state asymptotic flux is a general feature of passively driven flow through media in which increasing driving force feeds back negatively on the conductivity, like plant xylem (Sperry & Love, 2015) or soil (Cowan, 1965; Carminati & Javaux, 2020). While most previous work has focused on steady-states, this study shows how diurnal time-dependent declines in soil diffusivity can drive the increasing confinement of the soil gradient to near the root surface, causing the transient contribution to the maximum soil flux to decay. This decay is then transduced by the functional dependance of stomata on leaf water potential into feedforward control. Iso/anisohydric transitions emerge over longer times scales as either the quasi-steady component of the maximum soil flux declines or peak VPD stress increases. With respect to the generality of the Drake *et al*. (2018) results, the conserved transpiration rate of 1.5 mm day^-1^ obtained in the chambers was well-within the range for *Eucalyptus* forest and plantations, as well as North American temperate forest (Forrester et al, 2010; Zolfaghar 2017; Shveytser 2024). The generality of the modeled stomatal behavior is supported by the empirical parameter fits to the aggregated *g_s_* data, and the underlying soil and stomatal water potential ‘set points’ are within the expected values for a survey of soils and17 diverse tree species (Carminati et al, 2026).

Predicting maximum steady soil fluxes at scale hinges on the availability of soil texture data (Wankmüller *et al*., 2024), source depth (*e.g.,* a water table: Miguez-Macho & Fan, 2021) and rooting depth (Fan *et al*., 2017). Fan *et al*. (2017) describe how root growth follows infiltration depth; if this depth is shallower than the capillary fringe of ground water, a ‘dry gap’ is created between the rooting zone and deep soil moisture. For 501 taxa, Fan *et al*. (2017) found maximum rooting depth clusters between ∼25 cm and 2 m, with the highest density at 1m, paired with water table depths clustered between 1 and 10 m. Meter-scale dry gaps during drought may then occur for many, if not most, woody plants. Some trees obligately produce dimorphic roots, building both a shallow root system to access nutrients and seasonally available rainfall, as well as deep roots that cross a dry gap to reach groundwater and avoid seasonal drought (Skelton *et al*., 2021), shifting the characteristic depth of root uptake. Such deep sources need not be saturated (*i.e.,* a water table) to supply a steady maximum flux, just approximately fixed at some potential over the time scale of interest (Supplemental Information Notes S1; Fig. S4).

Successfully modeling a stationary-state with a flux of moisture from outside the root zone to a plane of root uptake implies that the root-to-shoot ratio no longer mediates soil and shoot fluxes as it does for uptake from an evenly-moist soil volume: this may be realistic when only the wettest part of the root zone matters (Dawson *et al.,* 2020). Yet, to the extent an external inward flux can be spread evenly over the total root surface area, greater rooting volume and density will damp away feedforward behavior, but not the emergence of isohydry (Supplemental Information Notes S1).

### The surprising sufficiency of stomatal conductance as a function of leaf water potential

The analyses presented here demonstrate that a model of stomatal aperture as a function of a single integrative variable – water potential – is sufficient to capture both anisohydric to isohydric transitions (Knipfer *et al*., 2020; Javaux & Carminati 2021) and feedforward responses to *VPD*. This is surprising as it was the failure of leaf water potential *ψ_L_* and *g_s_* to follow a 1:1 relationship in response to *VPD* and soil drying over physiological time scales that motivated work on the role of chemical signaling in stomatal behavior (Turner *et al*., 1984, 1985; Gollan *et al*., 1985; Tardieu & Davies, 1992; McAdam & Brodribb, 2016).

One possible reason for the failure of *g_s_* to track *ψ_L_* is that measurements of bulk leaf water potential may not capture the water status relevant to stomata (Rockwell *et al*., 2014; Buckley *et al*., 2017). Measurements with fluorescing nanogel reporters of water potential point to the existence of large (up to MPa scale) gradients in water potential from leaf xylem to the transpiring surface (Jain *et al*., 2024b,a), which may explain why Anderegg *et al*. (2017) found that a Ball-Berry based model of stomatal conductance that included both a leaf water potential and a *VPD* sensitivity term was preferred: The bulk measure of leaf water potential may be dominated by leaf tissue that sits close to leaf xylem potential, while a *VPD* sensitivity term may capture gradients within the leaf proportional to *g_s_*VPD/K_m_*, such that there is no ‘double-counting’ (Liu *et al*., 2020b). In this view, it seems relevant that in an influential series of papers demonstrating the failure of *g_s_* to track *ψ_L_* (Turner *et al*., 1984, 1985; Gollan *et al*., 1985), leaf water potential was measured by in-situ psychrometers, a technique shown to track stem xylem potential at the point of leaf insertion (Matzner & Comstock, 2001). Unsaturation in the leaf air spaces (Márquez *et al*., 2024; Cernusak *et al*., 2024; Diao *et al*., 2024) could also be a factor, yet, as noted by Grossiord *et al*. (2020), this effect should be ‘passed through’ by parameter observations that ultimately deliver correct predictions of stomatal responses.

### Biological vs physical controls on surface conductance and empirical model parameters

Mott & Parkhurst (1991) showed that stomata respond to *E* rather than directly to *VPD* or RH at the leaf surface. Yet, currently there is no complete description of the mechanisms within a leaf that are involved in feedforward and feedback type stomatal responses to *VPD* (Grossiord *et al*, 2020). The evident ability of a physical null model in this study to generate these behaviors does not mean that biology does not matter for *VPD* responses. Rather, a successful null model is useful for exposing what biological parameters are likely to matter most. With respect to the mechanistic interpretation of empirical model parameters, the model suggests that *m* and *g_ref_* are responsive to different physical forcings originating from different moisture states within the same soil (rhizosphere vs bulk). While correlated across biomes (Novick *et al.,* 2016a), these parameters may be to some degree uncoupled across species by biological factors like variations in root density, hairiness and root exudation that impact the scale and degree of soil-root contact (Carminati *et al*., 2009, 2016; Wankmüller *et al.,* 2024; Akale *et al.,* 2025). In this regard *m* may be expected to act like a biologically influenced parameter partially independent of bulk soil moisture and *g_ref_*.

From the perspective of optimization models of stomatal conductance, feedforward behavior and midday closure have been interpreted as conserving a target proportionality between carbon gain and water loss (Novick *et al*., 2016b; Anderegg *et al*., 2017; Dewar *et al*., 2018; Wang *et al*., 2020; Gardner *et al*., 2023). In the null model presented here, feedforward behavior is decidedly sub-optimal as it derives from the decay of excessive rates of water loss in the a.m. The suggestion is not that stomatal behavior in no-way optimizes instantaneous water-use efficiency, rather that optimization may also occur over longer time scales through adjustments of root-to-shoot ratio, root depth, or soil-root contact that effectively decrease *β* and premature peaking of *g_s_* out of phase with peak irradiance.

### Stomatal responses to VPD and the control of transpiration

In the ‘de-coupled limit’ of a thick boundary layer at the canopy scale (Jarvis & McNaughton, 1986), large changes in stomatal aperture achieve small changes in transpiration. SFE also predicts that at high RH in the ABL large reductions in surface conductance are associated with small reductions in *E*. Changes in RH in the ABL feedback on transpiration as a change in driving force, but there is no feedback of RH on the value of the surface conductance. That SFE can explain ETRHEQ and patterns of *g_s_*versus *VPD* from flux tower data (McColl & Rigden, 2020) suggests that at this scale apparent responses to *VPD* are not driven by physiological feed-backs of *VPD* on *g_s_*. In the moist land surface and atmosphere limit, the important intuition that emerges is not that *g_s_* is highly sensitive to a stress imposed by atmospheric aridity, but that *g_s_* exerts weak control over the evaporative flux.

In the dry limit, the apparent sensitivity of *g_s_* to *VPD* is less but the sufficiency of a theory lacking a feedback of *VPD* on *g_s_* still pertains. While for mesic systems *VPD* responses at the daily time scale show a single curve across all soil moisture quantiles as expected by ETRHEQ/SFE (Novick *et al*., 2016a), arid systems tend to show a family of curves comparable to the response of empirical models described here (Fig. **8**). This difference may arise due to drier systems requiring longer than daily time scales to reach Surface Flux Equilibrium (SFE) (McColl & Rigden, 2020). Yet once at SFE, stomatal behavior may be as physically constrained by SFE/ETRHEQ as the formation of a soil crust. (Vargas Zeppetello *et al*., 2023). From this perspective, the critical biological plant factors contributing to the partitioning of surface latent and sensible heat fluxes are likely those that mediate root and soil contact (Carminati *et al*., 2016) and so scale the maximum flux available to the evaporative surfaces of the canopy, rather than autonomous stomatal controls.

## Conclusion

The influence of *VPD* on stomatal aperture is strongly mediated by the state of the soil, and variations in apparent responses of stomatal conductance to *VPD* at constant bulk soil moisture are not necessarily under biological control. The emergence of isohydry is driven by the integrated daily balance of potential transpiration to maximum soil supply (*β*, in the stationary-state). Emergence of feedforward regulation *E* is time-dependent and driven by both supply and demand imbalance and the soil diffusivity, which is soil texture dependent. While steady-state empirical model parameters can be mapped to hydraulic mechanisms, these reduced models struggle to capture relevant responses of *E*. These results motivate further investigation into the physical and biological factors controlling the surface conductance coupling plant canopies to the atmospheric boundary layer.

## Acknowledgements

Abraham Stroock, Guido Salvucci, Kaighin McColl, Daniel Short Gianotti and N.M. Holbrook provided valuable feedback and discussion. This work was supported by NSF-MRSEC DMR-2011754.

## Data availability

No new data were generated in this study, and all simulated data are available by evaluating the code given in Supplemental Information Notes **S2**.

Notes **S1**. Discussion and derivation of governing equations and boundary conditions.

Notes **S2**. Computer code for model runs and figures.

Figure S1. Maximum soil fluxes for fine sandy loam and clay soils.

Figure S2. Maximum soil flux for the light clay soil used in the model.

Figure S3. Supply curve for a clay soil as the rooting plane dries.

Figure S4. Supply curves for unsaturated soil moisture sources.

Figure S5. Evolution of the potential flux φ for all modeled levels of *nVPD*.

Figure S6. Comparison of cylindrical and cartesian soil and root domain solutions.

---

**Figure S1.**
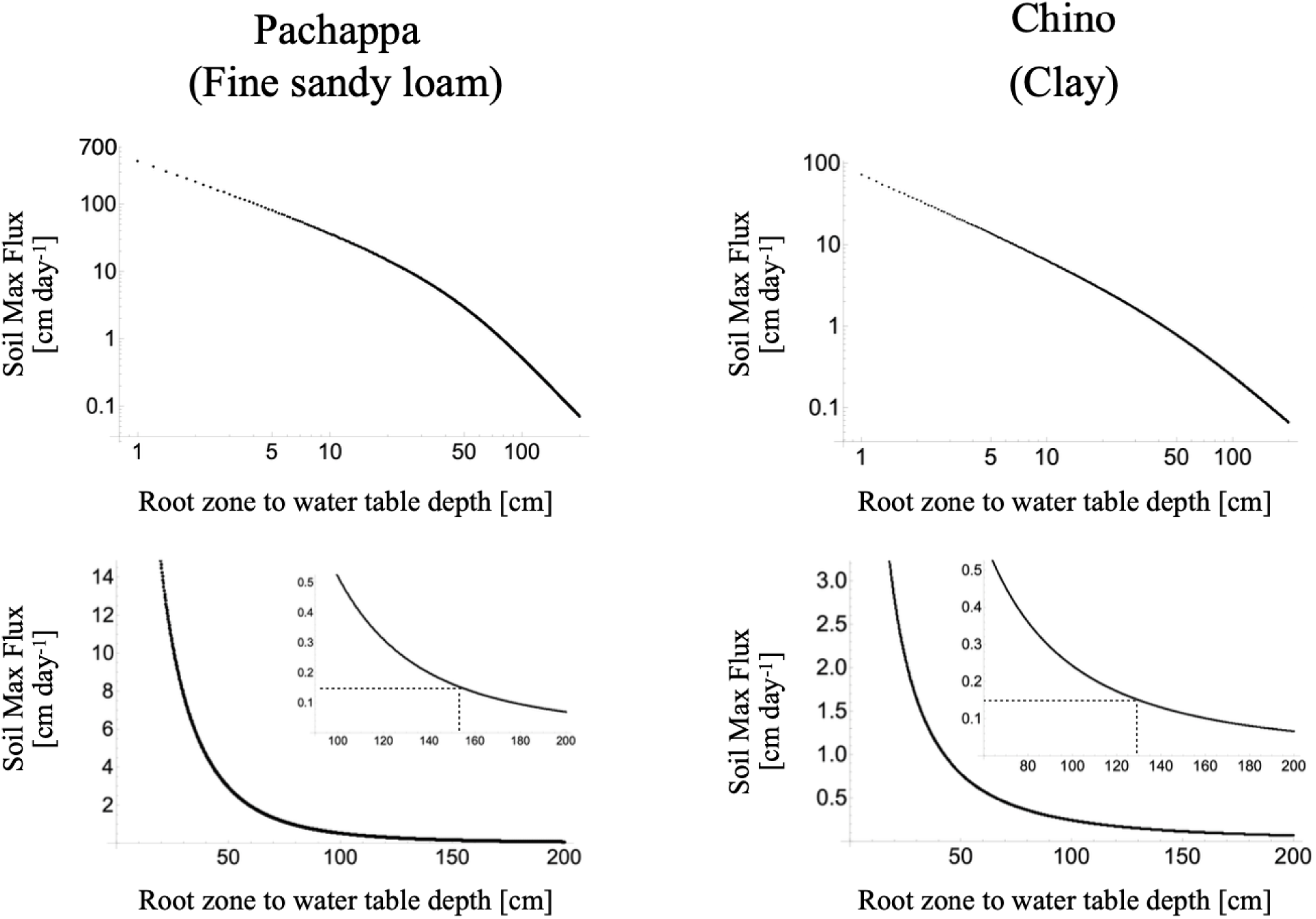
Maximum fluxes [cm day^-1^] for two soils parameterized by Jury *et al*. (1991). Although the saturated conductivities of the two soils differs six-fold, from 2 cm day^-1^ for the clay soil to 12 cm day^-1^ for the more porous fine sandy loam, the maximum soil fluxes converge over longer distances from water table to rooting zone. The flux chosen to match the daily totals for transpiration in the experiment of Drake *et al*. (2018) of 0.15 cm day^-1^ occurs over 130 cm for the clay, and 19% farther (over 155 cm) for the fine sandy loam (inset graphs with dotted lines).

**Figure S2.**
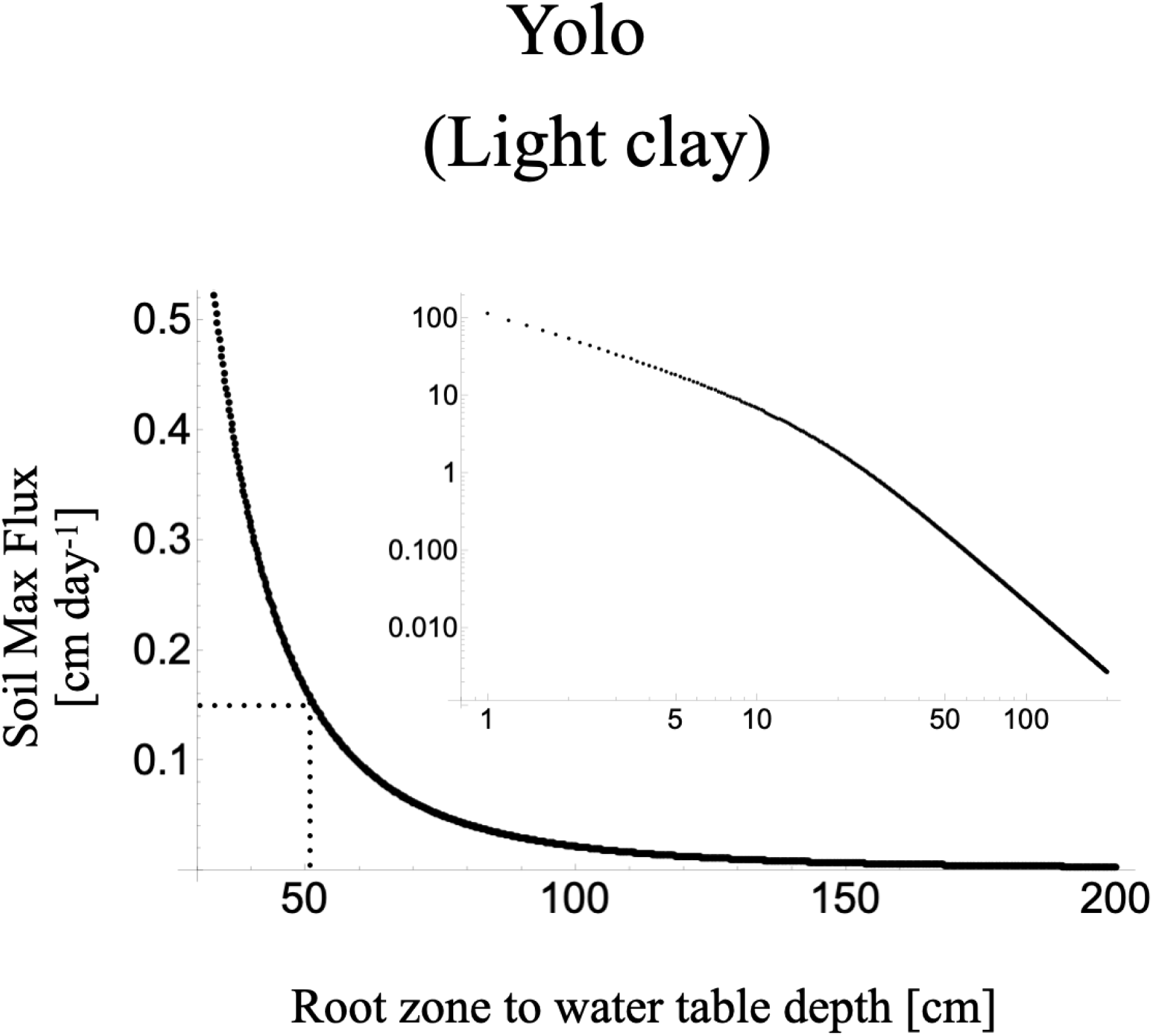
Maximum soil flux in the limit of an infinite driving force as a function of the distance from the water table to an uptake surface (i.e., surface evaporation, rooting plane). Data were generated by *eq*. (5) for the case the exponent *n* = 3, with *a* = -11 cm and *K_s_* = 9 cm day^-1^, based on a fit over a range of soil moisture θ from 0.1 to 0.25 by Cowan (1965) to data in Philip (1957) for a Yolo light clay soil reaching field capacity at θ = 0.5.

**Figure S3.**
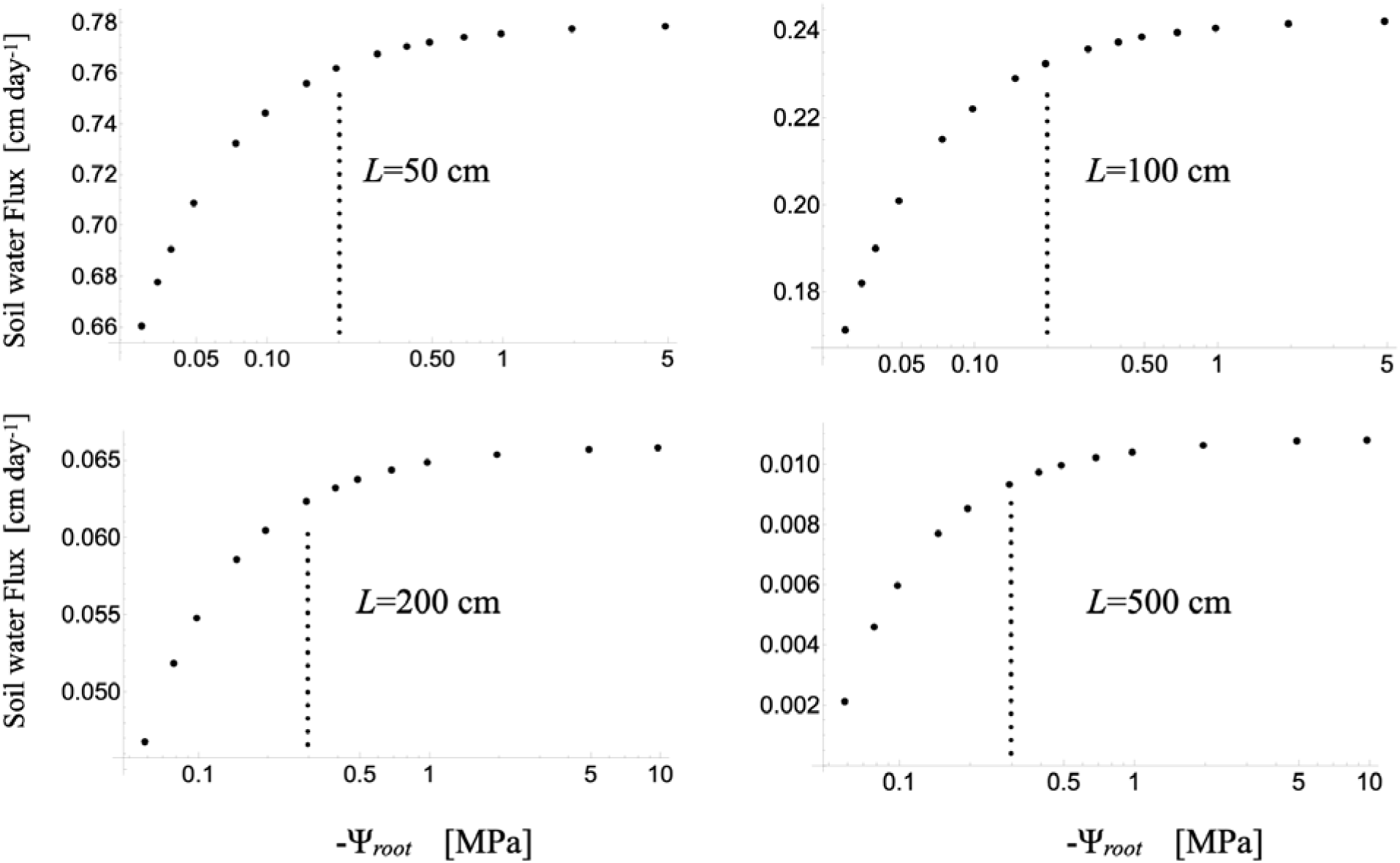
Supply curves for the water flux in a Chino clay soil from a water table to an uptake surface (i.e., rooting plane) located a distance *L* above, as a function of the water potential of that surface (*Ψ_root_*). As the root pulls with only 0.2 MPa of tension over moderate distances (50 and 100 cm), the flux approaches 95% (dotted vertical lines) of the asymptotic maximum fluxes (Fig. 2) for an infinite driving force. Over longer distances for *L* of 200 to 500 cm, the tension required to reach 95% of the maximum flux increases to a still moderate 0.3 MPa.

**Figure S4.**
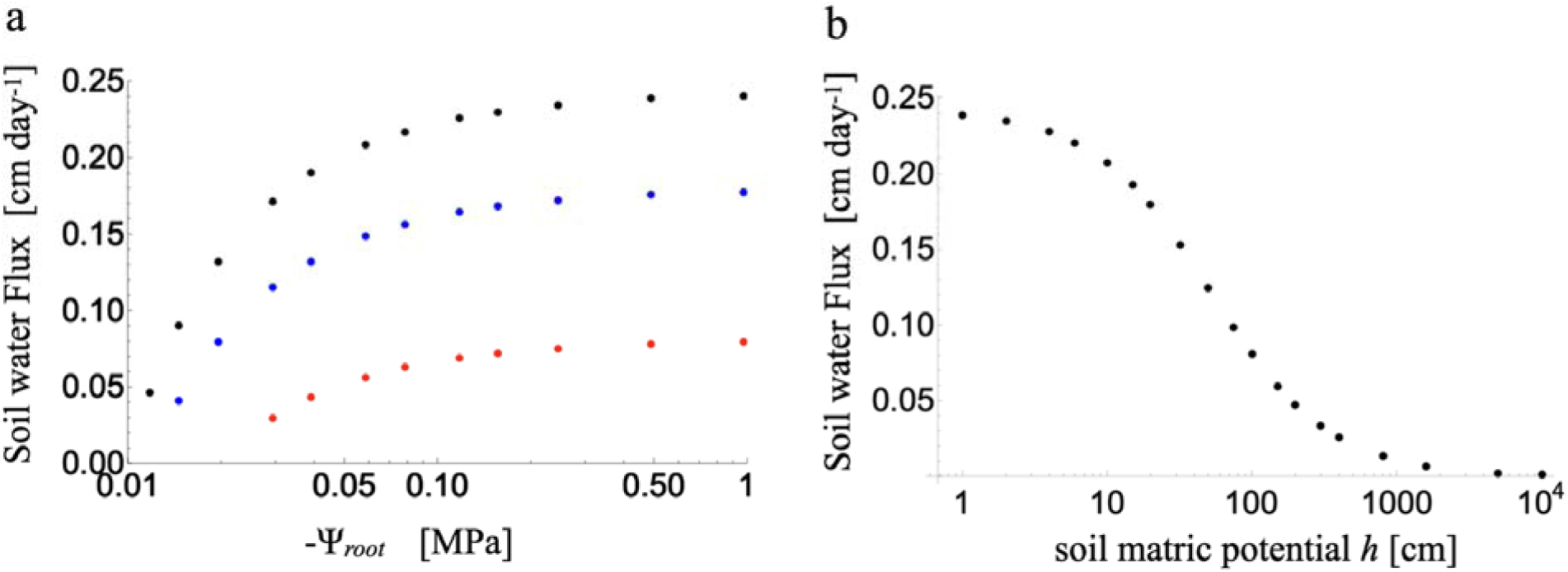
Effects of an unsaturated source of deep soil moisture on asymptotic soil water fluxes. **a** Supply curves in a Chino clay soil for a root located *L*=1 meter above a source with a saturated fixed matric potential of 0 (black), versus an unsaturated source held at -20 cm (blue) or -100 cm (Red), as a function of finite root water potentials. All three soil water fluxes reach an asymptotic maximum flux as root water potentials become more negative than approximately -0.2 MPa. **b** Decay of the maximum soil water flux to an infinitely dry sink *L*=1 meter away as a function of the source soil matric potential for a Chino clay soil.

**Figure S5.**
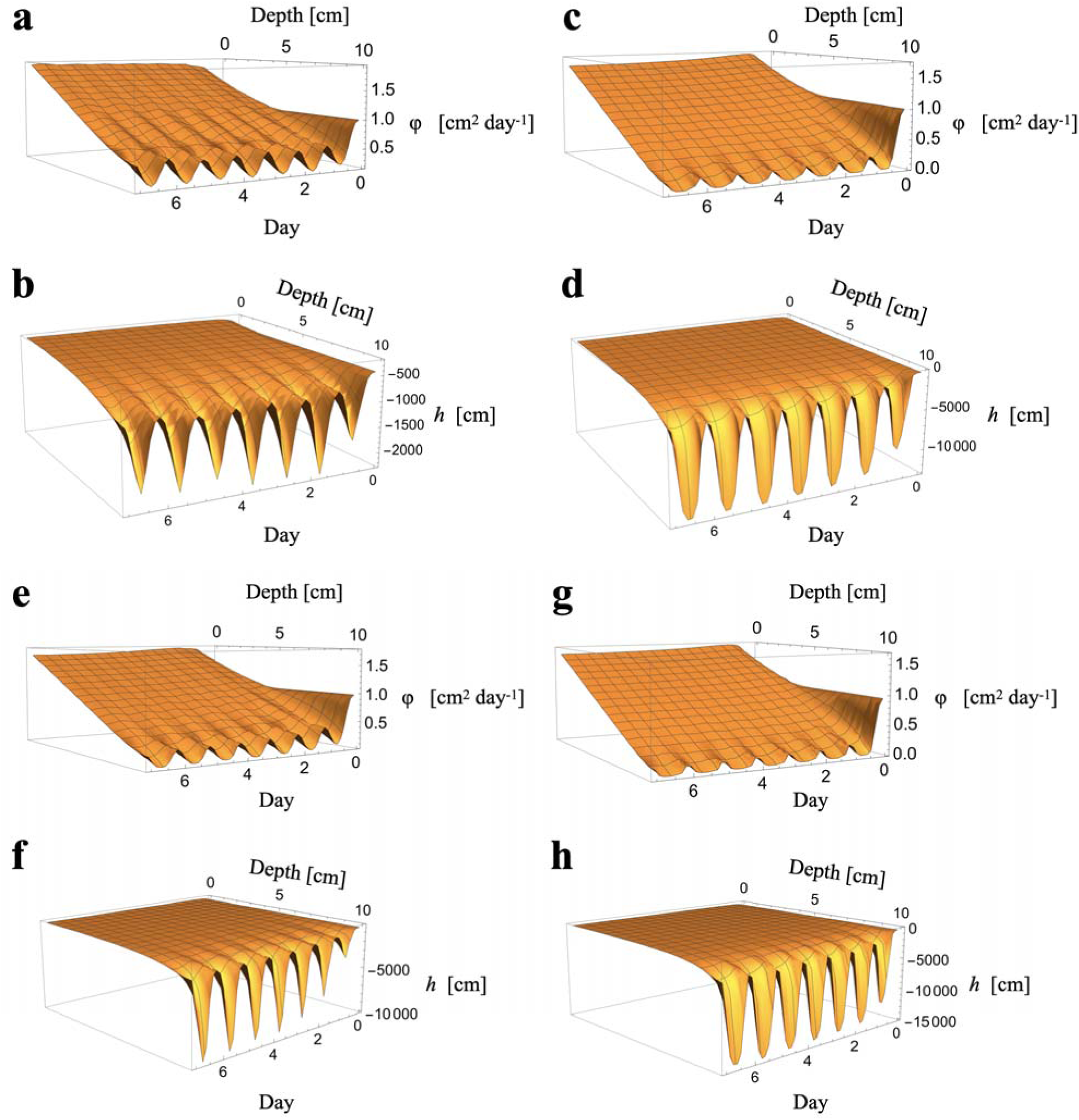
Evolution of the potential flux φ for all levels of *nVPD* (**a**,**b** 1; **c**,**d** 2; **e**,**f** 1.25; **g**,**h** 2.5) over the course of the seven-day simulation. Transformation of the results for the potential flux in **a** and **c** into hydraulic head *h* are shown in **b** and **d** respectively. Note that the relationship between water potential and head is approximately 10,000 cm ∼ 1 MPa. Note that while the gradients in potential flux evolve to nearly linear with only small diurnal fluctuations at the rooting plane, in the physical matric potential domain all of the large and diurnally varying declines occur in the last centimeter before root absorption.

**Figure S6.**
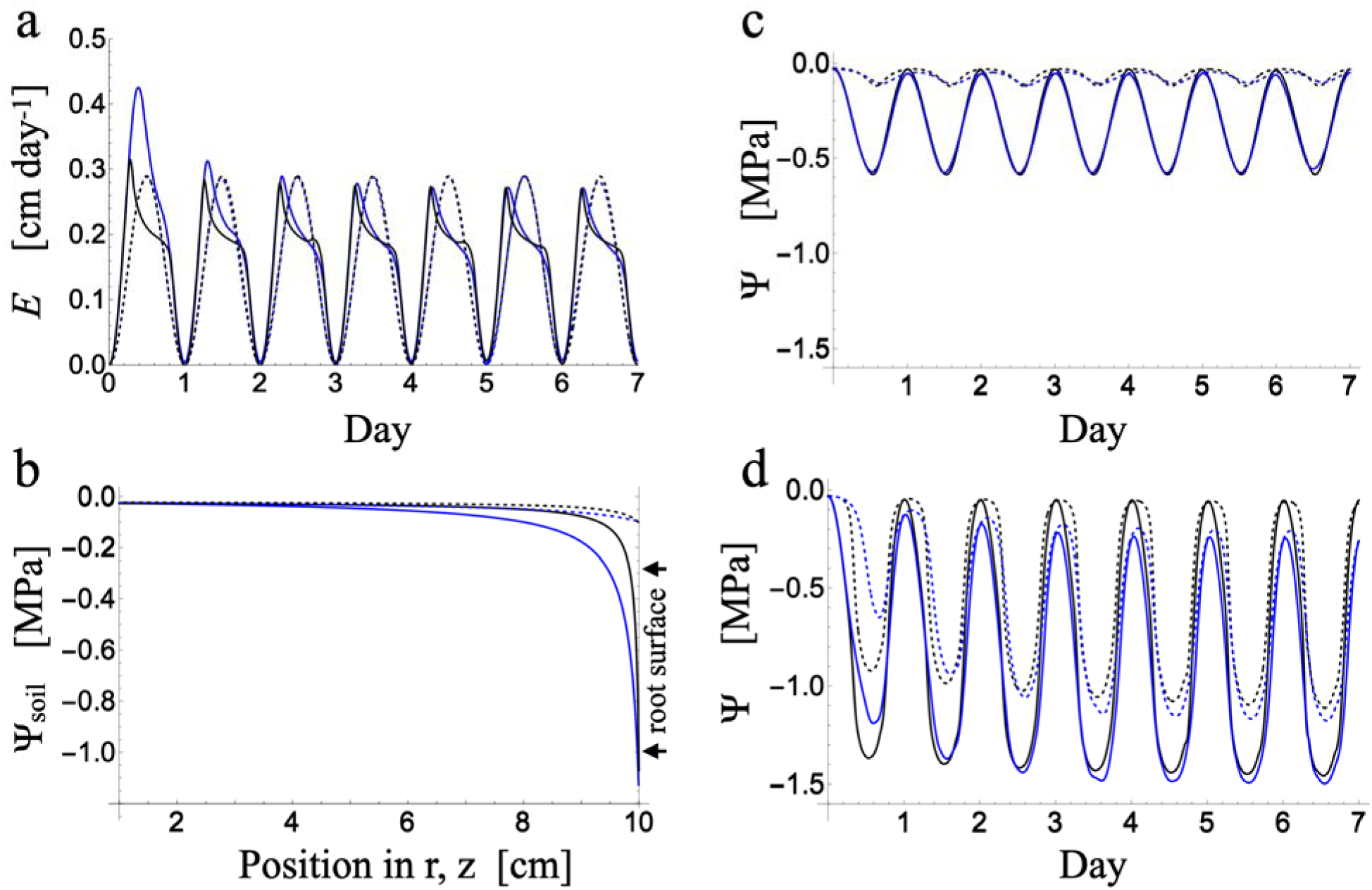
Comparison of a cylindrical soil + root domain (black) and the planar geometry adopted in this study (blue) at an *nVPD* of 1 (dashed in **a**,**b** all data in **c**) and 2 (solid in **a**,**b** all data in **d**). **a** In the cylindrical geometry feedforward reductions in *E* are slightly steeper in the a.m. but recover to slightly higher levels in the p.m., but stomata in both geometries respond similarly to the increase in evaporative demand at *nVPD* 2. **b** In the cylindrical domain the gradients start out shallower than in the planar geometry, but become steeper as the flux concentrates in the approach to the root surface, an effect more easily seen at *nVPD* 2 (solid) than 1 (dashed). Despite the difference in the steepness of the gradients, the daily minima of leaf (solid) and root (dashed) water potentials are broadly similar between geometries at both an *nVPD* of 1 (**c**) and 2 (**d**), although under higher diurnal demand overnight recovery is greater in the cylindrical case (**d)**.

