## Supplemental Notes S1 for "Emergent feedforward and isohydric responses to soil and atmospheric aridity: Insights from a time-dependent hydraulic model"

**Supporting Information Notes S1**: Discussion and derivation of governing equations and boundary conditions.

**Author**: Fulton E. Rockwell

**Article acceptance date**:

### 1.1 Derivation of an asymptotic 'maximum' soil flux from a saturated source to an evaporative surface or uptake plane

Jury, Gardner and Gardner (1991) describe the evolution of a steady upward flow to an evaporative surface in response to steady evaporative demand. They show that this flux tends to a maximum in the limit of infinite evaporative demand that depends on the soil hydraulic conductivity function and the depth from the evaporative surface to the water table. The soil hydraulic conductivity $K$ [cm day$^{-1}$] and the flux $J$ [cm day$^{-1}$] are taken to be functions of the soil matric potential $h$ (cm) that represents the capillary forces drawing water toward the evaporative surface. A suitable form for the conductivity function is,

$$K(h) = \frac{K_s}{1 + (1 + h/a)^n}, \tag{1}$$

where $K_s$ is the saturated hydraulic conductivity, and $a, n$ are the constant shape parameters of the function. The flux takes the Buckingham-Darcy form,

$$J = -K(h)\left(\frac{dh}{dz} + 1\right), \tag{2}$$

where the addition of 1 to the potential gradient results from the inclusion of gravitational effects on the flow. Combining these two equations and separating $h$ and $z$ is followed by integration of the $dh$ term from $h = 0$ at the water table to $h = -\infty$ at the evaporative surface, and integration of the $dz$ term over the height difference $L$, with the result that height is related to the maximum flux ($J_m$) in the limit of an infinitely dry surface as,

$$L = \int_{-\infty}^{0} \frac{dh}{1 + (J_m/K_s)\left(1 + (h/a)^n\right)}. \tag{3}$$

Jury *et al.* (1991) find tabulated solutions to approximations of the above integral that work well for very large (many meters) or very small (less than 10 cm) $L$, as no general solution appears to exist for the full problem. However, for specific integer values of $n$, Mathematica (Wolfram Inc, USA) does find solutions to the full problem. Solutions for other values of $n$ can be calculated by numerical inte-

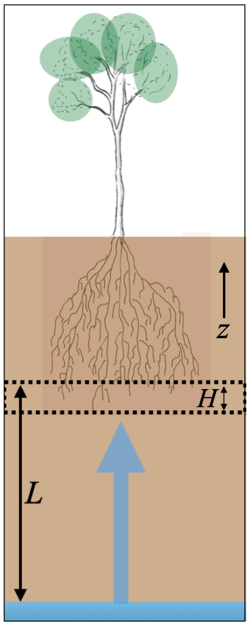


Figure 1 Daily discharge due to transpiration and recharge from an asymptotic flux from a deep soil reservoir, located a distance $L$ from a plane of root uptake, is modeled in a soil domain of height $H$.

gration. For the common values for soils of $n$ equal to two or three,

$$L = \frac{aK_s\pi}{2J_m}\left(K_s/J_m + 1\right)^{-1/2} \qquad n = 2, \tag{4}$$

$$L = \frac{2a^3K_s\pi}{3\sqrt{3}J_m}\left(-a^3\left(K_s/J_m + 1\right)\right)^{-2/3} \qquad n = 3. \tag{5}$$

The above solutions describe the maximum flux that can occur over a distance $L$ from the water table to an evaporative surface (Supplemental Information Figs. **S1**, **S2**). Note that this is a limiting behavior in which the water flux becomes asymptotic, tending toward a maximum it never quite reaches no matter how large the force driving the flux becomes. While this may be a reasonable limit for the the soil surface exposed to the dryness of air (on the order of -50 MPa at 70 RH), this idea may not be applicable to an uptake surface in the soil at which roots absorb the soil water flux, as root water potentials are definitely finite. Integrating eq (3) between zero matric potential and a finite matric potential provides an equation for plotting the flux $J$ as a function of sink matric potential $h_r$ and depth $L$, to check how dry the uptake surface has to become for $J$ to approach its maximum $J_m$. For the case that n=2,

$$L = \frac{aK_s \arctan\left[\frac{h_r}{a\sqrt{K_s/J+1}}\right]}{J\sqrt{K_s/J+1}}. \tag{6}$$

This solution is plotted in Supplemental Information Fig. **S3**. For other values of $n$ numerical integration can relate specific values of $L$,$h$, and $K_s$ to $J$.

#### 1.1.1 Unsaturated sources of soil moisture

Just as the soil flux integral in eq. 3 can be evaluated for a finite rather than infinitely dry sink, the source need not be saturated at a matric potential of zero (water table). The tendency toward an asymptotic maximum flux is a generic feature of media that lose conductivity as the driving force for transport increases, and will occur for any sufficiently large finite difference in the source and sink potentials that specify the limits of integration (Sperry & Love, 2015; Carminati & Javaux, 2020). In the case of a non-zero source at matric potential $h_s$ with $n = 2$, the solution to eq. 3 is just the sum of the finite and infinite limit solutions above,

$$L = \frac{aK_s\pi}{2J_m}\left(K_s/J_m + 1\right)^{-1/2} + \frac{aK_s \arctan\left[\frac{h_s}{a\sqrt{K_s/J_m+1}}\right]}{J_m\sqrt{K_s/J_m+1}} \qquad n = 2. \tag{7}$$

Similarly, the solution from a non-zero source to a finite sink is given by a combination of the two finite limit solutions:

$$L = \frac{aK_s\left(\arctan\left[\frac{h_s}{a\sqrt{K_s/J+1}}\right] - \arctan\left[\frac{h_r}{a\sqrt{K_s/J+1}}\right]\right)}{J\sqrt{K_s/J+1}}. \tag{8}$$

This solution is shown for $L = 100$ cm in Chino clay in Supplemental Information Fig. **S4**. The solutions for $n = 3$ involve many terms and are more easily numerically integrated, as for other values of $n$.

### 1.2 A time-dependent minimal SPAC model of transpiration as a 'null' model for stomatal responses to hydraulic supply and demand

While stomata are undeniable biologically regulated, their importance as a controlling resistance in the SPAC makes their inclusion in even a minimal purely hydraulic model essential (Van Den Honert 1948; Jarvis & McNaughton 1986). In doing so, a distinction between 'passive' (physical) stomatal responses and 'active' biological responses can be drawn. Over long (non-physiological) time scales, the average water potential-stomatal conductance relationship generally follows a monotonic decline of gs in response to water stress (Brodribb & Holbrook 2003; Anderegg *et al.*, 2017; Levin *et al.*, 2019). Stomatal closure in this case may be regarded as thermodynamically 'passive' in the sense that the response does not require the plant to do further work. With this framing, examples of 'active' or 'biological responses' include changes in the stomatal response curve to local water potential by osmotic adjustment (e.g., a shift in the water potential for 50% closure gs$\psi$50, or, over longer timescales, changes in root to shoot ratio), or deviations from the curve to satisfy a biological constraint (e.g., conservation of water use efficiency, internal $CO_2$ concentration). The active vs passive distinction here does not follow the definition of Brodribb & McAdam (2011), centered on ABA responses, but is meant to distinguish between stomatal behaviors that can be adequately captured by a single integrative variable - local water potential - versus behaviors requiring additional parameterizations. A minimal hydraulic model then is one that assembles a set of functions that are all passively driven by water potential.

#### 1.2.1 The interplay of supply and demand functions in limiting transpiration

Dynamic losses of conductivity somewhere along the SPAC have the potential to amplify stomatal sensitivity to atmospheric demand by increasing the hydraulic feedback of supply limitation on the leaf water potentials constraining stomatal conductance. The general behavior emerging from dynamic losses of conductivity – conductivity that declines as the local potential becomes more negative – is for the flux to approach a maximum in the limit of an infinite driving force, as in the steady-state analyses of Gardner in (1.1). Two principle sites for such dynamic losses have been proposed: within stem xylem (Sperry & Love, 2015), or in the soil proximal to the root (i.e., rhizosphere; Cowan, 1965; Carminati & Javaux, 2020). Dynamic losses of conductivity could also arise within leaves through minor vein collapse, or across the membranes of bundle sheath or mesophyll cells (i.e., outside xylem; Shatil-Cohen *et al.*, 2011; Zhang *et al.*, 2016; Scoffoni *et al.*, 2017; Jain *et al.*, 2024a,b). Conduit collapse may play a role in reversing transient wrong-way responses that are resolved well-within the time scale of interest for SPAC models that span hours to days, and so are neglected here (Zhang *et al.*, 2016). The losses of leaf mesophyll conductivity, attributed to down-regulation of plasma membrane conductivity, is a biological response that falls outside the scope of a physical null model. That is, failure of a purely physical model to capture observed stomatal behavior would point toward the necessity of accounting for such effects.

To address the question of whether soil or xylem conductivity losses are more likely to limit transpiration, Sperry *et al.*, (1998) modeled water transport through a discretized soil-plant continuum, concluding that xylem vulnerability was likely to be the dominant dynamic factor contributing to stomatal regulation. Carminati & Javaux (2020) took a similar approach to Sperry & Love (2015), finding that with their parameterization soil conductivity emerged as the factor limiting transpiration. In addition to losses of hydraulic conductivity in the soil itself, root-soil contact has emerged as another source of variable resistance that might explain apparent soil resistances that exceed the level predicted by root density and soil properties (Herkelrath *et al.*, 1977). Drying of root mucilage or root shrinkage with the onset of soil water stress could lead to an increase in air gaps between the soil and roots (McCully *et al.*, 1997), with liquid flow across the contact zone increasingly replaced by relatively inefficient water vapor diffusion, creating a bottle neck (Carminati *et al.*, 2009, 2016; Rodriguez-Dominguez & Brodribb, 2020; Manandhar *et al.*, 2024; Akale *et al.*, 2025).

Both the Sperry & Love and Carminati & Javaux analyses were limited to a steady-state perspective. Cowan (1965) modeled time-dependent behavior of E in drying finite volumes of soils (pots) to show that dynamic declines in soil water content and hydraulic conductivity proximal to the root could constrain E during the day, followed by recovery overnight, even as bulk soil moisture declined monotonically over multiple days (Fig. 11 in Cowan, 1965). In this finite-volume time dependent case, the maximum soil flux that emerges is not fixed as in the steady state analyses of Sperry & Love and Carminati & Javaux, but tracks the minimum soil water potential at the root surface in decaying over the day and partly recovering over night, with the daily average maximum flux decaying over days monotonically as total soil moisture in the finite volume similarly declines. In all these studies, stomatal conductance was not explicitly modeled so the relationship between dynamic conductivity and realized stomatal behavior remained unexplored. Instead, a critical leaf water potential for stomatal closure was defined that effectively served as a transition point from perfect anisohydry to perfect isohydry (Cowan, 1965; Hochberg *et al.*, 2018; Javaux & Carminati, 2021).

Tuzet *et al.*, (2003) constructed the most complete model of the SPAC to date, as far as the author is aware. This model follows Cowan (1965) in modeling time dependent soil moisture in a cylinder of soil around a representative root within a finite soil volume, but adds stomatal conductance as a logistic function of leaf water potential (with no explicit dependence on VPD) and additional forcing from internal CO2 concentrations and a photosynthesis model, driven by diurnally varying radiation and surface energy balance with a boundary layer. This model generated both isohydric and anisohydric behavior, and feed-forward stomatal closure that was suggested to be driven by the soil. However, the complexity of the model prevented fully accounting for the generated behavior, and, as acknowledged in Tuzet et al. (2003), they were unable to locate a sufficiently parameter-rich data set whose behavior they could attempt to reproduce and explain. This trade-off between complexity and applicability provides further motivation for a minimal physical model.

#### 1.2.2 Transient models of unbounded soil volumes

According to Tuzet *et al.* (2003), most models of root uptake from soils follow Philip (1957) and Cowan (1965) in modeling a representative root at the center of a single cylindrical soil layer. As the angular dependence is neglected, the flow depends only on a single dimension, the radial coordinate $r$. The important feature of this description is that the conserved quantity at steady-state in the soil domain is not the flux $J$, but the flux times the radial coordinate $Jr$, as the surface area over which the flux is spread declines in the direction toward the root. This constriction in the area over which flow occurs creates an additional spatially varying effective resistance in addition to the hydraulic conductivity of the soil that varies with soil water (or matric) potential. In this approach, the radius of the soil domain is determined by the root length per soil volume, such that the outer boundary of the cylinder represents the midpoint in the distance between two roots (Cowan, 1965), and flow is studied in a representative cross section between the outer boundary of the annular soil domain and the inner boundary of the root surface. This 'root cylinder' model seems particularly appropriate for potted plants where roots explore a well-defined finite volume. While Cowan (1965) does not provide explicit criteria for applying the root cylinder concept to less well spatially defined root zones as they exist in nature, it would seem a reasonable approximation as long as hydrological flows into the root zone from other parts of the soil remain remain negligible relative to transpiration.

Yet outside the case of time-dependent modeling of finite (pots) and relatively isolated soil domains, the root cylinder approach encounters both logical and practical difficulties. This is because the natural boundary condition at the outer edge of a the soil domain around a root is a no-flux or 'insulated' condition. On either side of an infinitely thin boundary between two root cylinders, flow occurs away from the boundary in opposite directions toward the two root-sinks: there can be no flow *across* the boundary. Yet the logical requirement for a no-flux condition on the outer boundary makes the cylindrical root+soil domain model inappropriate for steady-state modeling, as the only possible solution at steady state is zero flux. To put it another way, the only steady-state attainable in the transient problem with zero inward flux on the boundary is one in which soil moisture is completely depleted and the flux is zero. To get around this limitation preventing application of the root-cylinder model to steady-state, practitioners have simply fixed soil moisture (or potential flux) at the boundary, making the infinitely thin boundary between root cylinders a source for a steady-state flux (Sperry *et al.*, 1998); Carminati & Javaux, 2020). The non-physical nature of this condition means that the gradient of potential within the soil cylinder is never actually attained, it is simply the gradient that would obtain if one could have a water source at the boundary between two roots. Despite this artificiality, the steady-state root cylinder model has a certain theoretical or pedantic utility for understanding the general behavior of soil water potential gradients near the root surface (Cowan, 1965), and may be a reasonable approximation when the effective depth of an external source is much larger than the thickness of the root zone.

While modeling the soil as an isolated and finite volume composed of root-soil cylinders can be a reasonable approximation when soils are near field capacity (*i.e.*, the inward soil flux is small relative to transpiration), or the outer radius of the soil cylinder is set to be much smaller than that implied by root density, drying of the soil system ultimately breaks these approximations. The most obvious issue is that as the root zone dries down, deep lateral flows driven by topography, groundwater, or simply residual moisture in the unsaturated soil outside the root zone will generally provide sources of soil moisture that will at some rate flow toward the moisture depleted root zone (Fan *et al.*, 2017; Miquez-macho & Fan 2021). Such sources can simply be localized to the root cylinder boundary (see Section 1.3.6) if one is willing to neglect the unknown resistance to distributing the flux from the lower root zone boundary to all the root surfaces.

Here, we propose to resolve these difficulties to find an internally consistent treatment of root uptake through further abstraction: root uptake of water within the rooting volume is projected down to a rooting-plane or uptake surface (Fig. 1 located at some characteristic rooting depth (Binks *et al.*, 2022). A root plane approach captures neither the internal resistance of a root zone, nor the geometrical effects of a cylindrical domain, yet it is an idealization that is consistent with the overall level of geometrical abstraction in SPAC models (Philip, 1966) as well as with hydrologic concepts of a 1D effective root depth at scale (Fan *et al.*, 2017). A rooting surface may also be more physically consistent than a rooting volume in the case that transpiration depends on a flux of water from below the volume of soil explored by roots. When the root zone itself is the source of moisture for transpiration, the cylindrical soil and root domain models capture the effect of the full surface area of roots per unit volume of soil, and root to shoot ratio is a critical parameter governing the balance of resistance between the soil and plant. Yet, once transpiration depends on an external flux into the root zone from deeper soil, total root surface area may not matter. When uptake is dominated by the roots in contact with the wettest soil region (Dawson *et al.*, 2020) the limiting surface area may become the projected area of the root domain across which the flux from deeper sources of moisture occurs. A root plane approximation appears more suited to natural systems for which root competition has created an overall high root density than for agricultural settings such as recently established annual crops. Ultimately, the root plane concept represents the hypothesis that, given the expected behavior of soil fluxes becoming asymptotic as the system dries (Cowan, 1965; Jury *et al.*, 1991) neglecting cylindrical resistance will not qualitatively change the behavior of the results.

### 1.3 Transient model for a soil domain fed by an inward maximum flux from the water table.

To collapse the 3D flux water through the soil to a 1D SPAC model, we consider a soil domain that is homogenous in the neglected directions $(x, y)$ parallel to the surface and plant canopy with the only gradients in $z$ (Fig. 1). The movement of water is expressed as a flux density (mmol m$^{-2}$ s$^{-1}$, or equivalently cm$^3$ m$^{-2}$ s$^{-1}$= cm s$^{-1}$). The upper surface of the explicitly modeled soil domain at $z = H$ is assumed to represent a plant root uptake 'plane.' The lower surface of this domain is fed by a steady asymptotic soil flux that is a function of the distance $L$ from a deep saturated water source to the rooting plane (Section 1.1). The outward flux at the rooting plane (i.e., absorption by the plant) is driven by a time-varying function that describes the diel trends in both $VPD$ and canopy conductance. This transient function multiplies a dimensionless canopy conductance $g$ that describes the path from the leaf internal airspace to the well mixed air above the canopy where VPD is measured, and a dimensionless normalized driving force $nVPD$. Note that $g$ is a function of leaf water potential through its partial dependence on stomatal conductance $g_s$. Note too that $g_a$ or the canopy aerodynamic (or boundary layer) conductance is fixed at a characteristic peak value, that may be regarded as varying over the daily cycle as morning calm gives way to afternoon convection that then relaxes to calm conditions again overnight. Finally, the product of all the dimensionless quantities above multiplies a potential maximum transpiration rate $E_p$ in the velocity units of cm day$^{-1}$.

The height of the soil domain is set by the distance required for daily variations in soil moisture to damp out, $H > \sqrt{2Dt} \approx 10$ cm (Cowan, 1965), small enough that we neglect gravitational effects on soil water potential within the domain. The governing equation for the domain and boundary conditions that relates changes in water content ($\theta$ in cm$^3$/cm$^3$) to the flux of water then take the forms:

$$\frac{\partial \theta}{\partial t} = \frac{\partial}{\partial z}\left(K(h)\frac{\partial h}{\partial z}\right) \tag{9}$$

$$-K(h)\frac{\partial h}{\partial z}\bigg|_{z=0} = J_m(L), \tag{10}$$

$$-K(h)\frac{\partial h}{\partial z}\bigg|_{z=H} = E_p\, g(\psi_l)\; nVPD\, \left(\frac{1 - Cos\frac{2\pi t}{\tau}}{2}\right), \tag{11}$$

where $\tau$ is the period of oscillation, one day. Following Cowan (1965), plant capacitance and gravity (less than 0.07 MPa for trees less than 7 meters) are neglected,

$$\psi_l = \psi_{root}(h|_{z=H}) + \frac{K(h)\frac{\partial h}{\partial z}\big|_{z=H}}{K_p}. \tag{12}$$

Stomatal conductance is modeled as a sigmoid function of water potential,

$$g(\psi_l) = \left(\frac{1}{g_s(\psi_l)} + \frac{1}{g_a}\right)^{-1}, \qquad g_s = \left(1 + \exp\left(\frac{\psi 50 - \psi_l}{b}\right)\right)^{-1}, \tag{13}$$

where $\psi 50$ is the water potential at 50% stomatal closure, and $b$ is a shape parameter. Finally, to express all the driving forces in a common unit, in standard SI units plant water potentials $\psi$ (Pa) are converted to matric potential (m) using the relation,

$$h = \frac{\psi}{\rho g}. \tag{14}$$

#### 1.3.1 Linearization of the governing equation (Kirchhoff transform).

Solution of the above set of equations is simplified by partially linearizing the governing equation eq (9) using the Kirchhoff transform (Peavy, 1996), which implements a change of variables to a potential flux $\phi$ (cm$^2$ day$^{-1}$) that integrates the conductivity function from $-\infty$ up to a local value of $h$:

$$\phi = \int_{-\infty}^{h} K(h)\ \mathrm{dh} \tag{15}$$

The governing equation in terms of $\phi$, with the definition of the diffusivity $D$, become,

$$\frac{\partial \phi}{\partial t} = D(\phi)\frac{\partial^2 \phi}{\partial z^2} \tag{16}$$

$$D(\phi) = \left(\frac{\partial \theta}{\partial \phi}\right)^{-1} \tag{17}$$

Finding the function $D(\phi)$ as well as transformation of the boundary conditions requires specification of a particular set of soil characteristics prescribing $K(h)$ and $h(\theta)$, which determine the relation between the potential flux $\phi$ and volumetric water content $\theta$. Cowan (1965) adopts approximate exponential fits to data for Yolo light clay given by Philip (1957) over a range of soil moisture from $\theta = 0.25$ to 0.1, which aids solution of the system. As we have no particular knowledge of the soil in Drake *et al.* (2018), for modeling that experiment we adopt this same parameterization to define the set of relations:

$$K(\theta) = 6.03 \times 10^{-9}\ \exp(51.4\theta) \tag{18}$$
$$h(\theta) = -2.67 \times 10^{5} \exp(-25.7\theta) \tag{19}$$
$$\phi(\theta) = 0.0016\ \exp(25.7\theta) \tag{20}$$
$$D(\theta) = 0.0414\ \exp(25.7\theta) \tag{21}$$
$$D(\phi) = 25.7\ \phi \tag{22}$$
$$h(\phi) = \frac{-430.11}{\phi}. \tag{23}$$

With the above, the boundary conditions are re-cast as,

$$-\left.\frac{\partial\phi}{\partial z}\right|_{z=0} = J_m(L), \tag{24}$$
$$-\left.\frac{\partial\phi}{\partial z}\right|_{z=H} = E_p\ g(\phi)\ nVPD\ \left(\frac{1 - Cos\frac{2\pi t}{\tau}}{2}\right). \tag{25}$$

The transformed stomatal conductance contributing to $g(\phi)$ is given by,

$$g_s(\phi) = \left(1 + \exp\left(\frac{-430.11\rho g\left(\phi 50^{-1} - \phi^{-1}\right)}{b}\right)\right)^{-1} \tag{26}$$

To initialize the transient model we arbitrarily set the potential flux to 1, corresponding to $\theta = 0.25$ at $T = 0$ (near field capacity), and evolve the system toward a quasi-steady diurnal cycle of soil drying, as transient transpiration extracts water, and re-wetting due to an inward steady (maximum) flux from the water table.

$$\phi(T = 0, z) = 1. \tag{27}$$

Table 1 Parameter definitions and values

| Quantity | Symbol | Units | Value |
|---|---|---|---|
| Distance from water table to root zone | $L$ | $m$ | - |
| Water potential to matric conversion factor | $(\rho g)^{-1}$ | cm MPa$^{-1}$ | $1.0227\times10^{4}$ |
| Depth of model domain | H | cm | 10 |
| Shape factor ($g_s$) | $b$ | MPa | 0.23 |
| Water potential at 50% closure $g_s$ | $\psi 50$ | MPa | -1.33 |

#### 1.3.2 Dimensional analysis

To analyze the behavior of the governing equation and boundary conditions, we non-dimensionalize the variables:

$$z = L_c Z, \qquad t = t_c T, \qquad \phi = \phi_c \Phi, \tag{28}$$

where the subscript c denotes a quantity characteristic of the system by which we normalize the variables so that their expected values are on the order of 1. For the characteristic length we choose the depth of the model domain $H$. The governing equation and boundary conditions then become,

$$\frac{\partial \Phi}{\partial T} = \frac{t_c \phi_c D_o}{H^2} \Phi \frac{\partial^2 \Phi}{\partial Z^2}, \quad \rightarrow \quad D(\Phi) = D_o \Phi, \quad D_o = 25.7 \tag{29}$$

$$-\left.\frac{\partial \Phi}{\partial Z}\right|_{Z=0} = \frac{H J_m}{\phi_c}, \tag{30}$$

$$-\left.\frac{\partial \Phi}{\partial Z}\right|_{Z=1} = \frac{H E_p\, nVPD}{\phi_c} g(\phi) \left( \frac{1 - Cos \frac{2\pi t_c T}{\tau_d}}{2} \right). \tag{31}$$

With the choice $\phi_c = H J_m$ for the characteristic potential flux, and the period of oscillation of one day for the characteristic time scale, $t_c = \tau_d$, the fully non-dimensionalized set of equations becomes:

$$\frac{\partial \Phi}{\partial T} = \frac{\tau_d J_m D_o}{H} \Phi \frac{\partial^2 \Phi}{\partial Z^2}, \quad \rightarrow \quad D(\Phi) = D_o \Phi, \quad D_o = 25.7 \tag{32}$$

$$-\left.\frac{\partial \Phi}{\partial Z}\right|_{Z=0} = 1, \tag{33}$$

$$-\left.\frac{\partial \Phi}{\partial Z}\right|_{Z=1} = \frac{E_p\, nVPD}{2 J_m} g(\phi) \left(1 - Cos 2\pi T\right). \tag{34}$$

This last boundary condition for the plant flux defines $nVPD$. We define a 'stress-free' initial state as a series of days over which midday $VPD$ and transpiration $E_p$ are fairly constant, which by mass conservation implies that $E_p = 2J_m$. The expected value of $1 - Cos 2\pi T$ over a day is one, as is the stress free value of $g$, and we take the value of $VPD$ under these conditions as a reference state and define $nVPD = VPD/VPD_{ref}$.

There are then two dimensionless parameter groups that influence the behavior of the solution.The first is a ratio of time scales, the period for oscillations of evaporation $\tau_d$ of one day, and the second the time-scale for the diffusion of the potential flux in the soil $H/(J_m D_o)$, which together govern the time to establish a diel stationary state. The second is the balance of evaporative demand to the steady maximum soil flux from the water table,

$\beta = E_p nVPD/(2J_m)$. With this, the the full problem is given by,

$$\frac{\partial \Phi}{\partial T} = \frac{\tau_d}{\tau_s}\Phi\frac{\partial^2 \Phi}{\partial Z^2}, \qquad \tau_s = \frac{H}{J_m D_o} \tag{35}$$

$$-\left.\frac{\partial \Phi}{\partial Z}\right|_{Z=0} = 1, \tag{36}$$

$$-\left.\frac{\partial \Phi}{\partial Z}\right|_{Z=1} = \beta\, g(\Phi)\,\left(1 - Cos 2\pi T\right), \qquad \beta = \frac{E_p nVPD}{2J_m}, \tag{37}$$

$$\Phi(0, Z) = \frac{1}{HJ_m}, \tag{38}$$

$$g_s\left(\Phi\right) = \left(1 + \exp\left(\frac{\frac{-430.11\rho g}{HJ_m\Phi 50} - \left(\frac{-430.11\rho g}{HJ_m\Phi|_{Z=1}} + \frac{J_m \partial\Phi/\partial Z|_{Z=1}}{K_p}\right)}{b}\right)\right)^{-1}. \tag{39}$$

Note that the factor -430.11 converting matric potential to potential flux carries units of $cm^3$ $day^{-1}$.

#### 1.3.3 Notes on parameterization and application of the model for Drake *et al.* (2018).

In the Drake *et al.* (2018) experiment, individual trees were rooted in soil with the above ground portion contained within gas exchange chambers with sufficent fan speeds that air and leaf temperatures converged; under these conditions the boundary layer can be neglected and the VPD of the well mixed air can be taken as the VPD at the leaf surface, such that $g \approx g_s$. These conditions create no loss of generality as they represent the typical gas exchange cuvette conditions under which stomatal responses to VPD are studied (Mott & Parkhurst, 1991; Franks *et al.*, 1999), or are assumed to hold at larger scales in uncontrolled environments (Grossiord *et al.*, 2020; Mencuccini *et al.*, 2024). The neglect of cavitation in the plant is further justified in this specific case by leaf and stem measurements in the Supplemental Information of Drake *et al.* (2018) that show no reduction in hydraulic conductance occurred during the experiment. As the soil properties for the Heat Wave experiment were unavailable from Drake *et al.* (2018), the hydraulic conductivity and matric potential functions of water content were parameterized based on Cowan (1965) for Yolo light clay. With this parameterization, a maximum steady soil flux of 0.15 cm day-1 develops over a distance of 50 cm between a water source and rooting plane (Supplemental Information Figure S3), similar in magnitude to the one-meter distance between the depth of greatest change in soil water content (50 cm) and no change in soil water content (150 cm) given by neutron counts for the soils in Drake *et al.* (2018). By way of comparison, for the fine sandy loam (Pachappa) and clay (Chino) soils studied by Jury *et al.* (1991), a flux of 0.15 cm $day^{-1}$ occurs over a distance of 130 and 155 cm respectively (Supplemental Information Fig. S2). These distances support a hypothesis that in the physical experiment soil properties and distance from source to sink on the order of a meter could have imposed the observed daily maximum flux of 0.15 cm $day^{-1}$. Finally, to initialize the model soil water content in the model domain was set to a uniform value $\Theta = 0.25$ ($i.e., \phi = 1$), about half field capacity (Philip, 1957).

##### 1.3.4 Stomatal sensitivity to VPD in a diel stationary-state

We consider a system in a diel stationary state for which $g_a \gg g_s$. Mass conservation requires that soil supply (24) and atmospheric demand (25) balance over a 24 hour period,

$$J_m \approx \frac{E_p\ g_{s,c}\ nVPD}{2}. \tag{40}$$

Here $g_{s,c}$ refers to the average or characteristic value of stomatal conductance over the diurnal steady-state. With $VPD_c$ the peak value of $VPD$ over the period, re-arranging the above leads to an expression for the non-dimensional stomatal conductance as,

$$g_{s,c} \approx \frac{2J_m}{E_p\ nVPD}, \qquad nVPD = \frac{VPD}{VPD_c}. \tag{41}$$

Operationally, (41) can be put to use by identifying a period over which $J_m$ is expected or known to be steady, and choosing a characteristic time for observation of the relationship between conductance and atmospheric dryness, such as midday. Observing this system at some low $VPD_{ref}$ defines a reference state:

$$g_{s,ref} \approx \frac{2J_m}{E_p\ nVPD_{ref}}, \qquad nVPD_{ref} = \frac{VPD_{ref}}{VPD_c} \equiv 1. \tag{42}$$

Normalizing by this reference state leads to a general form for comparing responses across environments and species,

$$\frac{g_{s,c}}{g_{s,ref}} \approx \frac{2J_m}{E_p\ g_{s,ref} nVPD} \equiv \frac{1}{nVPD}. \tag{43}$$

For the purposes of fitting the simulated diurnal stomatal conductances in Figure 7, we write,

$$g_{s,c} \approx \frac{g_{s,ref}}{nVPD}, \tag{44}$$

where $g_{s,ref}$ is averaged over the 'observed' values of $g_s$ at $nVPD = 1$ across all simulations. Note that the expected value based on the steady-state equivalence of total daily supply and demand in (42) is 1, but diurnal hysteresis of $g_s$ with $nVPD$ will lead to fits less than 1.

More generally, for any pair of conductances and driving forces determining transpiration $J(g, \chi)$, and where $(\bar{g}, \bar{\chi}, \bar{J})$ represent averages taken over a period for which the soil flux and transpiration are believed to be in balance (e.g., instantaneously at midday for physiological time scales, or over a 24 hour period for diurnal stationary states), mass balance (40) leads to the following expression and reference state,

$$\bar{g} \approx \frac{\bar{J}}{\bar{\chi}}, \qquad \bar{g}_{ref} \approx \frac{\bar{J}_{ref}}{\bar{\chi}_{ref}} \tag{45}$$

And the decay of normalized conductance is described by,

$$\frac{\bar{g}}{\bar{g}_{ref}} \approx \left(\frac{\bar{J}}{\bar{J}_{ref}}\right)\left(\frac{\bar{\chi}_{ref}}{\bar{\chi}}\right). \tag{46}$$

This form shows clearly how the soil flux controls the sensitivity of conductance to atmospheric aridity.

#### 1.3.5 Special case of zero inward flux: The root zone as a finite volume

In well-drained soils with frequent rainfall, root growth may remain relatively shallow (Fan *et al.*, 2017; Miquez-macho & Fan 2021). Under drought, if the flux from deep sources of soil moisture is negligible (e.g., hill tops) then the domain of soil moisture available to plants may be well-approximated as a finite volume (root zone) within which bulk water content decays monotonically over time (Cowan 1965; Tuzet *et al.*, 2003). We again consider the general case with an inward flux at the lower boundary,

$$\frac{\partial \Phi}{\partial T} = \frac{t_c \phi_c D_o}{H^2} \Phi \frac{\partial^2 \Phi}{\partial Z^2}, \quad \rightarrow \quad D(\Phi) = D_o \Phi, \quad D_o = 25.7 \tag{47}$$

$$- \left. \frac{\partial \Phi}{\partial Z} \right|_{Z=0} = \frac{H J_m}{\phi_c}, \tag{48}$$

$$- \left. \frac{\partial \Phi}{\partial Z} \right|_{Z=1} = \frac{H E_p \, nVPD}{\phi_c} g(\phi) \left( \frac{1 - Cos \frac{2\pi t_c T}{\tau_d}}{2} \right). \tag{49}$$

We again choose $t_c = \tau_d$ as a time scale. For the characteristic potential flux $\phi_c$, as an alternative scale to the maximum soil flux we choose the transpiration flux, $H \, E_p \, nVPD/2$. With these choices,

$$\frac{\partial \Phi}{\partial T} = \frac{\tau_d E_p \, nVPD \, D_o}{2H} \Phi \frac{\partial^2 \Phi}{\partial Z^2}, \quad \rightarrow \quad D(\Phi) = D_o \Phi, \quad D_o = 25.7 \tag{50}$$

$$- \left. \frac{\partial \Phi}{\partial Z} \right|_{Z=0} = \frac{2 J_m}{E_p \, nVPD}, \tag{51}$$

$$- \left. \frac{\partial \Phi}{\partial Z} \right|_{Z=1} = g(\Phi) \, (1 - Cos 2\pi T) \, . \tag{52}$$

Again, the factor of 2 is included in $\phi_c$ such that the time varying term $1 - Cos 2\pi T$ varies from 0 to 1. The condition for treating the soil domain as an isolated finite volume is than given by the lower boundary condition, which goes to zero in the limit $E_p \, nVPD \gg 2J_m$. Again, the behavior of the system is controlled by the ratio of two time scales, daily oscillations in evaporative demand versus the time for the effect of atmospheric demand to invade the soil,

$$\frac{\partial \Phi}{\partial T} = \frac{\tau_d}{\tau_s} \Phi \frac{\partial^2 \Phi}{\partial Z^2}, \qquad \tau_s = \frac{2H}{E_p \, nVPD \, D_o} \tag{53}$$

$$- \left. \frac{\partial \Phi}{\partial Z} \right|_{Z=0} = 0 \qquad (\text{for } E_p \, nVPD \gg 2J_m) \, , \tag{54}$$

$$- \left. \frac{\partial \Phi}{\partial Z} \right|_{Z=1} = g(\Phi) \, (1 - Cos 2\pi T) \, . \tag{55}$$

The invasion time $\tau_d$ will be fast for high evaporative demand and high soil diffusivities (high conductivity/low capacity), leading to large diurnal variations relative to the size of the soil domain; for low evaporative demand and low conductivity/high capacity soils the intrusion of diurnal variations into the soil domain from the rooting plane will be relatively damped and simply track the monotonic decline of the bulk soil water content over time.

The size of that domain can again be set by according to Cowan (1965) as $H = \sqrt{2Dt}$ (for the Drake experiment parameterization ≈ 7cm), or if absorptive fine root length per unit soil volume $L_R$ (cm/cm$^3$) and the thickness of root zone $T_S$ (cm) as well as fine root diameter $r_R$ are all known, and the radius of the soil cylinder around the root is estimated as $r_S = (\pi L_R)^{-1/2}$, then an equivalent cartesian thickness for the same flux-force relation is,

$$H = \frac{\ln (r_R/r_S)}{2\pi L_R T_S}. \tag{56}$$

This relation follows from the steady-state expressions for the conservative flux for cylindrical (radial coordinate $r$ ) and slab (cartesian $z$ coordinate) geometries (Peavey, 1996). Cast in terms of the soil matric potential $h$,

$$q_c = qr = \frac{-1}{\ln (r_R/r_S)} \int_{h_R}^{h_S} k(h)\ \mathrm{dh} \tag{57}$$

$$q_c = \frac{-1}{H} \int_{h_R}^{h_S} k(h)\ \mathrm{dh} \qquad \mathrm{H = \Delta z = z_R - z_S}. \tag{58}$$

Note that the integrations are over the conductivity function over the span of matric potentials from root to the domain boundary, and so are independent of the geometry for the same $h_r$, $h_S$ (only the spatial distribution of the potential drop from bulk soil to root differs in the solutions for the two geometries). Summing the total radial flux across all the root surface area in the the soil volume with a thickness $T_S$ and normalizing to the per unit land surface area leads to a flux density that arises directly in the slab geometry:

$$\frac{-1}{H} \int_{h_R}^{h_S} k(h)\ \mathrm{dh} = \frac{\mathrm{-2\pi L_R T_S}}{\mathrm{\ln (r_R/r_S)}} \int_{\mathrm{h_R}}^{\mathrm{h_S}} \mathrm{k(h)\ dh} \tag{59}$$

##### 1.3.6 Annular soil domain with a converging radial flux toward the root and a maximum inward flux at the outer boundary

To check the effect of neglecting the effect of radial geometry of the soil around real roots in the planar SPAC model adopted here, the governing equations were recast into cylindrical coordinates. The planar boundary conditions (transpiration and maximum soil fluxes) were mapped to the outer and inner radii of the annular soil domain by employing a 'big root' model of the root zone analogous to the 'big leaf' model of plant canopies. The root zone is considered a single layer of roots, and the flux across the projected root area onto a plane at the lower boundary of the root zone is then spread around the outer radius of a representative soil and root domain (Figure 2). The flux across the inner radius of the soil domain (i.e., the root surface ) is then spread back over another plane parallel to the first and of equal area that represents the plant canopy. The original problem statement re-cast for the soil

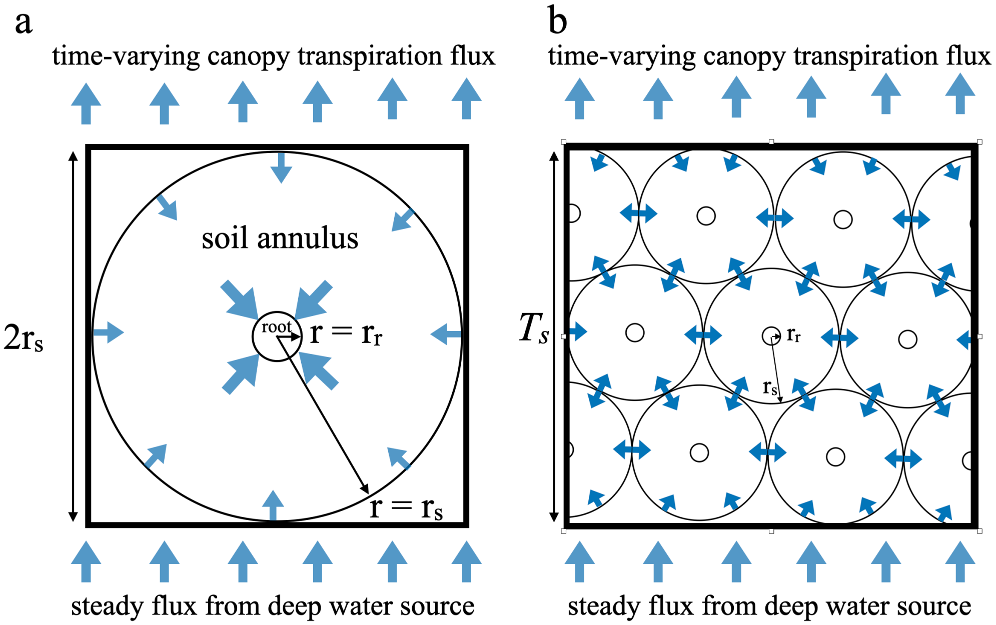


Figure 2 Cylindrical soil and root domains joined to planar fluxes of deep soil moisture from below and canopy transpiration above. The root serves a sink for an annular soil domain within which water flow is considered axisymmetric. Axisymmetry in the annular soil domain requires that the inward and outward water flux be homogenously spread over the boundaries of at $r_s$ and $r_r$, implying no resistance between the planar flux at the deepest boundary of the root zone and the entire root surface area. This true whether the root zone is idealized in a 'big root' model (a), or is described by a density of active roots $L_r$ (cm root length per cm$^{-3}$ soil volume) and a total root zone thickness $T_s$.

annulus domain becomes,

$$\frac{\partial \theta}{\partial t} = \frac{1}{r}\frac{\partial}{\partial r}\left(rK(h)\frac{\partial h}{\partial r}\right) \tag{60}$$

$$-(2\pi r_s l_r)K(h)\frac{\partial h}{\partial r}\bigg|_{r=r_s} = (2r_s l_r)J_m, \tag{61}$$

$$-(2\pi r_r l_r)K(h)\frac{\partial h}{\partial r}\bigg|_{r=r_r} = (2r_s l_r)E_p\; g(\psi_l)\; nVPD\left(\frac{1-Cos\frac{2\pi t}{\tau_d}}{2}\right), \tag{62}$$

where $l_r$ is the root length per soil volume, and the bracketed terms spread the absolute fluxes across the respective planar and cylindrical surface areas. Consolidating the geometrical terms scaling the fluxes leads to,

$$\frac{\partial \theta}{\partial t} = \frac{1}{r}\frac{\partial}{\partial r}\left(rK(h)\frac{\partial h}{\partial r}\right) \tag{63}$$

$$K(h)\frac{\partial h}{\partial r}\bigg|_{r=r_s} = \frac{J_m}{\pi}, \tag{64}$$

$$K(h)\frac{\partial h}{\partial r}\bigg|_{r=r_r} = \frac{r_s}{\pi r_r}E_p\; g(\phi_l)\; nVPD\left(\frac{1-Cos\frac{2\pi t}{\tau_d}}{2}\right), \tag{65}$$

Note that the change in sign for the boundary conditions is because we define a flux that flows in the direction of *decreasing* $r$ (toward the root) as positive. Applying the Kirchhoff transform,

$$\frac{\partial \phi}{\partial t} = \frac{D(\phi)}{r}\frac{\partial}{\partial r}\left(r\frac{\partial \phi}{\partial r}\right) \tag{66}$$

$$\left.\frac{\partial \phi}{\partial r}\right|_{r=r_s} = \frac{J_m}{\pi}, \tag{67}$$

$$\left.\frac{\partial \phi}{\partial r}\right|_{r=r_r} = \frac{r_s}{\pi r_r}E_p\; g(\phi_l)\; nVPD\; \left(\frac{1-Cos\frac{2\pi t}{\tau}}{2}\right), \tag{68}$$

Non-dimesionalizing again by adopting the re-scaled variables $\Phi = \phi/\phi_c$, $T = t/\tau_d$, $R = r/r_s$,

$$\frac{\partial \Phi}{\partial T} = \frac{\tau_d \phi_c D_o \Phi}{r_s^2 R}\frac{\partial}{\partial R}\left(R\frac{\partial \Phi}{\partial R}\right) \tag{69}$$

$$\left.\frac{\partial \Phi}{\partial R}\right|_{R=1} = \frac{r_s J_m}{\phi_c \pi}, \qquad \rightarrow \phi_c = \frac{r_s J_m}{\pi}, \tag{70}$$

$$\left.\frac{\partial \Phi}{\partial R}\right|_{R=r_r/r_s} = \frac{r_s^2}{\phi_c \pi r_r}E_p\; g(\Phi_l)\; nVPD\; \left(\frac{1-Cos2\pi T}{2}\right). \tag{71}$$

We again choose a scaling for $\phi_c$ that makes the derivative term order 1, leading to,

$$\frac{\partial \Phi}{\partial T} = \frac{\tau_d}{\tau_s}\frac{\Phi}{R}\frac{\partial}{\partial R}\left(R\frac{\partial \Phi}{\partial R}\right) \qquad \rightarrow \tau_s = \frac{\pi r_s}{J_m D_o} \tag{72}$$

$$\left.\frac{\partial \Phi}{\partial R}\right|_{R=1} = 1, \tag{73}$$

$$\left.\frac{r_r}{r_s}\frac{\partial \Phi}{\partial R}\right|_{R=r_r/r_s} = \beta\; g(\Phi_l)\; (1-Cos2\pi T) \qquad \rightarrow \beta = \frac{E_p\, nVPD}{2J_m}. \tag{74}$$

The principal effects of the change to an annular soil domain can be now be easily seen by comparing the non-dimensionalized forms of the governing equation and boundary conditions to those for the cartesian (planar) form (Section 1.3.2). $\tau_d$ and $\beta$ are unchanged, while $\tau_s$ is faster by a factor of $\pi$ (for $H \approx r_s$). The important difference is in the boundary condition at the root surface (eq.80). The RHS is unchanged, with an expected value of 1, but the LHS side derivative term is scaled by $r_r/r_s \ll 1$. As the LHS must balance the RHS, and so also have an expected value of 1, this requires that the derivative $\partial\Phi/\partial R|_{R=r_r/r_s}$ greatly exceed 1. This means that in the radial case, the gradients at the root surface are steeper than those of the cartesian case by a factor of $r_s/r_r$ for the same flux and overall water potential drop. The behavior of the two solutions respond to the value of $\beta$ in the same fashion, although the feedforward behavior that emerges in the radial case falls initially more steeply before recovering to a higher value in the afternoon (Supplemental Information Fig. S7).

The 'big root' set of equations also hold exactly for the case of a soil domain defined by a root length per unit volume of soil $L_r$ and root zone thickness $T_s$ (Fig.2b). Replacing total root length $l_c$ with $L_r T_s$ re-scales the total flux across the surfaces of the annular domain to the planar geometry of the deep soil and transpiration fluxes,

$$\frac{\partial \theta}{\partial t} = \frac{1}{r}\frac{\partial}{\partial r}\left(rK(h)\frac{\partial h}{\partial r}\right) \tag{75}$$

$$-(2\pi r_s L_r T_s)K(h)\frac{\partial h}{\partial r}\bigg|_{r=r_s} = J_m, \tag{76}$$

$$-(2\pi r_r L_r T_s)K(h)\frac{\partial h}{\partial r}\bigg|_{r=r_r} = E_p\, g(\psi_l)\, nVPD\left(\frac{1 - Cos\frac{2\pi t}{\tau_d}}{2}\right). \tag{77}$$

The Kirchhoff transform and non-dimensionalizing leads to,

$$\frac{\partial \Phi}{\partial T} = \frac{\tau_d}{\tau_s}\frac{\Phi}{R}\frac{\partial}{\partial R}\left(R\frac{\partial \Phi}{\partial R}\right) \qquad \rightarrow \tau_s = \frac{2\pi r_s^2 L_r T_s}{J_m D_o} \tag{78}$$

$$\frac{\partial \Phi}{\partial R}\bigg|_{R=1} = 1, \tag{79}$$

$$\frac{r_r}{r_s}\frac{\partial \Phi}{\partial R}\bigg|_{R=r_r/r_s} = \beta\, g(\Phi_l)\ (1 - Cos 2\pi T) \qquad \rightarrow \beta = \frac{E_p\, nVPD}{2J_m}. \tag{80}$$

As before, the geometrical scaling for the fluxes at the two domain boundaries only differ by $r_r/r_s$, with the result that the non-dimensionalization and re-scaling results in the exact same expressions for the boundary conditions as the big root case. However, here knowledge of the fine root density $L_r$ specifies a value for $r_s = \sqrt{1/(\pi L_r)}$ (Cowan, 1965) which together with an estimate of the typical fine root radius $r_r$ fixes the value of the ratio $r_r/r_s$. Another important difference is that the characteristic flux is now $J_m/(2\pi L_r T_s)$ which reflects the spreading of the inward steady flux over the surface area of all the roots in the root zone. While this does not effect the value of $\beta$ as demand is spread in the same proportion (the balance of the absolute fluxes in and out of the domain is invariant with rooting density), it does effect the characteristic time for the soil to respond to changes in the flux, leading to slower soil responses/greater damping of demand for high root density in thick root zones. The weakness of both approaches is the same. In both the big root and root density approaches, there is a hidden missing resistance between the arrival of the deep water flux at the lower planar boundary of the root zone, and the evenly spread inward flux that occurs across the entire outer surface area of the root and soil cylinder, regardless of how far any particular patch of surface on that cylinder is from the lower planar boundary. It is the hidden or cryptic nature of the neglected resistance that is of most concern, as SPAC modeling should as much as possible develop rational arguments for why some effects are retained in the model and others neglected. The approximation implicit in placing a source or fixed potential between two root cylinders may be defensible and not a bad approximation in some cases, but the artificiality should be clearly acknowledged (as done here), and not hidden behind the apparent realism of a cylindrical root and soil domain derived from rooting densities.

### 1.4 Surface Flux Equilibrium and ETRHEQ

Salvucci *et al.* (2013) and Rigden & Salvucci (2015) showed that at the daily time scale across a suite of Ameriflux sites surface conductance could be estimated by minimizing the vertical variance of $RH$ from weather data, an empirical finding termed ETRHEQ. McColl *et al.* (2019) provided a simple theory accounting for ETRHEQ through a box model of the Atmospheric Boundary layer, with the net radiation received at the lower surface partitioned into sensible and latent het fluxes. Unlike equilibrium evaporation (Raupach, 2001), the box into which heat and water vapor are transported from the surface is open at the top to the tropopause, resulting in higher $VPD$ and $E$ than in the equilibrium evaporation case. This system is then evolved to a steady-state in which the state of the atmosphere (represented by $RH$) has fully responded to the state of the land surface. At this quasi-equilibrium, latent and sensible heat fluxes become balanced in a manner that scales with the $RH$ of the atmosphere, termed Surface Flux Equilibrium. The relation of SFE to the equilibrium evaporation limit can be appreciated by considering the result for the evaporative fraction $EF$ at SFE,

$$EF = \frac{\lambda E}{R_n + G} = \frac{RH\ \epsilon}{RH\ \epsilon + 1}, \qquad \epsilon = \frac{c_p}{\lambda \frac{\partial \chi^*}{\partial T}} \tag{81}$$

Here $G$ or the flux to storage in the surface (plant and soil) is introduced into the energy balance for completeness, and the slope of the saturation curve with temperature is evaluated at 'screen level,' or two meters above the canopy. SFE takes the form of equilibrium evaporation of a wet surface into stagnant air (no lateral convection) corrected for unsaturated RH (Raupach 2001; Pieruschka *et al.*, 2010). The physical nature of this balance is more evident when expressed as the Bowen ratio of surface sensible to latent heat fluxes (McColl & Rigden 2020):

$$B = \frac{H}{\lambda E} = \frac{c_p}{\lambda \frac{\partial \chi^*}{\partial T} RH} \tag{82}$$

At SFE, sensible and latent heat fluxes are partitioned according to the amount of sensible and latent energy that air above the surface would absorb for a one degree increase in temperature at saturation, with the amount of latent then re-scaled to the observed de-saturated level of the atmosphere. Absorbed energy then simply flows into each channel of dissipation, sensible and latent, according to their relative capacity to absorb it. Notably all of the quantities on the right hand side of (81) can be evaluated from the state of the atmosphere, as in ETRHEQ, yet it would be a mistake to think of atmospheric RH (or 1-RH $\approx$ $VPD$) as controlling the surface flux balance. At SFE, the relative humidity and temperature profile of the ABL simply contains all the information about the moisture limitation in the land surface.

Table 2 Symbol definitions

| Quantity | Symbol | Units | Value |
|---|---|---|---|
| Aerodynamic conductance | $g_a$ | $m\ s^{-1}$ | 0.02 |
| Net Incident Energy | $R_n - G$ | $W\ m^{-2}$ | 1000 |
| Temperature 2 m above surface | $T$ | $K$ | 280, 300, 320 |
| Molar density of air | $C$ | $mol\ m^{-3}$ | 40.86 |
| Latent heat of vaporization | $\lambda$ | $J\ mol^{-1}$ | 44000 |
| Heat capacity of air | $C_p$ | $J\ mol^{-1}\ K^{-1}$ | 28.03 |
| gas constant | $R$ | $J\ mol^{-1}\ K^{-1}$ | 8.3145 |
| Atmospheric pressure | $p_{atm}$ | $Pa$ | $1.013 \times 10^5$ |

Following the ETRHEQ hypothesis, McColl and *et al.* (2019) find the surface conductance in molar units as,

$$g_s = \frac{g_a RH}{1 + \frac{\lambda C g_a \chi^*}{\epsilon(R_n - G)}(1 + \epsilon RH)} \frac{1}{1 - RH}, \tag{83}$$

where the last term is proportional to $1/VPD$. The plots in Figure 8 of the main text were generated from equations (81) and (83), using values taken from Table (2), based on Table 1 of McColl *et al.*(2019), with all temperature sensitive terms evaluated at the temperature 2 $m$ above the canopy, $T$. Only temperature was varied in Figure 8, as varying the other adjustable parameters in Table 2 had insignificant impacts on these normalized results. The slope of the saturation curve with temperature was calculated as,

$$\frac{\partial \chi^*}{\partial T} = \frac{\lambda \chi^*(T)}{RT^2}, \tag{84}$$

where the saturated mole fraction $\chi^*(T)$ was calculated according to Rockwell *et al.* (2014).

For the purposes of presentation and comparison to the empirical models, after calculating $E(RH, T)$ and $G_s(RH, T)$, values of $RH$ along the x axis were converted to $VPD$ through the relation,

$$VPD = (1 - RH)\,\chi^*(T) p_{atm}, \qquad \chi^*(T) \equiv \frac{p_v^*}{p_{atm}}. \tag{85}$$

The plots were then normalized as $VPD/VPD_{ref}$, $E/E_{ref}$, $g_s/g_{s,ref}$, with $E_{ref}$ and $g_{s,ref}$ evaluated at the reference $VPD_{ref} = 1\,kPa$.

### 1.5 Additional References (those not cited in the main text)


**Brodribb TJ, & Holbrook NM. 2003.** Stomatal closure during leaf dehydration, correlation with other leaf physiological traits. *Plant Physiology* 132: 2166–2173.

**Herkelrath WN, Miller EE, Gardner WR. 1977.** Water uptake by plants II: The root contact model. *Soil Science* 41: 1039–1043.

**McCully ME, McCully JSB, And Boyer ME. 1997.** The expansion of maize root-cap mucilage during hydration. 3. Changes in water potential and water content. *Physiologia Plantarum* 99: 169–177.

**Peavy, BA. 1996.** A heat transfer note on temperature dependent thermal conductivity. *J Thermal Insul. and Bldg. Envs.* 20:76-90.

**Philip, J. 1957.** The Physical Principles of soil water movement during the irrigation cycle. *Proceedings of 3rd International Commission on Irrigation and Drainage* 125–154.

**Pieruschka R, Huber G, Berry JA. 2010.** Control of transpiration by radiation. *Proc Natl Acad Sci USA* 107: 13372–13377.

**Scoffoni C, Albuquerque C, Brodersen CR, Townes S V., John GP, Bartlett MK, Buckley TN, McElrone AJ, Sack L. 2017.** Outside-xylem vulnerability, not Xylem embolism, controls leaf hydraulic decline during dehydration. *Plant Physiology* 173: 1197–1210.

**Shatil-Cohen A, Attia Z, Moshelion M. 2011.** Bundle-sheath cell regulation of xylem-mesophyll water transport via aquaporins under drought stress: A target of xylem-borne

ABA? *Plant Journal* 67: 72–80.
**Sperry JS, Adler F, Campbell G, Comstock J. 1998.** Limitation of plant water use by rhizosphere and xylem conductance: results from a model. *Plant, Cell and Environment* 21: 347–359.
**Zhang YJ, Rockwell FE, Graham AC, Alexander T, Holbrook NM. 2016.** Reversible leaf xylem collapse: A potential 'circuit breaker' against cavitation. *Plant Physiology* 172: 2261–2274.
