## Supplemental Notes S2 for "Emergent feedforward and isohydric responses to soil and atmospheric aridity: Insights from a time-dependent hydraulic model"

In[ ]:=
```
Remove["Global`*"]
```

New Phytologist Supporting Information
Article title: Emergent feedforward and isohydric responses to soil and atmospheric aridity: Insights from a time-dependent hydraulic model.
Author: Fulton E. Rockwell
Article acceptance data:
File: Notes S2. Computer code for model runs and figures.

#### Stomatal parameters

```
b = .23;(* Shape parameter: controls steepness of stomatal curve *)

ψhalf = -1.33; (* Water potential units: stomatal 50% closure MPa*)
τhalf = ψhalf * 10 227; (* matric potential units stomatal 50% closure cm *)

ℊhalf = -430.109/τhalf ;(* Potential flux units: stomatal 50% closure cm^2/day *)

-430.109/(10 227 ℊhalf) == ψhalf (* check units conversion logic *)
```

Out[ ]=

```
True
```

##### Stomatal conductance as f($\psi$): Fig. 1C

In[ ]:=
```
Plot[1/(1 + Exp[(ψhalf-x)/b]), {x, 0, -3}, PlotRange → Full,
 AxesStyle → Directive[Black, 28], PlotStyle → {Black, Thick}]
```

Out[ ]=

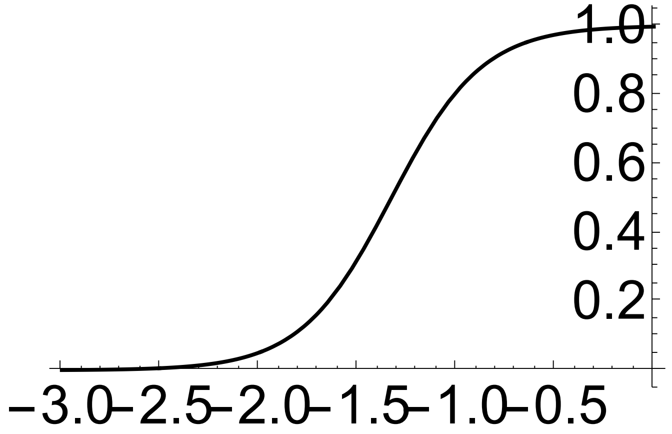

#### Stomatal conductance as f($\tau$)

In[ ]:=

$$\text{Plot}\left[\frac{1}{1+\text{Exp}\left[\frac{\tau\text{half}-x}{10\,227\,b}\right]},\ \{x,\ 0,\ -30\,000\},\ \text{PlotRange}\to\text{Full}\right]$$

Out[ ]=

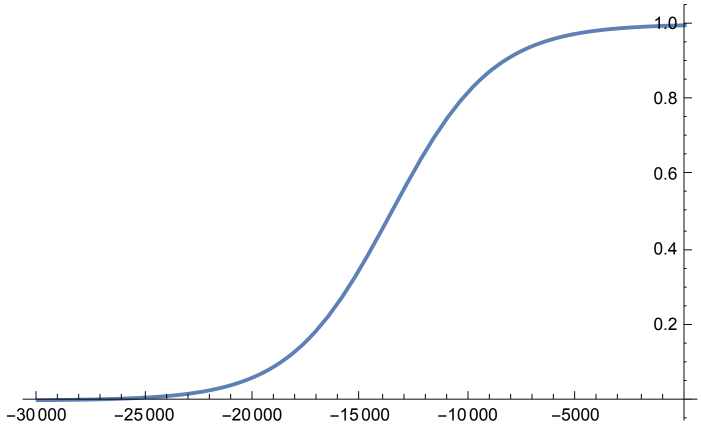


#### Stomatal conductance as f($\zeta$): Fig. 1D

In[ ]:=

$$\text{Plot}\left[\frac{1}{1+\text{Exp}\left[\frac{\frac{-430.109}{10\,227\,\zeta\text{half}}-\left(\frac{-430.109}{10\,227\,x}\right)}{b}\right]},\ \{x,\ 0,\ 1\},\ \text{PlotRange}\to\text{Full},\ \text{AxesStyle}\to\text{Directive}[\text{Black},\ 28],\ \text{PlotStyle}\to\{\text{Black},\ \text{Thick}\}\right]$$

Out[ ]=

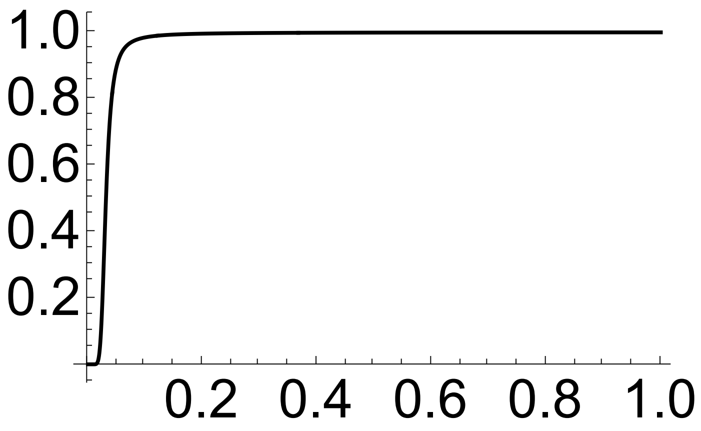

#### Diurnal forcing function: Fig. 1b

In[ ]:=
```
Plot[(1 - Cos[2 π t])/2, {t, 0, 7}, PlotRange → Full,
 AxesStyle → Directive[Black, 28], PlotStyle → {Black, Thick}]
```

Out[ ]=

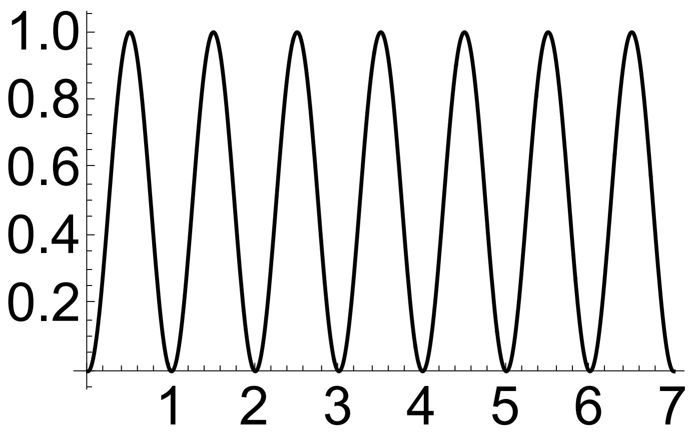


In[ ]:=
```
TotalV = NIntegrate[(1 - Cos[2 π t])/2, {t, 3, 7}]
```

Out[ ]=
```
2.
```

#### Model parameters:

In[ ]:=
```
H = 10;(* height in cm of root zone above water table: note that zeta=
 1 is for theta =0.25,
or 0.25 cm of water available to transpire per cm of depth.  *)
```

In[ ]:=
```
days = 7; (* length of time to run simulation *)
```

In[ ]:=
```
Q = .3;
```

In[ ]:=
```
q = .3; (* max flux cm/day from leaf *)
```

In[ ]:=
```
df = 2; (* divide potential flux to find max flux from soil*)
```

In[ ]:=
```
kp = q/(.5 × 10 227); (* k plant, units 1/day,
conductance of root to leaf path scaled so that the max
 flux gives a potential drop of .5 MPa in pressure units *)
```

```
ℊo = 1; (* Initial value of potential flux int he
 domain: may be modified by about 0.1 to imrpove stability of solution *)
```

```
vpd = 1; (* Nominal VPD levels: 1,1.25, 2, 2.5*)
```

#### Solve system: Main model

In[ ]:=

$$\text{sol2} = \text{NDSolve}\Big[\Big\{\text{D}[\zeta[t, x], t] == \frac{25.7\,\zeta[t, x]}{1}\,\text{D}[\zeta[t, x], x, x],$$

$$\zeta[0, x] == \zeta\text{o},\ \text{Derivative}[0, 1][\zeta][t, H] ==$$

$$\frac{q}{1 + \text{Exp}\left[\dfrac{\frac{-430.109}{10227\,\zeta\text{half}} - \left(\frac{-430.109}{10227\,\zeta[t,H]} + \frac{(\text{Derivative}[0,1][\zeta][t,H])}{10227\,\text{kp}}\right)}{b}\right]}\ \frac{(-\text{vpd} + \text{vpd}\,\text{Cos}[2\,\pi\,t])}{2},$$

$$\text{Derivative}[0, 1][\zeta][t, 0] == -\frac{Q}{\text{df}} * \left(1 - e^{-1000\,t}\right)\Big\},\ \zeta,\ \{t, 0, \text{days}\},$$

$$\{x, 0, H\},\ \text{Method} \to \text{"StiffnessSwitching"},\ \text{MaxSteps} \to \text{Infinity}\Big]$$

Out[ ]=

{{ζ → InterpolatingFunction[ Domain: {{0., 7.}, {0., 10.}} Output: scalar ]}}

#### Plot solution in zeta (cm/day): Supplemental Fig. S1

In[ ]:=

```
Plot3D[Evaluate[ζ[t, x] /. sol2], {t, 0, days},
 {x, 0, H}, PlotRange → All, AxesStyle → Directive[Black, 28]]
```

Out[ ]=

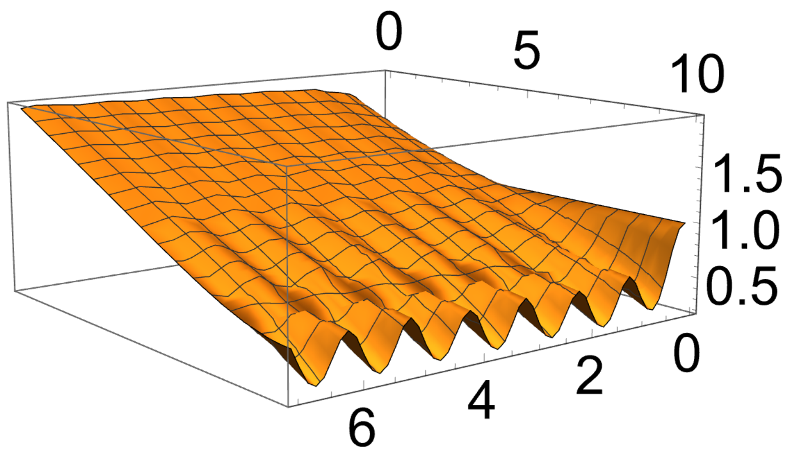

#### Plot solution in tau (cm)

In[ ]:=
```
Plot3D[Evaluate[-430.109/ξ[t, x] /. sol2], {t, 0, days},
 {x, 0, H}, PlotRange → All, AxesStyle → Directive[Black, 28]]
```

Out[ ]=

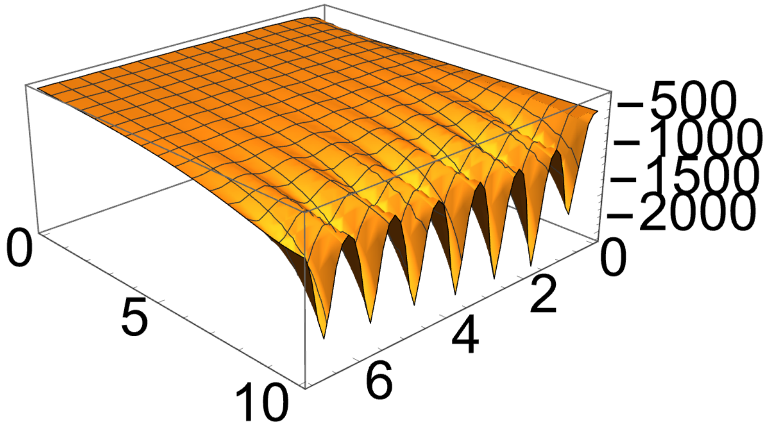


#### Plot solution in psi (MPa)

In[ ]:=
```
Plot3D[Evaluate[-430.109/(10 227 ξ[t, x]) /. sol2], {t, 0, days},
 {x, 0, H}, PlotRange → All, AxesStyle → Directive[Black, 28]]
```

Out[ ]=

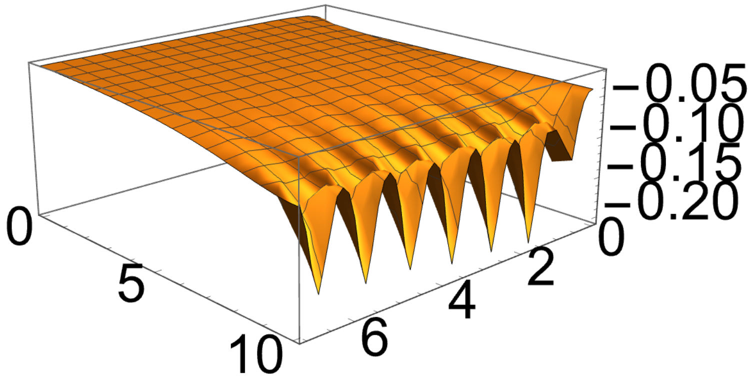

### Fluxes into and out of plant

#### Plot flux into root (cm/day): FIg. 4a and 5a, 6a

In[ ]:=
```
Plot[Evaluate[-Derivative [0, 1] [ξ][t, H] /. sol2],
 {t, 0, days}, PlotRange → {{0, days}, {0, .5}},
 AxesStyle → Directive[Black, 28], PlotStyle → {Blue, Thick}]
```

Out[ ]=

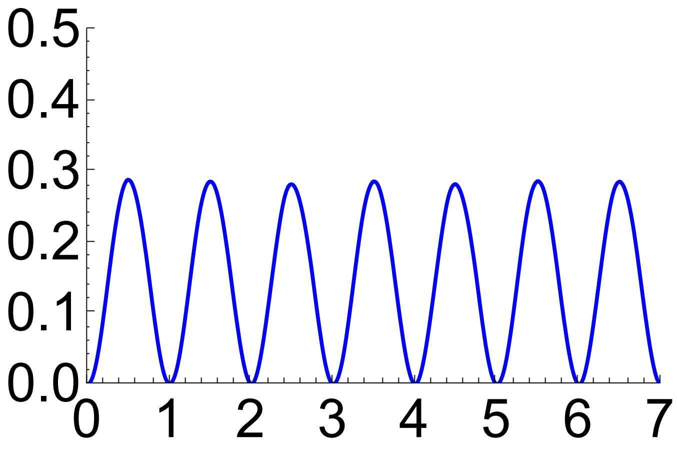


In[ ]:=
```
Evaluate[Derivative [0, 1] [ξ][4, H] /. sol2]
```

Out[ ]=

{0.00024194}

#### Find the average daily E (cm/day) into root over the experimental window Days 3,4,5,6

```
NIntegrate[-Derivative [0, 1] [ξ][t, H] /. sol2, {t, 3, days}] / (days - 3)
```

Out[ ]=

{0.145025}

#### Flux from leaf in cm/day

In[•]:=

```
Plot[
 Evaluate[ -q / (1 + Exp[ ( -430.109/(10 227 ζhalf) - ( -430.109/(10 227 ζ[t,H]) + (Derivative[0,1][ζ][t,H])/(10 227 kp) ) ) / b ]) * ( (-vpd + vpd Cos[2 π t]) / 2 ) /. sol2],
 {t, 0, days}, PlotRange → {{0, days}, {0, .35}},
 AxesStyle → Directive[Black, 28], PlotStyle → {Black, Thick}]
```

Out[•]=

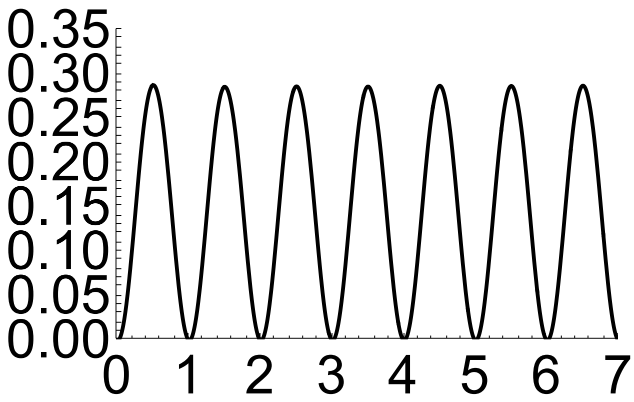


In[•]:=

```
TotalE =
 NIntegrate[ -q / (1 + Exp[ ( -430.109/(10 227 ζhalf) - ( -430.109/(10 227 ζ[t,H]) + (Derivative[0,1][ζ][t,H])/(10 227 kp) ) ) / b ]) * ( (-vpd + vpd Cos[2 π t]) / 2 ) /. sol2,
  {t, 3, days}]
```

Out[•]=

{0.582461}

```
DayE = TotalE / (days - 3)
```

Out[•]=

{0.145615}

#### Plot potential stomatal evaporation (gs)
#### [independent of diurnal driving force that ranges 0 to 1] Fig 4c and 5c, Fig 6b

In[ ]:=
```
pgs = Plot[Evaluate[1 / (1 + Exp[(-430.109/(10 227 ξhalf) - (-430.109/(10 227 ξ[t,H]) + (Derivative [0,1] [ξ][t,H])/(10 227 kp)))/b]) /. sol2],
  {t, 0, days}, PlotRange → {{0, days}, {0, 1}},
  AxesStyle → Directive[Black, 28], PlotStyle → {Blue, Thick}]
```

Out[ ]=

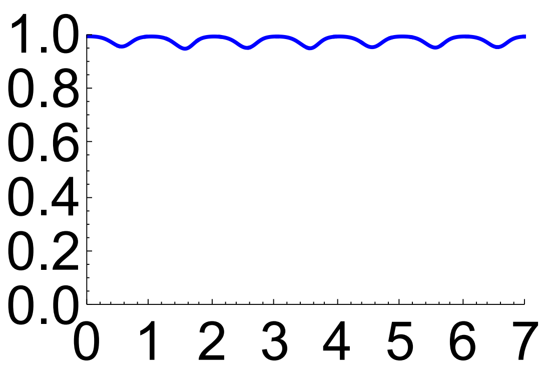


In[ ]:=
```
TotalA = NIntegrate[1 / (1 + Exp[(-430.109/(10 227 ξhalf) - (-430.109/(10 227 ξ[t,H]) + (Derivative [0,1] [ξ][t,H])/(10 227 kp)))/b]) /. sol2, {t, 3, days}]
```

Out[ ]=

```
{3.92184}
```

#### Leaf psi (MPa)

In[ ]:=
```
PL =
 Plot[Evaluate[-430.109/(10 227 ξ[t, H]) + (Derivative [0, 1] [ξ][t, H])/(10 227 kp) /. sol2], {t, 0, days},
  PlotRange → All, AxesStyle → Directive[Black, 28], PlotStyle → {Blue, Thick}]
```

Out[ ]=

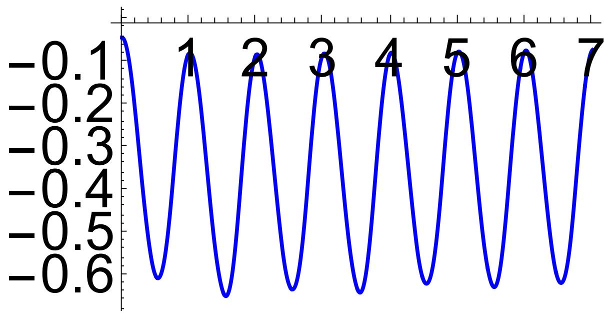

#### Root psi (MPa)

In[•]:=
```
PR = Plot[Evaluate[-430.109/(10 227 ξ[t, H]) /. sol2], {t, 0, days}, PlotRange → All,
  AxesStyle → Directive[Black, 28], PlotStyle → {Blue, Dashed}]
```

Out[•]=

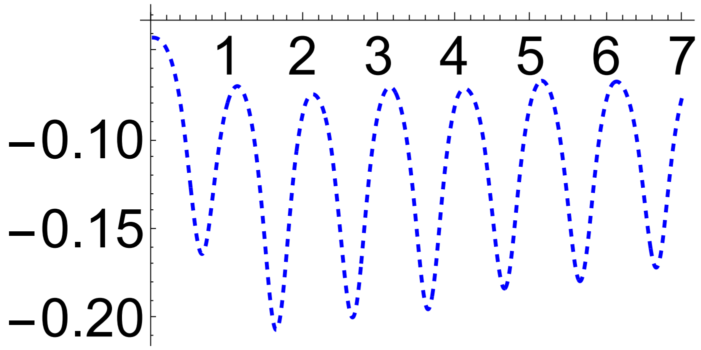


#### Root and leaf water potential: Fig 4b and 5b

In[•]:=
```
Show[PL, PR]
```

Out[•]=

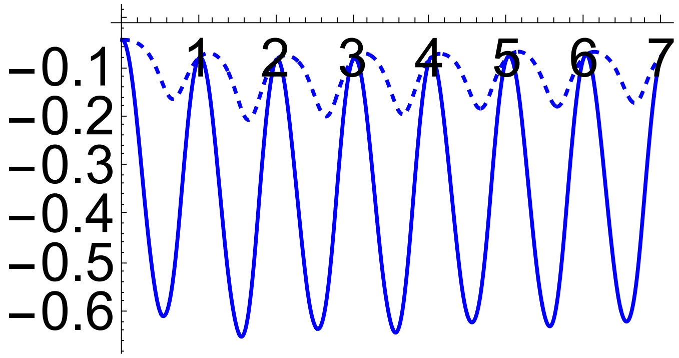


#### VPD response composite plot for nVPD v1=1, v2=1.25, v3=2, v4=2.5: Fig 7. Data for v# are from running the above model for each value of nVPD

In[•]:=
```
Remove["Global`*"]
```

In[•]:=
```
ListPlot[v1, AxesOrigin → {0, 0}, PlotRange → {{0, 3}, {0, 1}},
 AxesStyle → Directive[Black, 28], PlotStyle → {Cyan, Thick}]
```

Out[•]=

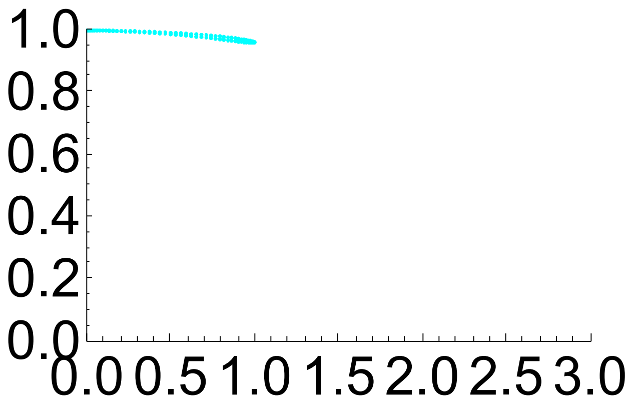


In[•]:=
```
ListPlot[v2, AxesOrigin → {0, 0}, PlotRange → {{0, 3}, {0, 1}},
 AxesStyle → Directive[Black, 28], PlotStyle → {Green, Thick}]
```

Out[•]=

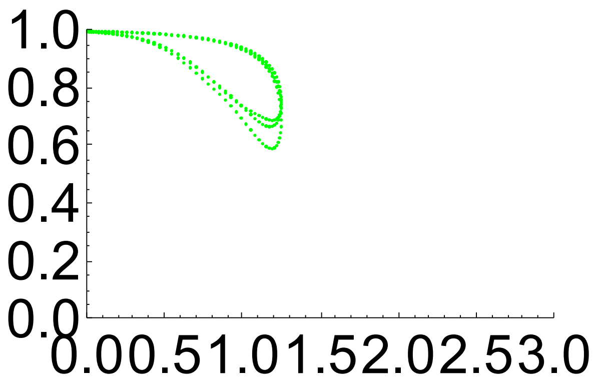


In[•]:=
```
ListPlot[v3, AxesOrigin → {0, 0}, PlotRange → {{0, 3}, {0, 1}},
 AxesStyle → Directive[Black, 28], PlotStyle → {Red, Thick}]
```

Out[•]=

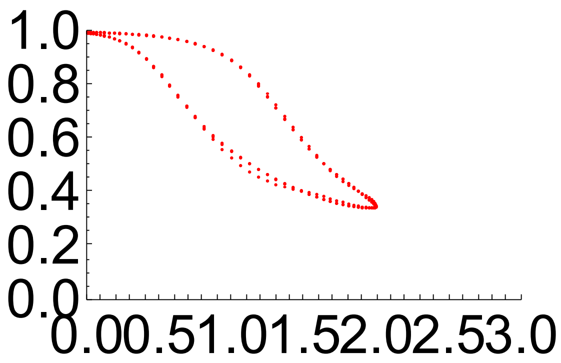

In[ ]:=
```
ListPlot[v4, AxesOrigin → {0, 0}, PlotRange → {{0, 3}, {0, 1}},
 AxesStyle → Directive[Black, 28], PlotStyle → {Red, Thick}]
```

Out[ ]=

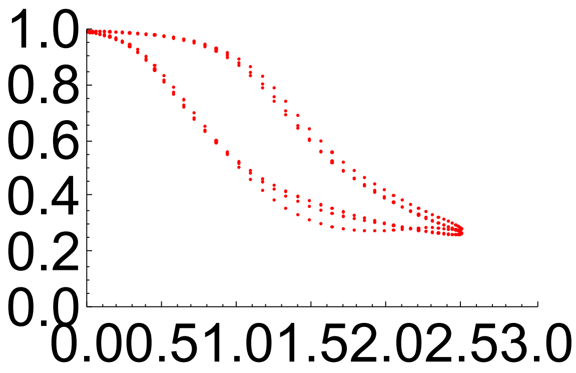


In[ ]:=
```
vall = Join[{v1, v2, v3, v4}];
```

In[ ]:=
```
ListPlot[vall, AxesOrigin → {0, 0},
 PlotRange → {{0, 2.5}, {0, 1}}, AxesStyle → Directive[Black, 28]]
```

Out[ ]=

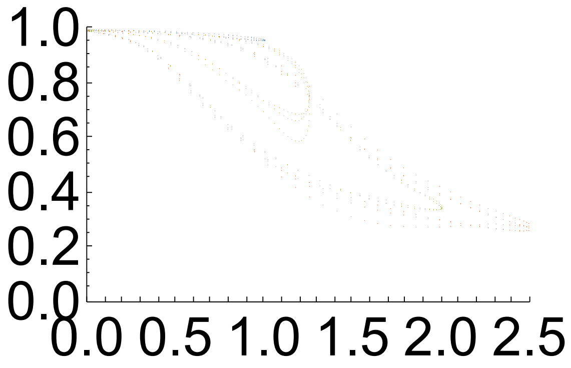


#### Generate model fit based on values  for which Oren's model is defined (nVPD>1)

In[ ]:=
```
data = Flatten[vall, 1];
```

In[ ]:=
```
datatofit = Select[data, #[[1]] ≥ 1 &];
```

In[ ]:=
```
vpd1 = Select[data, #[[1]] == 1 &];
```

In[ ]:=
```
AVG = Mean[vpd1]
```

Out[ ]=
```
{1., 0.79477}
```

In[ ]:=
```
Gref = AVG[[2]]
```

Out[ ]=
```
0.79477
```

In[•]:=
```
M = NonlinearModelFit[datatofit, a (1 - m Log[x]), {a, m}, x]
```
Out[•]=

FittedModel[ 0.783 (1 − 0.786 Log[x]) ]

In[•]:=
```
M["BestFitParameters"]
```
Out[•]=

{a → 0.782762, m → 0.785821}

In[•]:=
```
M["ParameterTable"]
```
Out[•]=

| | Estimate | Standard Error | t-Statistic | P-Value |
|---|---|---|---|---|
| a | 0.782762 | 0.0101269 | 77.2955 | $2.46567\times10^{-247}$ |
| m | 0.785821 | 0.0161448 | 48.6733 | $6.1573\times10^{-173}$ |

In[•]:=
```
M["AdjustedRSquared"]
```
Out[•]=

0.959935

In[•]:=
```
CI = M["MeanPredictionBands", ConfidenceLevel → .99]
```
Out[•]=

$$\{0.782762 - 0.61511\,\text{Log}[x] - 2.58781\,\sqrt{0.000102553 - 0.000322611\,\text{Log}[x] + 0.000349893\,\text{Log}[x]^2},\ 0.782762 - 0.61511\,\text{Log}[x] + 2.58781\,\sqrt{0.000102553 - 0.000322611\,\text{Log}[x] + 0.000349893\,\text{Log}[x]^2}\}$$

In[•]:=
```
CI[[1]]
```
Out[•]=

$$0.782762 - 0.61511\,\text{Log}[x] - 2.58781\,\sqrt{0.000102553 - 0.000322611\,\text{Log}[x] + 0.000349893\,\text{Log}[x]^2}$$

In[•]:=
```
p1 = ListPlot[datatofit, AxesOrigin → {1, 0}, PlotRange → {{1, 2.5}, {0, 1}},
  AxesStyle → Directive[Black, 28], PlotStyle → {Black, Thick}]
```
Out[•]=

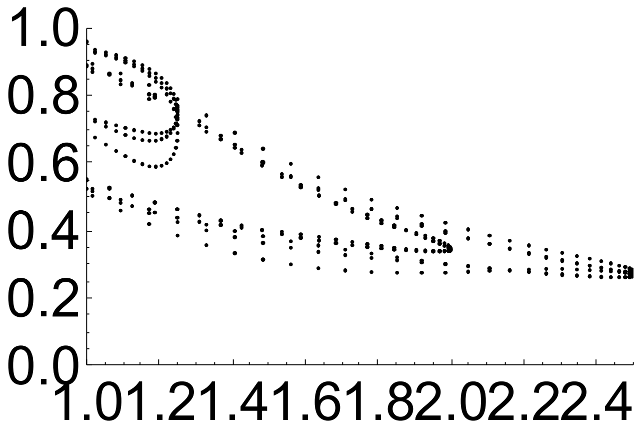

```
p2 = Plot[a (1 - m Log[x]) /. M["BestFitParameters"],
    {x, 1, 2.5}, AxesOrigin → {1, 0}, PlotRange → {{1, 2.5}, {0, 1}},
    AxesStyle → Directive[Black, 28], PlotStyle → {Black, Thick}];

p3 = Plot[CI⟦1⟧, {x, 1, 2.5}, AxesOrigin → {1, 0},
    PlotRange → {{1, 2.5}, {0, 1}}, PlotStyle → {Black, Dashed, Thick}];

p4 = Plot[CI⟦2⟧, {x, 1, 2.5}, AxesOrigin → {1, 0},
    PlotRange → {{1, 2.5}, {0, 1}}, PlotStyle → {Black, Dashed, Thick}];
```

In[ ]:=
```
Show[p1, p2, p3, p4]
```

Out[ ]=

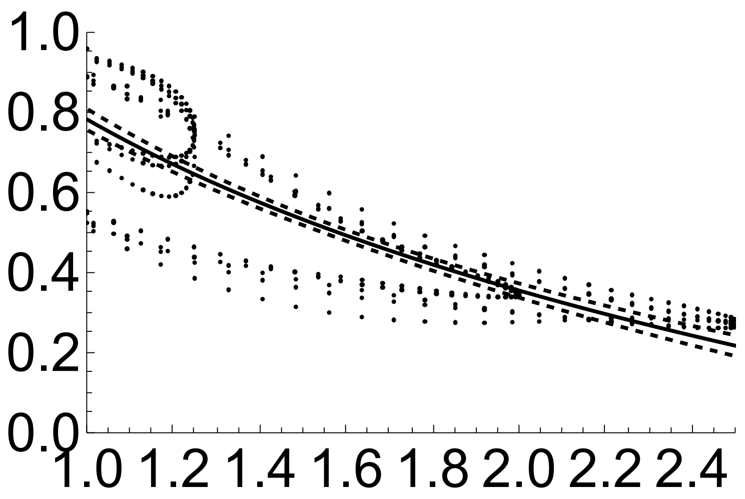


### Asymptotic soil steady-state soil fluxes for lower boundary condition at z=0.

#### Soil parameters: cm/day units

In[949]:=
```
k = ksat / (1 + (h / a) ^n); (* soil conductivity function, p 96  *)
```

In[950]:=
```
hwt = 0;
hmin = -∞;
a = -23.8; (* [cm], Gardner p 97  Chino clay *)
n = 2; (* Chino clay, Ibid *)
```

In[954]:=
```
ksat = 1.95 (* 1.95 cm/day  *)
```

Out[954]=
```
1.95
```

#### Solve Full Problem (retaining gravity and finite K and flux as psi-> 0), special case of n=2: (T is transpiration)

In[955]:=

$$\int_{-\infty}^{0} \frac{1}{1 + T / ks + (T / ks)\left(\frac{x}{b}\right)^2} \, dx$$

Out[955]=

$$\frac{b^2 \, ks \, \pi \sqrt{\frac{T}{b^2 (ks+T)}}}{2 \, T} \text{ if } \mathrm{Re}\left[\frac{b^2 (ks + T)}{T}\right] \geq 0 \;||\; \frac{b^2 (ks + T)}{T} \notin \mathbb{R}$$

In[956]:=

$$\frac{a^2 \, ksat \, \pi \sqrt{\frac{T}{a^2 (ksat+T)}}}{2 \, T} \; /. \; T \to .1$$

Out[956]=

```
161.011
```

In[957]:=

```
L = Range[1, 200, .1];
```

In[958]:=

```
TM = Table[0, Length[L]];
```

In[959]:=

$$\text{FindRoot}\left[L[[100]] == \frac{a^2 \, ksat \, \pi \sqrt{\frac{T}{a^2 (ksat+T)}}}{2 \, T}, \{T, 1\}\right]$$

Out[959]=

```
{T → 5.78383}
```

In[960]:=

$$\text{Do}\left[\text{TM}[[i]] = T \; /. \; \text{FindRoot}\left[L[[i]] == \frac{a^2 \, ksat \, \pi \sqrt{\frac{T}{a^2 (ksat+T)}}}{2 \, T}, \{T, 1\}\right], \{i, \text{Length}[L]\}\right]$$

In[961]:=

```
data = Transpose[{L, TM}];

Pfull = ListPlot[data, PlotStyle → Black];
```

#### For unsaturated source at head H

In[963]:=

```
TMH = Table[0, Length[L]];
```

In[964]:=

```
H = -10; (* matric potential  cm*)
```

In[965]:=

```
Do[
 TMH[[i]] =
  T /. FindRoot[L[[i]] == (a^2 ksat π Sqrt[T/(a^2 (ksat+T))])/(2 T) + (a^1 ksat ArcTan[(H Sqrt[T])/(a Sqrt[T+ksat])])/(Sqrt[T] Sqrt[ksat + T]), {T, 1}], {i,
  Length[L]}]
```

In[966]:=

```
dataH = Transpose[{L, TMH}];
PfullH = ListPlot[dataH, PlotStyle → Blue];
```

In[968]:=

```
TMHH = Table[0, Length[L]];
```

In[969]:=

```
H = -100; (* matric potential  cm*)
```

In[970]:=

```
Do[
 TMHH[[i]] =
  T /. FindRoot[L[[i]] == (a^2 ksat π Sqrt[T/(a^2 (ksat+T))])/(2 T) + (a^1 ksat ArcTan[(H Sqrt[T])/(a Sqrt[T+ksat])])/(Sqrt[T] Sqrt[ksat + T]), {T, 1}], {i,
  Length[L]}]
```

In[971]:=

```
dataHH = Transpose[{L, TMHH}];
PfullHH = ListPlot[dataHH, PlotStyle → Red];
```

In[973]:=

```
Show[Pfull, PfullH, PfullHH, PlotRange → {{10, 200}, {0, 1}},
 AxesOrigin → {10, 0}, AxesStyle → Directive[Black, 28], ImageSize → 600]
```

Out[973]=

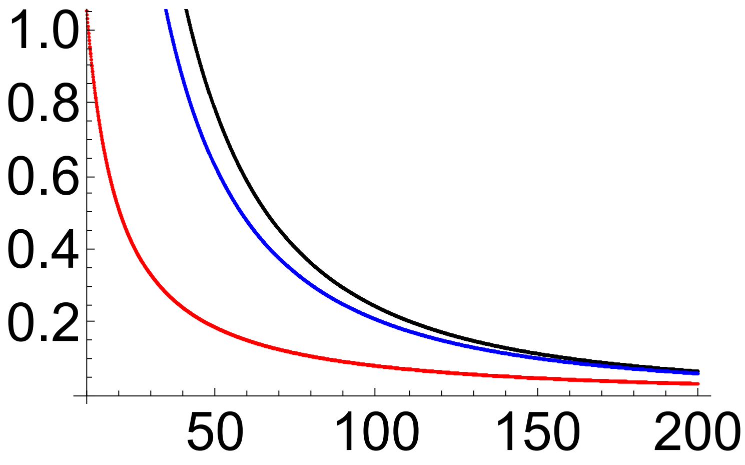

#### Asymptotic fluxes for a range of unsaturated sources

In[974]:=

```
h = {-1, -2, -4, -6, -10, -15, -20, -32, -50, -75,
   -100, -150, -200, -300, -400, -800, -1600, -5000, -10 000};
```

In[975]:=

```
TML = Table[0, Length[h]];
```

In[976]:=

```
l = 100; (* distance to source  cm*)
```

In[977]:=

Do[

$$\text{TML}[[i]] = T\ /.\ \text{FindRoot}\left[l == \frac{a^2\,\text{ksat}\,\pi\sqrt{\frac{T}{a^2\,(\text{ksat}+T)}}}{2\,T} + \frac{a^1\,\text{ksat}\,\text{ArcTan}\left[\frac{h[[i]]\,\sqrt{T}}{a\,\sqrt{T+\text{ksat}}}\right]}{\sqrt{T}\,\sqrt{\text{ksat}+T}},\ \{T, 1\}\right],$$

{i, Length[h]}]

In[978]:=

```
datah = Transpose[{-h, TML}];
```

In[979]:=

```
hfullLog = ListLogLinearPlot[datah, PlotStyle → Black,
  AxesStyle → Directive[Black, 28], ImageSize → 600]
```

Out[979]=

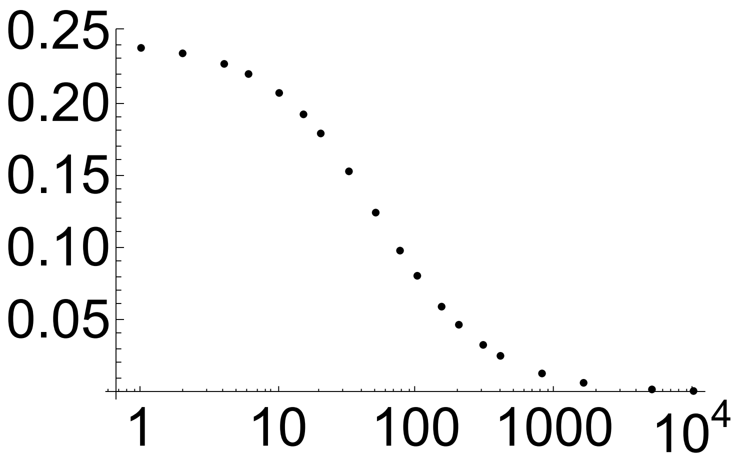


In[980]:=

```
P = 0.00009778` h ;(*convert h  in cm to P in MPa*)
```

In[981]:=

```
dataP = Transpose[{-P, TML}];
```

In[982]:=

```
PfullLog = ListLogLinearPlot[dataP, PlotStyle → Black,
  AxesStyle → Directive[Black, 28], ImageSize → 600]
```

Out[982]=

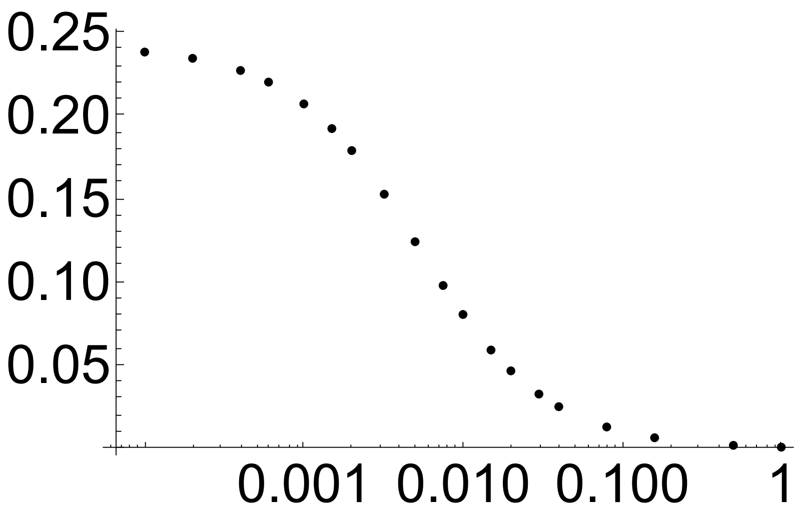


#### Finite fluxes approaching asymptote for a range of unsaturated sources -20 cm

In[983]:=

```
h = {-150, -200, -300, -400, -600, -800, -1200, -1600, -2500, -5000, -10000};
(* potential of absorbing *)
```

In[984]:=

```
y = -20; (* potential of source *)
```

In[985]:=

```
TML = Table[0, Length[h]];
```

In[986]:=

```
l = 100; (* distance to source  cm*)
```

In[987]:=

```
Do[
 TML[[i]] = T /. FindRoot[l == (a ksat (ArcTan[Sqrt[T] y / (a Sqrt[ksat+T])] - ArcTan[Sqrt[T] h[[i]] / (a Sqrt[ksat+T])])) / (Sqrt[T] Sqrt[ksat + T]), {T, 1}],
 {i, Length[h]}]
```

$$TML[\![i]\!] = T \;/.\; \text{FindRoot}\left[l == \frac{a\,ksat\left(\text{ArcTan}\left[\frac{\sqrt{T}\,y}{a\,\sqrt{ksat+T}}\right] - \text{ArcTan}\left[\frac{\sqrt{T}\,h[\![i]\!]}{a\,\sqrt{ksat+T}}\right]\right)}{\sqrt{T}\,\sqrt{ksat+T}}, \{T, 1\}\right]$$

In[988]:=

```
datah = Transpose[{-h, TML}];
```

In[989]:=

```
hfullLog = ListLogLinearPlot[datah, PlotStyle → Black,
  AxesStyle → Directive[Black, 28], ImageSize → 600]
```

Out[989]=

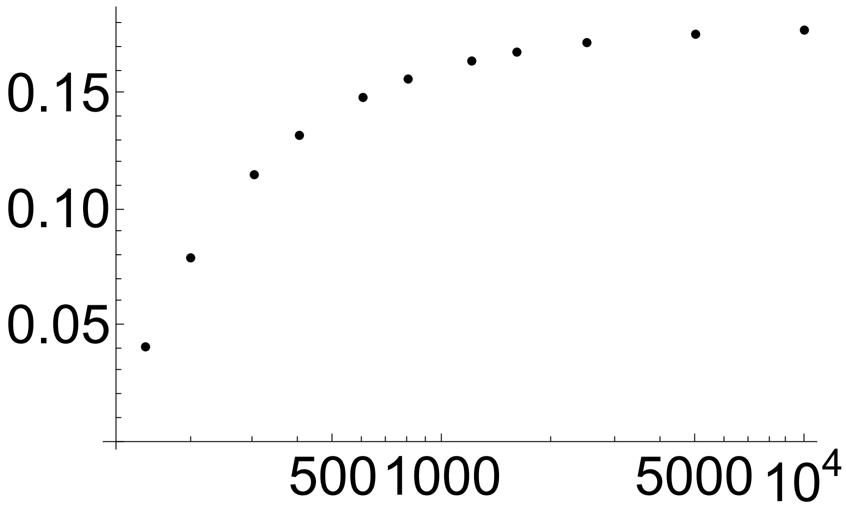


In[990]:=

```
P = 0.00009778` h ;(*convert h  in cm to P in MPa*)
```

In[991]:=

```
dataP = Transpose[{-P, TML}];
```

In[992]:=

```
PfullLog =
 ListLogLinearPlot[dataP, PlotRange → {{0.01, 1}, {0, .25}}, PlotStyle → Blue,
  AxesStyle → Directive[Black, 28], AxesOrigin → {0, 0}, ImageSize → 600]
```

Out[992]=

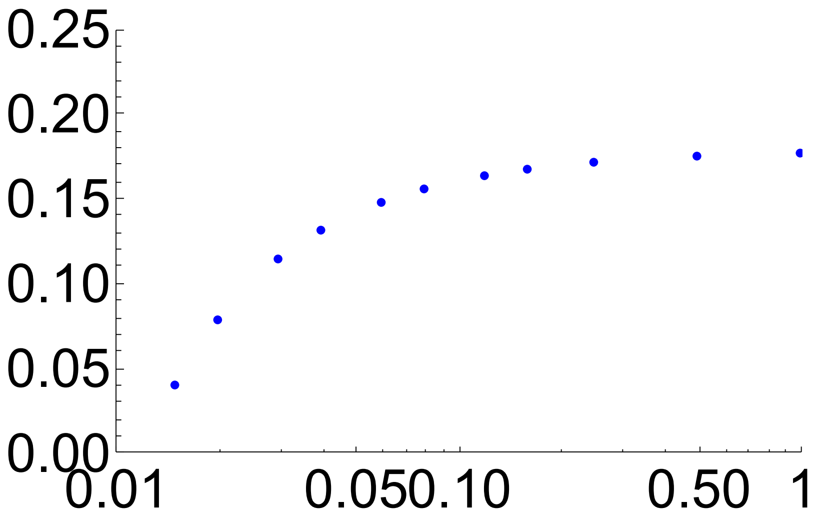

#### Finite fluxes approaching asymptote for an unsaturated source -100 cm

In[993]:=

```
h = { -300, -400, -600, -800, -1200, -1600, -2500, -5000, -10 000} ;
(* potential of absorbing *)
```

In[994]:=

```
y = -100; (* potential of source *)
```

In[995]:=

```
TML = Table[0, Length[h]];
```

In[996]:=

```
l = 100; (* distance to source  cm*)
```

In[997]:=

```
Do[
```

$$\text{TML}[\![i]\!] = T \;/.\; \text{FindRoot}\left[l == \frac{a\,\text{ksat}\left(\text{ArcTan}\left[\frac{\sqrt{T}\,y}{a\,\sqrt{\text{ksat}+T}}\right] - \text{ArcTan}\left[\frac{\sqrt{T}\,h[\![i]\!]}{a\,\sqrt{\text{ksat}+T}}\right]\right)}{\sqrt{T}\,\sqrt{\text{ksat}+T}}, \{T, 1\}\right],$$

```
 {i, Length[h]}]
```

In[998]:=

```
datah = Transpose[{-h, TML}];
```

In[999]:=

```
hfullLog = ListLogLinearPlot[datah, PlotStyle → Black,
  AxesStyle → Directive[Black, 28], ImageSize → 600]
```

Out[999]=

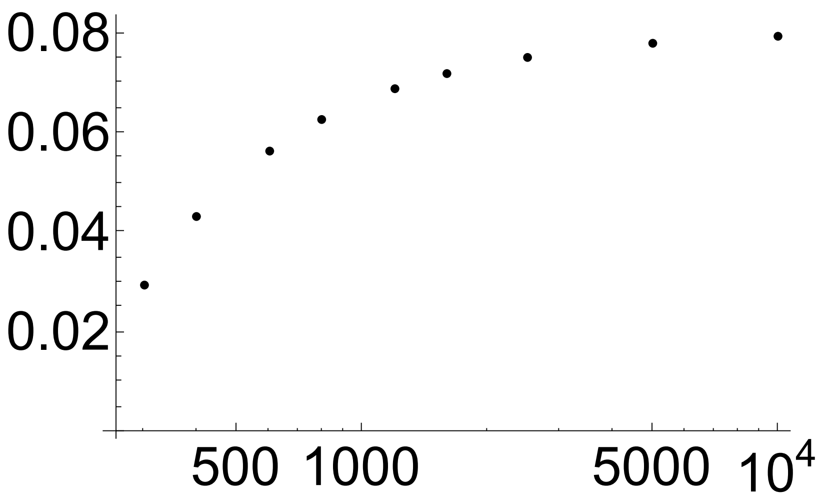


In[1000]:=

```
P = 0.00009778` h ;(*convert h  in cm to P in MPa*)
```

In[1001]:=

```
dataP = Transpose[{-P, TML}];
```

In[1002]:=

```
PfullLog =
 ListLogLinearPlot[dataP, PlotRange → {{0.01, 1}, {0, .25}}, PlotStyle → Red,
  AxesStyle → Directive[Black, 28], AxesOrigin → {0, 0}, ImageSize → 600]
```

Out[1002]=

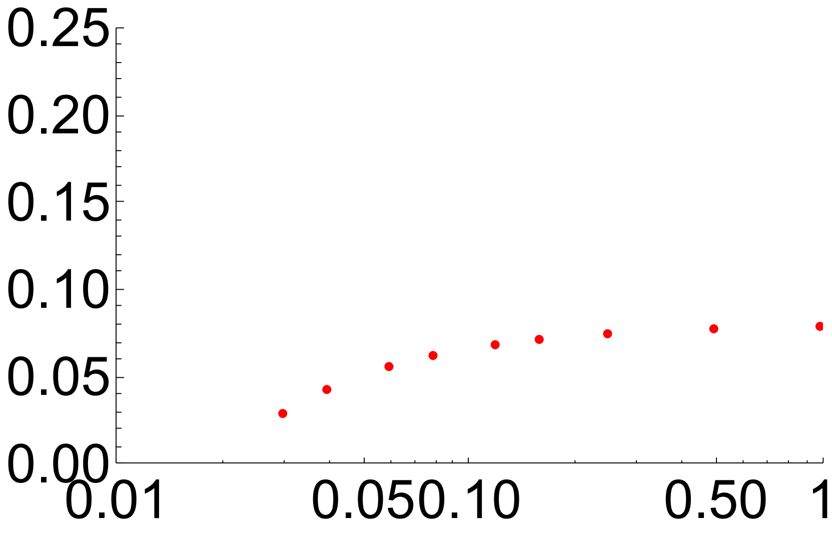


#### Finite fluxes approaching asymptote for an unsaturated source -10 cm

In[1003]:=

```
h = { -120, -150, -200, -300, -400, -600, -800, -1200,
   -1600, -2500, -5000, -10 000} ;(* potential of absorbing *)
```

In[1004]:=

```
y = -10; (* potential of source *)
```

In[1005]:=

```
TML = Table[0, Length[h]];
```

In[1006]:=

```
l = 100; (* distance to source  cm*)
```

In[1007]:=

```
Do[
 TML[[i]] = T /. FindRoot[l == (a ksat (ArcTan[(√T y)/(a √(ksat+T))] - ArcTan[(√T h[[i]])/(a √(ksat+T))]))/(√T √(ksat + T)), {T, 1}],
 {i, Length[h]}]
```

$$l = \frac{a\,ksat\left(\mathrm{ArcTan}\left[\frac{\sqrt{T}\,y}{a\,\sqrt{ksat+T}}\right]-\mathrm{ArcTan}\left[\frac{\sqrt{T}\,h[[i]]}{a\,\sqrt{ksat+T}}\right]\right)}{\sqrt{T}\,\sqrt{ksat+T}}$$

In[1008]:=

```
datah = Transpose[{-h, TML}];
```

In[1009]:=

```
hfullLog = ListLogLinearPlot[datah, PlotStyle → Black,
  AxesStyle → Directive[Black, 28], ImageSize → 600]
```

Out[1009]=

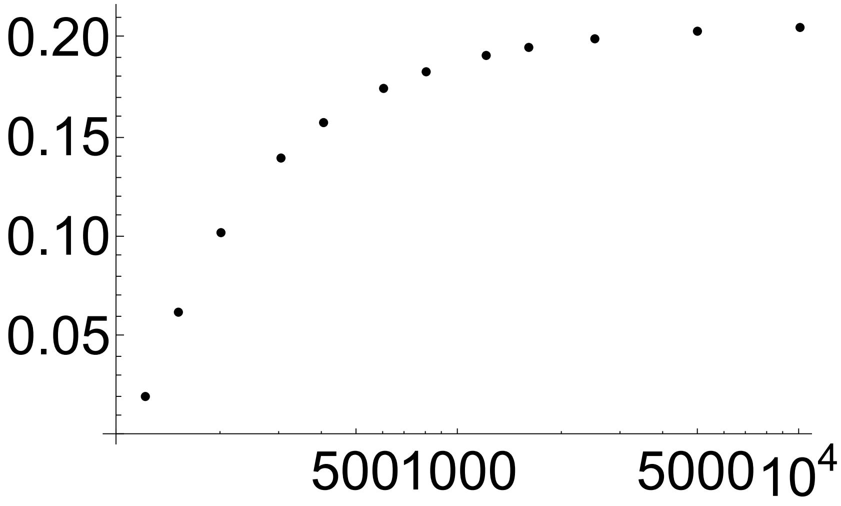


In[1010]:=

```
P = 0.00009778` h ;(*convert h  in cm to P in MPa*)
```

In[1011]:=

```
dataP = Transpose[{-P, TML}];
```

In[1012]:=

```
PfullLog =
 ListLogLinearPlot[dataP, PlotRange → {{0.01, 1}, {0, .25}}, PlotStyle → Black,
  AxesStyle → Directive[Black, 28], AxesOrigin → {0, 0}, ImageSize → 600]
```

Out[1012]=

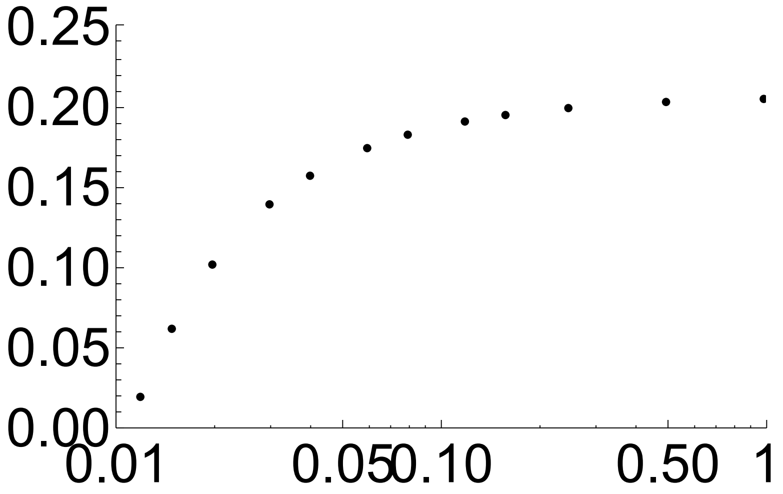

### Radial coordinates solution for soil and root cylinder domain

In[ ]:=
```
R = 10;(* height in cm of root zone above water table: note that zeta=
 1 is for theta =0.25,
or 0.25 cm of water available to transpire per cm of depth.  *)
```

In[ ]:=
```
r = 1;
```

In[ ]:=
```
days = 7; (* length of time to run simulation *)
```

In[ ]:=
```
q = .3; (* max flux cm/day from leaf *)
```

In[ ]:=
```
df = 2; (* divide potential flux to find max flux from soil*)
```

In[ ]:=
```
J = q / df;
```

In[ ]:=

$$kp = \frac{q}{.5 \times 10227};$$

```
(* k plant, units 1/day,
conductance of root to leaf path scaled so that the max
 flux gives a potential drop of .5 MPa in pressure units *)
```

In[ ]:=
```
ζo = 1.75;
```

In[ ]:=
```
vpd = 1;
```

#### Solve system

In[ ]:=

$$\text{sol2} = \text{NDSolve}\Big[\Big\{D[\zeta[t,x],t] == \frac{25.7\,\zeta[t,x]}{x} D[x\,D[\zeta[t,x],x],x],$$

$$\zeta[0,x] == \zeta o,\ \text{Derivative}[0,1][\zeta][t,r] ==$$

$$\frac{R}{\text{Pi}\ r}\ \frac{q}{1+\text{Exp}\left[\frac{\frac{-430.109}{10227\,\zeta\text{half}}-\left(\frac{-430.109}{10227\,\zeta[t,r]}-\frac{\text{Pi}\ r}{R}\ \frac{(\text{Derivative}[0,1][\zeta][t,r])}{10227\,kp}\right)}{b}\right]}\ \frac{(vpd - vpd\,\text{Cos}[2\pi t])}{2},$$

$$\text{Derivative}[0,1][\zeta][t,R] == \frac{J}{\text{Pi}} * \left(1 - e^{-1000\,t}\right)\Big\},\ \zeta,\ \{t, 0, \text{days}\},$$

$$\{x, r, R\},\ \text{Method} \to \text{"StiffnessSwitching"},\ \text{MaxSteps} \to \text{Infinity}\Big]$$

Out[ ]=

{{ζ → InterpolatingFunction[ Domain: {{0., 7.}, {1., 10.}} Output: scalar ]}}

In[ ]:=
```
sol1 = sol2
```

Out[ ]=

{{ζ → InterpolatingFunction[ Domain: {{0., 7.}, {1., 10.}} Output: scalar ]}}

#### Plot solution in zeta (cm/day)

In[∘]:=
```
Plot3D[Evaluate[ζ[t, x] /. sol2], {t, 0, days},
  {x, r, R}, PlotRange → All, AxesStyle → Directive[Black, 28]]
```

Out[∘]=

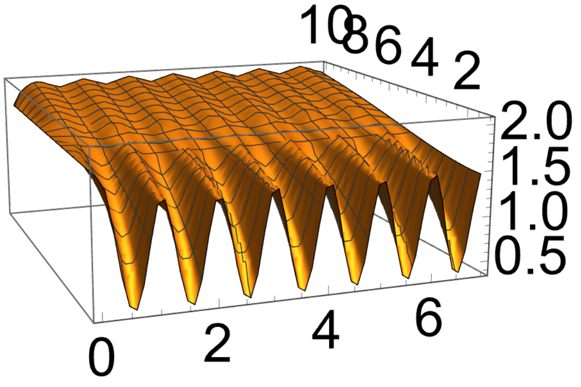


#### Plot solution in tau (cm)

In[∘]:=
```
Plot3D[Evaluate[-430.109/ζ[t, x] /. sol2], {t, 0, days},
  {x, r, R}, PlotRange → All, AxesStyle → Directive[Black, 28]]
```

Out[∘]=

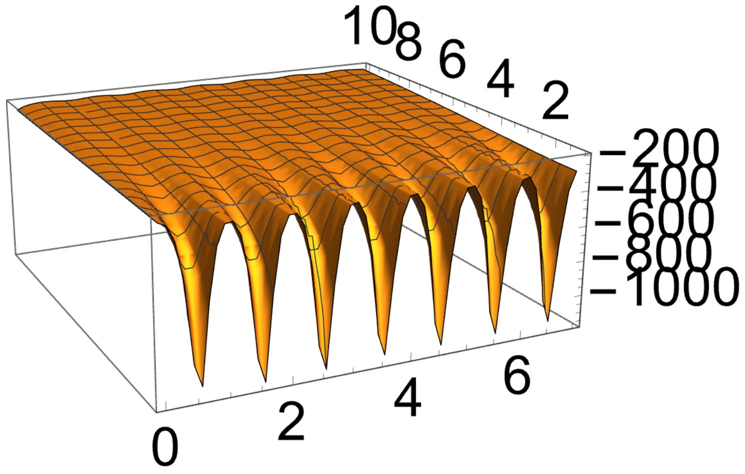

#### Plot solution in psi (MPa)

In[∘]:=
```
Plot3D[Evaluate[-430.109/(10227 ξ[t, x]) /. sol2], {t, 0, days},
  {x, r, R}, PlotRange → All, AxesStyle → Directive[Black, 28]]
```

Out[∘]=

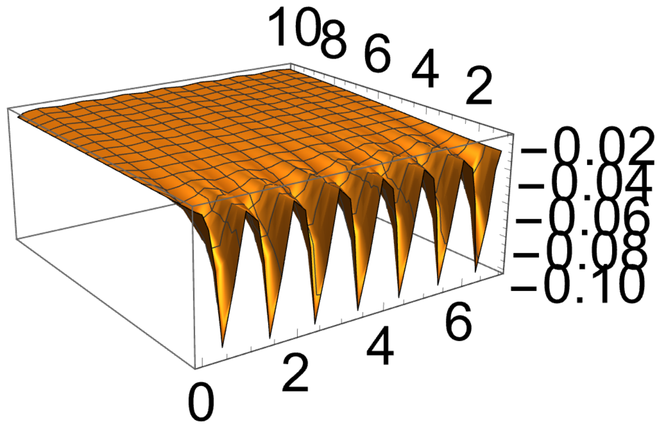


In[∘]:=
```
Plot[Evaluate[(Pi r)/R Derivative[0, 1][ξ][t, r] /. sol2], {t, 0, days},
  PlotRange → {{0, days}, {0, .5}}, AxesStyle → Directive[Black, 28],
  PlotStyle → {Black, Dashed, Thick}, ImageSize → 600]
```

Out[∘]=

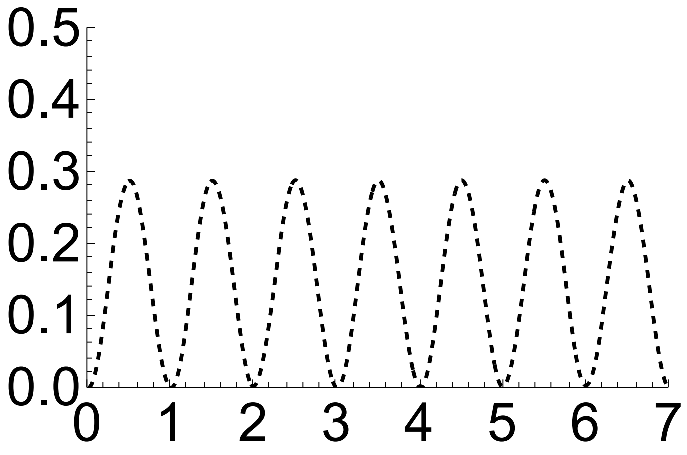

In[ ]:=
```
Plot[Evaluate[(Pi x)/R Derivative [0, 1] [ζ][4.5, x] /. sol2], {x, r, R},
  PlotRange → All, AxesStyle → Directive[Black, 28], PlotStyle → {Black, Thick}]
```

Out[ ]=

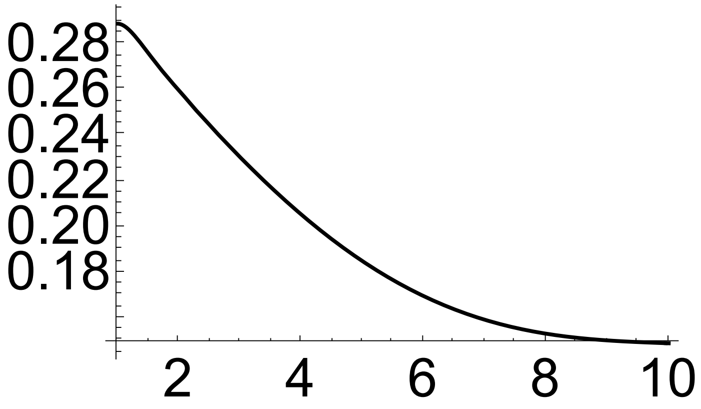


In[ ]:=
```
Plot[Evaluate[Pi/1 Derivative [0, 1] [ζ][t, R] /. sol2], {t, 0, days},
  PlotRange → All, AxesStyle → Directive[Black, 28], PlotStyle → {Black, Thick}]
```

Out[ ]=

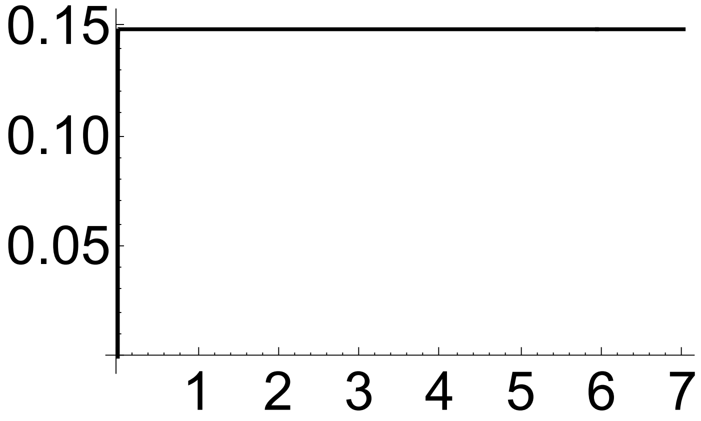

In[∘]:=
```
pl = Plot[Evaluate[-430.109/(10 227 ξ[t, r]) - (Pi r)/R (Derivative[0, 1] [ξ][t, r])/(10 227 kp) /. sol2],
  {t, 0, days}, PlotRange → All,
  AxesStyle → Directive[Black, 28], PlotStyle → {Black, Thick}]
```

Out[∘]=

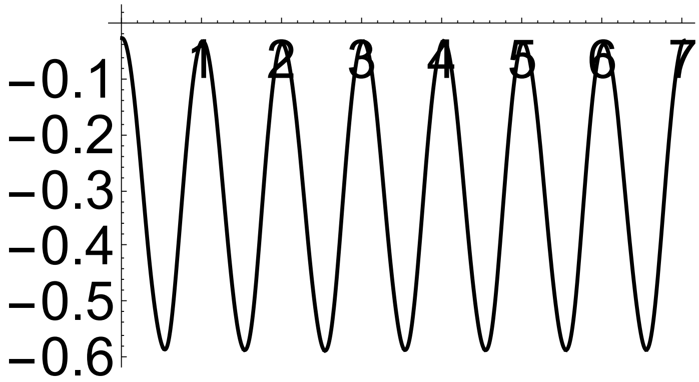


In[∘]:=
```
pr = Plot[Evaluate[-430.109/(10 227 ξ[t, r]) /. sol2], {t, 0, days}, PlotRange → All,
  AxesStyle → Directive[Black, 28], PlotStyle → {Black, Dashed, Thick}]
```

Out[∘]=

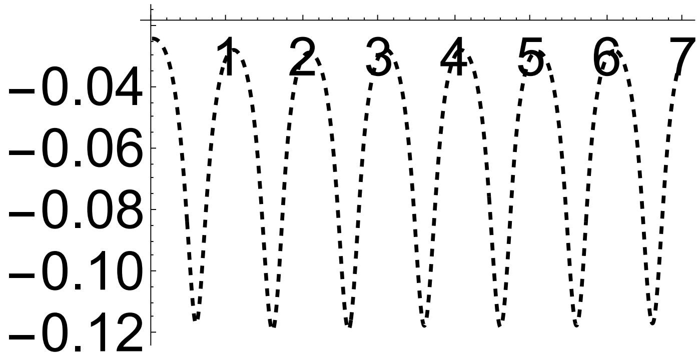

In[ ]:= Show[pl, pr, ImageSize → 600, AxesOrigin → {0, -1.6}]

Out[ ]=

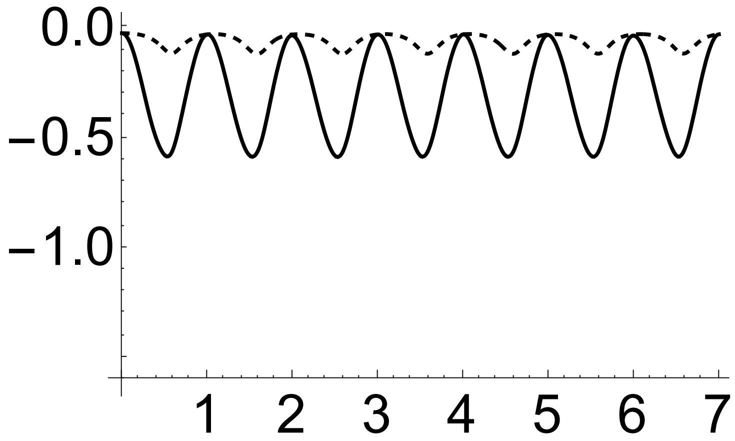


In[ ]:= Plot[Evaluate[$\frac{-430.109}{10\,227\,\xi[t, R]}$ /. sol2], {t, 0, days}, PlotRange → All, AxesStyle → Directive[Black, 28], PlotStyle → {Black, Thick}]

Out[ ]=

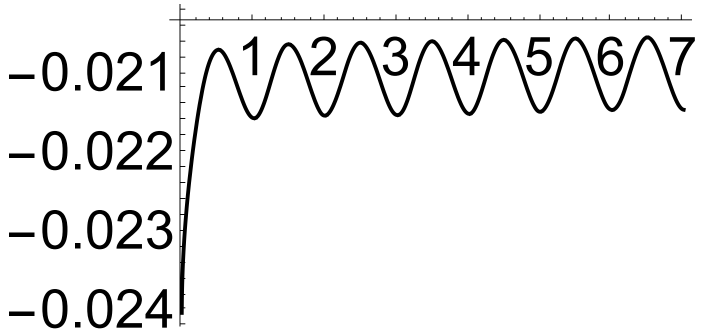

In[ ]:=
```
Plot[Evaluate[-430.109/(10 227 ξ[4.5, x]) /. sol2], {x, r, R},
  PlotRange → All, AxesStyle → Directive[Black, 28],
  PlotStyle → {Black, Dashed, Thick}, ImageSize → 600, AxesOrigin → {1, -1.2}]
```

Out[ ]=

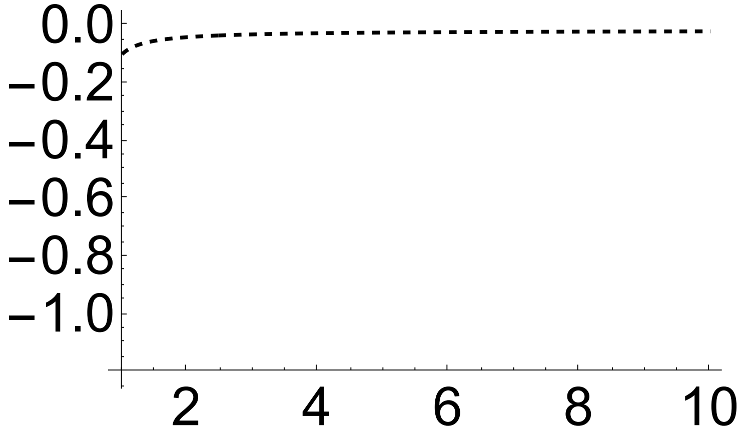
